# Chemical modification of proteins by insertion of synthetic peptides using tandem protein trans-splicing

**DOI:** 10.1101/2020.01.31.925495

**Authors:** K.K. Khoo, I. Galleano, F. Gasparri, R. Wieneke, H. Harms, M.H. Poulsen, H.C. Chua, M. Wulf, R. Tampé, S.A. Pless

## Abstract

Manipulation of proteins by chemical modification is a powerful way to decipher their function or harness that function for therapeutic purposes. Despite recent progress in ribosome-dependent and semi-synthetic chemical modifications, these techniques sometimes have limitations in the number and type of modifications that can be simultaneously introduced or their application in live eukaryotic cells. Here we present a new approach to incorporate single or multiple post-translational modifications or non-canonical amino acids into soluble and membrane proteins expressed in eukaryotic cells. We insert synthetic peptides into proteins of interest via tandem protein trans-splicing using two orthogonal split intein pairs and validate our approach by investigating different aspects of GFP, Na_V_1.5 and P2X2 receptor function. Because the approach can introduce virtually any chemical modification into both intracellular and extracellular regions of target proteins, we anticipate that it will overcome some of the drawbacks of other semi-synthetic or ribosome-dependent methods to engineer proteins.

## Introduction

Chemical or genetic engineering of proteins provides great potential to study protein function and pharmacology or to generate proteins with novel properties. However, despite recent technical achievements^1, 2^, the type of chemical modification that can be accomplished by genetic means (e.g. amber codon suppression) is limited to incorporation of non-canonical amino acids (ncAAs) due to the tolerance of the cell’s translational machinery. Additionally, insertion of multiple chemical modifications by genetic code expansion remains a challenge, particularly in eukaryotic cells. Semi-synthetic approaches offer an alternative means to manipulate proteins post-translationally, but these modifications have typically been performed *in vitro*^3–8^. We thus sought to complement these approaches with a method that could incorporate synthetic peptides carrying multiple post-translational modifications (PTMs) or ncAAs into both cytosolic and membrane proteins in live eukaryotic cells.

Split intein pairs comprise complementary N- and C-terminal intein fragments (Int^N^ and Int^C^) that assemble with extraordinary specificity and affinity to form an active intein. This assembly results in a spontaneous, essentially traceless splicing reaction that covalently links the two flanking protein segments through native chemical ligation^9^. The critical requirement for splicing to occur is typically the presence of a Cys, Ser or Thr side chain (depending on the split intein in question) in the +1 position of the extein (the sequence flanking the split intein) and multiple split inteins have recently been optimized for increased splicing efficiency^10–12^. The latter facilitates the simultaneous use of two orthogonal split inteins within the same peptide or protein, an approach termed tandem protein trans-splicing (tPTS). However, tPTS has largely been conducted *in vitro* or restricted to bacterial expression systems, cell lysates, nuclear extracts, or selection protocols^8, 13–15^. Indeed, most live cell applications of PTS utilize single split inteins for the purpose of N/C-terminal tagging^16–18^ or manipulating protein assembly/expression^19, 20^.

Here we employ tPTS using two orthogonal split intein pairs to insert synthetic peptides into proteins between two splice sites (A and B). This approach permits the introduction of a virtually limitless array of modifications, including PTMs, PTM mimics and ncAAs, into live eukaryotic cells and allows multiple modifications to be made simultaneously (Fig 1). We validate our approach by using tPTS to modify GFP, intracellular regions of the Na_V_1.5 channel, and the extracellular domain of the P2X2 receptor, allowing us to gain insight into the role of PTMs and PTM mimics in ion channel function and the importance of spatial positioning of charge in ligand sensitivity.

**Figure 1.**
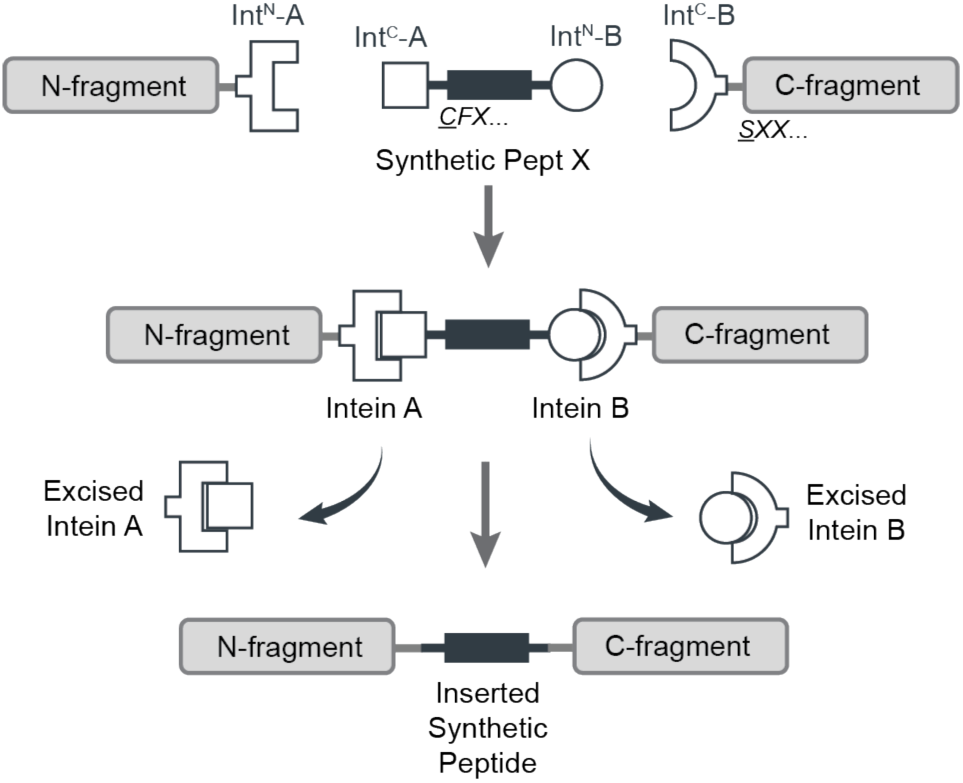
Schematic diagram of the tPTS strategy to incorporate recombinant and synthetic proteins. The chosen split inteins used in this study, inteins A (*Cfa*DnaE) and B (*Ssp*DnaB^M86^), are indicated by square and round symbols, respectively. Flanking +1, +2, +3 extein residues for each intein are indicated in italics at their respective positions in the top panel. The +1 extein residue (underlined) is a critical requirement for splicing to occur. *X* denotes that the type of residue at that position is not critical for splicing, although they might affect the kinetics or splicing efficiency.

## Results

### Strategy for post-translational incorporation of synthetic peptides

Our goal was to generate semi-synthetic proteins in live eukaryotic cells by post-translationally incorporating ncAAs or PTMs into a protein of interest. We achieved this by using two orthogonal split inteins (A and B) to insert a synthetic peptide carrying these modifications. We designed three fragments of the protein of interest (Fig 1), corresponding to N and C terminal fragments (N and C) and a shorter central fragment containing the desired modification (peptide X). Fragments N and C were heterologously expressed in the cell, while peptide X was generated synthetically and inserted into the cell via an appropriate technique (e.g. injection). To covalently assemble the three fragments, the highly efficient engineered derivative of the *Npu*DnaE split intein (termed *Cfa*DnaE)^11^ was employed as split intein A. The first 101 amino acids of its N-terminal (Int^N^-A) were expressed as a fusion construct at the C-terminus of protein fragment N. The corresponding C-terminal part (Int^C^-A), consisting of amino acids 102-137, were attached to the N-terminal end of peptide X. The optimized split intein *Ssp*DnaB^M86^ (ref. 10) was chosen as split intein B because it can be split highly asymmetrically and has previously been shown to be orthogonal to the *Npu*DnaE split intein^21^. Its N-terminal part (Int^N^-B), comprising only the first 11 amino acids, was added to the C-terminus of peptide X. The corresponding C-terminal part (Int^C^-B), consisting of amino acids 12-154, was expressed as a fusion construct at the N-terminus of protein fragment C (Fig 1).

### Replacing Na_v_1.5 inter-domain linkers with synthetic peptides

To demonstrate the feasibility of our approach, we chose the well-characterized cardiac voltage-gated sodium channel isoform Na_V_1.5, which is crucial for the initiation and propagation of the cardiac action potential^22^. This large 2016-amino acid protein comprises four homologous domains (DI-DIV) that are connected by intracellular linkers (Fig 2a). Dysfunction of Na_V_1.5 can arise from mutations, as well as dysregulated PTMs. For example, acetylation of K1479 and changes in phosphorylation of the linker between DIII-DIV have been shown to play a role in cardiac disease^23, 24^. However, interrogation of the role of PTMs has been hampered by the inability to express a homogenous population of channels containing a defined number of PTMs in living cells. Thus, although phosphorylation at Y1495 is known to affect channel function^25^ and phosphorylation can be prevented in a channel population by mutating Y1495 to phenylalanine, this population cannot be compared with one that is fully modified because the extent of phosphorylation cannot be controlled. Similarly, it is not known if there are synergistic effects with other PTMs in the vicinity, e.g. acetylation of K1479.

**Figure 2.**
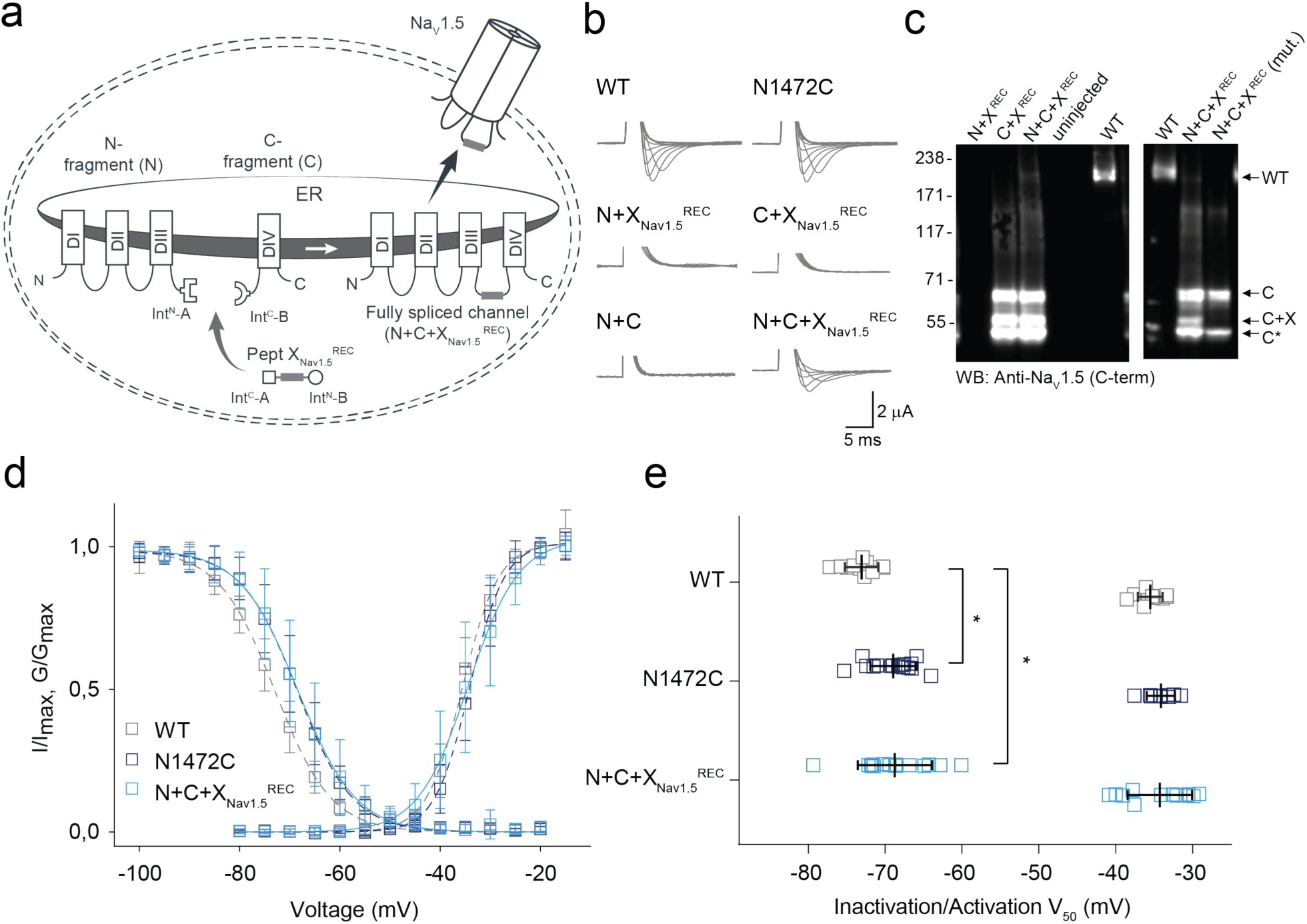
Insertion of recombinantly expressed peptides into the DIII-IV linker of Na_V_1.5. (a) Schematic presentation of the strategy to reconstruct full-length Na_V_1.5 from recombinantly expressed N-/C-terminal fragments (N and C) and a recombinantly expressed peptide corresponding to amino acids 1472 to 1502 of the Na_V_1.5 DIII-DIV linker (X) in *Xenopus laevis* oocytes. Inteins A (*Cfa*DnaE) and B (*Ssp*DnaB^M86^) are indicated by square and round symbols, respectively. Note that we cannot exclude the possibility that splicing takes place at a different subcellular location than depicted here. (b) Representative sodium currents in response to sodium channel activation protocol (see methods; only voltage steps from −50 mv to +10 mV in 10 mV steps are displayed), demonstrating expression of functional Na_V_1.5 only in the presence of all three components (N+C+X), along with WT and N1472C channels (the latter mutation was introduced to create an optimized splice site for intein A. (c) Immunoblots verifying the presence of fully spliced Na_V_1.5 only when all three components (N+C+X) were co-expressed (using antibody against Na_V_1.5 C-terminus). Na_V_1.5 band was not detected when one component was missing (left blot) or when non-splicing mutation (N+C+X mut.) was introduced to prevent splicing (right blot; see Fig S2). Black arrows indicate band positions of the respective constructs (Actual MW of constructs: WT, 227 kDa; C-construct, 79 kDa; C+X, 65 kDa; C-terminal cleavage product, C*, 58 kDa). Note that X and X^REC^ refer to X_Nav1.5_^REC^ in this panel. (d) Steady-state inactivation and conductance-voltage (G-V) relationships for functional constructs tested. (e) Comparison of values for half-maximal (in)activation (V_50_) (values are displayed as mean +/- SD; n = 12-16). Significant differences were determined by one-way ANOVA with a Tukey post-hoc test. *, p<0.03. Source data are provided as a Source Data file.

We tested the tPTS strategy by reconstituting full-length Na_V_1.5 from three recombinantly co-expressed channel fragments. To this end, we designed three different gene constructs: 1) an N-terminal construct (N) comprising amino acids 1-1471 of Na_v_1.5 (equivalent to DI-III) fused to the N-terminal part of *Cfa*DnaE (Int^N^-A); 2) a C-terminal construct (C) corresponding to the C-terminal part of *Ssp*DnaB^M86^ (Int^C^-B) linked to the C-terminal amino acids 1503-2016 of Na_v_1.5 (equivalent to DIV); and 3) a peptide X fragment (termed X_Nav1.5_^REC^) corresponding to the sequence to be replaced in the DIII-DIV linker of Na_V_1.5 (amino acids 1472 to 1502 of Na_V_1.5 but with N1472 mutated to Cys to enable splicing) flanked N- and C-terminally by Int^C^ of *Cfa*DnaE (Int^C^-A) and Int^N^ of *Ssp*DnaB^M86^ (Int^N^-B), respectively (Fig 2a). These constructs were transcribed into mRNA and injected into *Xenopus laevis* oocytes for recombinant expression. This approach is well-established for assessing ion channel function using electrophysiology and, conveniently, allows for direct delivery of mRNA and/or pepties into the cytosol using microinjection^26^.

As the peptide X fragment contained the N1472C mutation, we first compared the function of WT channels with N1472C mutant channels and the spliced product resulting from co-injection of N+C+X_Nav1.5_^REC^ (Fig 2b). As expected, injection of full-length WT and N1472C mRNA constructs resulted in robust channel expression, although the steady-state inactivation profile of N1472C was shifted slightly to more depolarized potentials, consistent with earlier reports suggesting for the N1472 locus to be potentially involved in cardiac disease^27^ (Fig 2b-e). Remarkably, co-injection of mRNA corresponding to N+C+X_Nav1.5_^REC^ (i.e. containing the N1472C mutation) resulted in full-length channels that showed robust current levels and were functionally indistinguishable from the full-length, recombinantly expressed channel construct also bearing the N1472C mutation (Fig 2d-e). Importantly, co-expression of only two of the three constructs (i.e. N+C, N+X_Nav1.5_^REC^ or C+X_Nav1.5_^REC^) did not result in any voltage-dependent sodium current (Fig 2b). Immunoblot analysis of co-expressed proteins also verified the presence of fully spliced Na_v_1.5 when X_Nav1.5_^REC^ was co-expressed with both N and C constructs, although the relative abundance of fully spliced product was low compared to unspliced or splicing side products (<2% estimated based on immunoblots of total cell lysates; Fig. 2c). Importantly, a band corresponding to fully spliced product was not detected when a splicing-incompetent mutation (+1 extein Ser to Ala mutation in the C construct at splice site B) was introduced (N+C+X_Nav1.5_ (mut.) in Fig 2c). Indeed, non-covalent assembly arising from split intein cleavage products and/or partially spliced channel fragments did not occur within the typical timeframe of our experiments (Fig S1). Together, these data demonstrate that tPTS can be used to assemble full-length Na_v_1.5 in live cells.

Having established that recombinant expression of N+C+X_Nav1.5_^REC^ can yield functional Na_v_1.5 channels, we next generated synthetic versions of peptide X (X_Nav1.5_^SYN^; see Fig S2 for synthesis strategy) for injection into cells expressing only the N and C fragments recombinantly (Fig 3a). Specifically, we synthesized X_Nav1.5_^SYN^ constructs that contained one of the following four variants: i) mutations K1479R and Y1495F (termed [NM]^Syn^) to prevent acetylation and phosphorylation, respectively; ii) a thio-acetylated Lys analog at position 1479 (tAcK1479) that mimics PTM but displays increased metabolic stability against sirtuins compared to regular acetylation^28^; iii) a phosphonylated Tyr analog at position 1495 (phY1495) that provides a non-hydrolysable phosphate mimic; or iv) both tAcK1479 and phY1495 to mimic a dual PTM scenario (Fig 3b). The N and C fragments were recombinantly expressed in oocytes for 24 h before injection of the synthetic X_Nav1.5_^SYN^ variants. Successful splicing of full-length Na_v_1.5 containing one of the four synthetic X_Nav1.5_^SYN^ variants was verified by immunoblotting and electrophysiology (Fig 3c,d). As before, the relative abundance of fully spliced product estimated from immunoblot analysis was low compared to the abundance of unspliced or splicing side products (< 1% in total cell lysates), but expression of robust voltage-gated sodium currents was achieved within 12 h of X_Nav1.5_^SYN^ variant injection. In fact, observed current levels at 24 h post peptide injection (i.e. 48 h after injection of N- and C-mRNA) were comparable to those observed 48 h after injection of WT mRNA (Fig 3e, Fig S3). To the best of our knowledge, this represents the first incorporation of a tAcK residue and the first insertion of two distinct PTM mimics in a full-length protein in eukaryotic cells. Functional analysis demonstrated that the voltage dependence of activation was not affected by any of the introduced PTM mimics or the conventional K1479R and Y1495F mutations. Conversely, insertion of phY1495, either alone or in combination with tAcK1479, induced a clearly discernable (15 mV) rightward-shift in the voltage-dependence of fast inactivation (Fig 3f). These data are consistent with earlier reports suggesting that acetylation of K1479 primarily affects current density^24^ whereas phosphorylation of Y1495 affects fast inactivation properties^25^.

**Figure 3.**
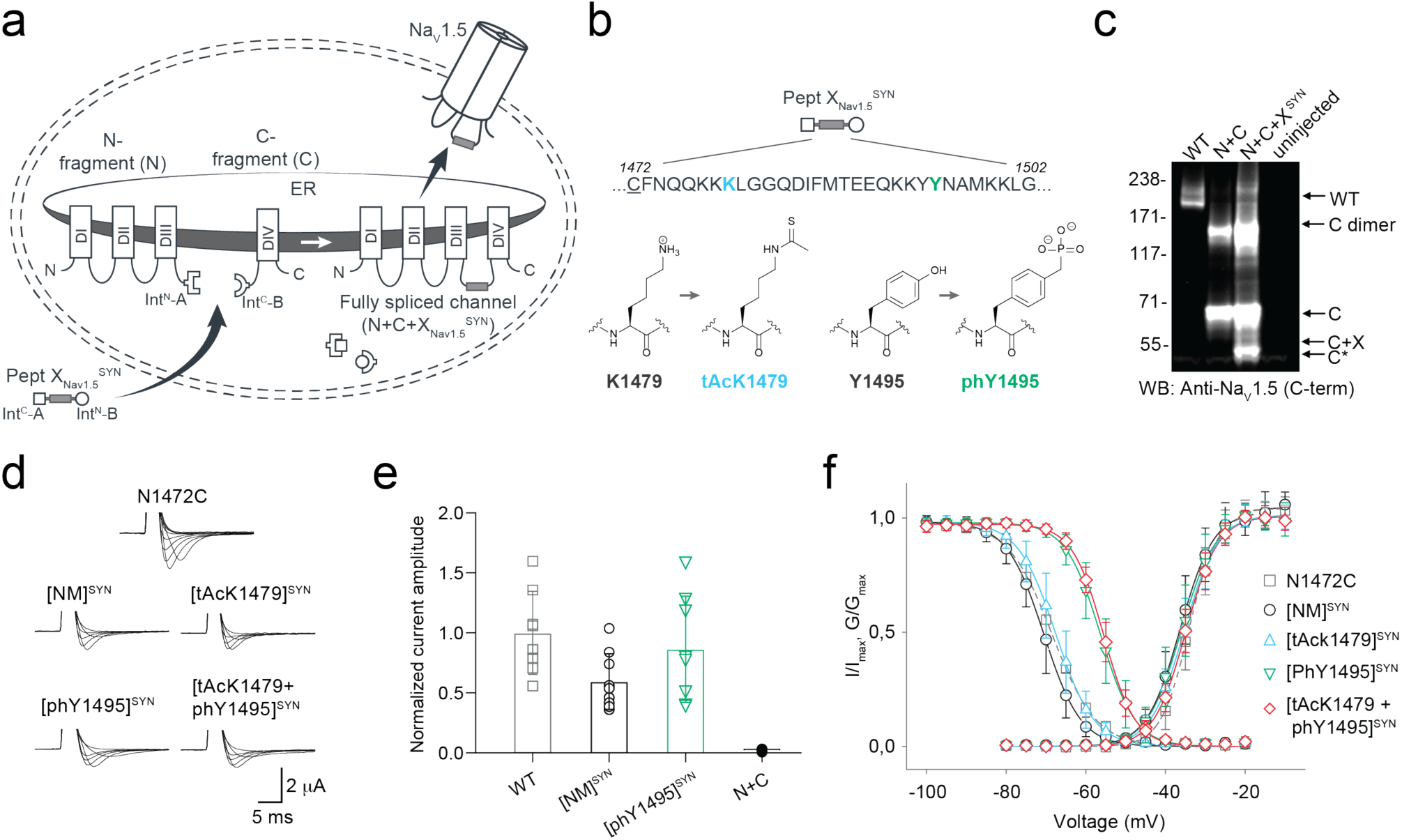
Insertion of synthetic peptides into Na_V_1.5. (a) Schematic presentation of the strategy to reconstruct full-length Na_V_1.5 from recombinantly expressed N-/C-terminal fragments (N and C) and a synthetic peptide (X_Nav1.5_ ^SYN^) in *Xenopus laevis* oocytes. Inteins A (*Cfa*DnaE) and B (*Ssp*DnaB^M86^) indicated by square and round symbols, respectively. Note that we cannot exclude the possibility that splicing takes place at a different subcellular location than depicted here. (b) Sequence of peptide X_Nav1.5_ ^SYN^ corresponding to the amino acids replaced in the Na_V_1.5 DIII-DIV linker and chemical structures of native amino acids and PTMs (tAcK/ phY) incorporated via chemical synthesis of peptide X_Nav1.5_ ^SYN^. The underlined N1472C mutation was introduced to optimize splicing (see also Fig S1). (c) Immunoblot verifying the presence of fully spliced Na_V_1.5 only when peptide X_Nav1.5_ was co-injected with N and C. Black arrows indicate band positions of the respective constructs (Actual MW of constructs: WT, 227 kDa; C-construct, 79 kDa; C+X, 65 kDa; C-terminal cleavage product, C*, 58 kDa). Band at ∼150 kDa is possibly a dimer or aggregate product of the C construct. (d) Representative sodium currents (see methods; only voltage steps from −50 mv to +10 mV in 10 mV steps are displayed), demonstrating expression of functional Na_V_1.5 when *Xenopus laevis* oocytes expressing N and C constructs were injected with synthetic peptides containing non-modifiable side chains in positions 1479 and 1495 (K1479R and Y1495F, NM), tAcK1479 or phY1495 or both PTMs together. (e) Average current amplitudes recorded at −35 mV from oocytes expressing N and C constructs and injected with synthetic peptide variant NM or phY1495 depicted as a bar plot (mean +/- SD; n = 6-9) Current amplitudes were normalized to the mean current measured from oocytes expressing the full-length WT construct. To ensure adequate control of voltage clamp, [Na^+^] in the extracellular recording solution was reduced (see Fig S3 for more details) (f) Steady-state inactivation and conductance-voltage (G-V) relationships for PTM-modified/non-modified constructs (values displayed as mean +/- SD; n = 10-21). Source data are provided as a Source Data file.

To further validate our approach and demonstrate its suitability for other target sequences, we applied the same strategy to the intracellular linker connecting DI and II of Na_V_1.5. Similar to the DIII-IV linker, mutations or aberrant PTMs in this region of Na_V_1.5 have been implicated in cardiac disease^23, 29^. Using appropriate N and C constructs, together with both recombinantly expressed and synthetic versions of a peptide X_Nav1.5_ variant corresponding to amino acids 505 to 527 of Na_V_1.5, we demonstrated that tPTS can be used to probe the function of different intracellular regions of Na_V_1.5 in *Xenopus* oocytes. Specifically, we found that neither methylation of R513 (meR513), nor phosphonylation of S516 (phS516), nor their combined presence^30^, affected activation or inactivation of Na_V_1.5 (Figs S4 and S5).

### Semi-synthesis of GFP in HEK cells

The above data showed that tPTS could be used to insert synthetic peptides into large membrane proteins in live eukaryotic cells, but it was important to demonstrate delivery into mammalian cells, which can be more challenging. To demonstrate the feasibility of this approach in mammalian cells, we split GFP into three fragments (analogous to our approach with Na_v_1.5 described above): 1) an N-terminal construct (N-GFP) corresponding to amino acids 1-64 of GFP, fused to Int^N^ of *Cfa*DnaE; 2) a C-terminal construct (C-GFP) corresponding to Int^C^ of *Ssp*DnaB^M86^ linked to amino acids 86-238 of GFP and 3) a peptide X fragment (X_GFP_^REC^) corresponding to amino acids 65 to 85 and flanked by Int^C^ of *Cfa*DnaE at the N-terminus and by Int^N^ of *Ssp*DnaB^M86^ at the C-terminus (Fig 4a). The constructs were co-expressed in different combinations in human embryonic kidney (HEK) cells, which expressed functional GFP only when all three constructs (N-GFP+C-GFP+X_GFP_^REC^) were transfected, albeit with low yields (∼4%, as estimated by fluorescence-activated cell sorting (FACS), Fig S6). No GFP fluorescence was detected with the co-expression of any two constructs or when cells were transfected with constructs containing splicing-incompetent mutations (C65A at +1 extein X_GFP_^REC^ (splice site A) or S85A at +1 extein C-GFP (splice site B); Fig 4b).

**Figure 4:**
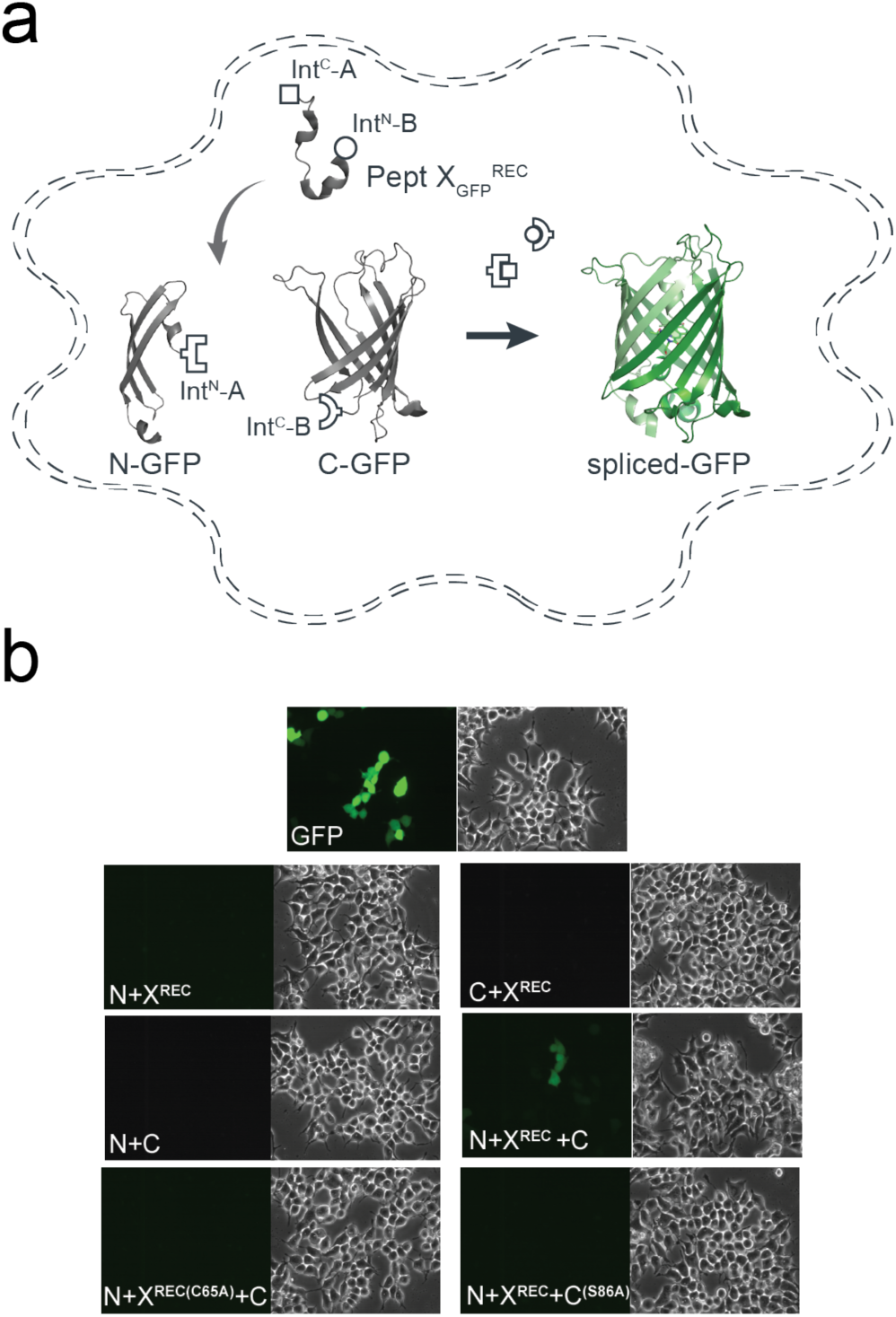
Insertion of a recombinantly expressed peptide into GFP expressed in mammalian cells. (a) Schematic presentation of the strategy applied to reconstitute GFP from recombinantly expressed N-/C- terminal fragments and recombinantly expressed peptide X_GFP_ corresponding to amino acids 65-85 of GFP in HEK293 cells. Inteins A (*Cfa*DnaE) and B (*Ssp*DnaB^M86^) are indicated by square and round symbols, respectively. (b) Bright field (right panels) and fluorescence (left panels) images of HEK293 cells expressing the indicated constructs. GFP fluorescence was only detected when all three constructs (N+X+C) were co-transfected. GFP fluorescence was not detected when one of the three constructs was absent or when +1 extein residues of each split intein is mutated to alanine (C65A for intein A and S86A for intein B) to prevent splicing.

We subsequently sought to generate a semi-synthetic GFP by delivering synthetic peptide X_GFP_^SYN^ variants into HEK cells that recombinantly expressed N- and C-terminal fragments of GFP (Fig 5a). We achieved delivery of synthetic peptides using the transient cell permeabilization method known as cell squeezing, which involves rapid viscoelastic deformation^31^. Although yields were low (approx. 1%), GFP fluorescence was detected only in N-GFP- and C-GFP-transfected cells that had been squeezed in the presence of peptide X_GFP_^SYN^ (Fig 5b). The approach further allowed us to incorporate the ncAA 3-nitro-tyrosine at position 66 of GFP to replace the tyrosine that is involved in chromophore formation. This modification resulted in a blue-shift in the spectral properties of GFP and confirmed the utility of tPTS for creating semi-synthetic variants in mammalian cells (Fig S7).

**Figure 5.**
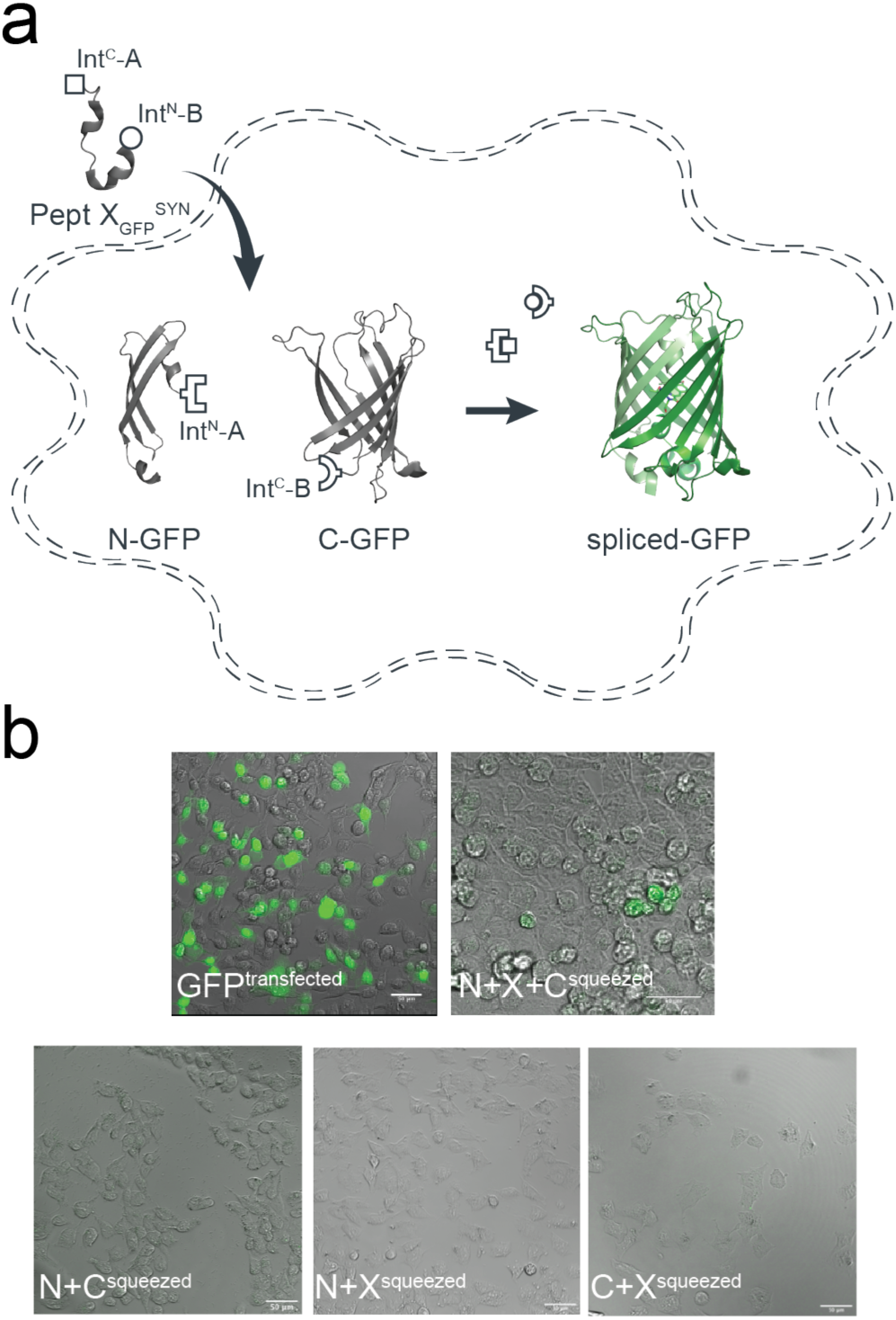
Insertion of synthetic peptides into GFP. (a) Schematic of strategy applied to reconstitute GFP from recombinantly expressed N-/C-terminal fragments and a synthetic peptide X_GFP_ in HEK293 cells. (b) Overlay of brightfield and fluorescence images taken of HEK293 cells transfected with only WT GFP, or N and C fragments and squeezed in the presence or absence of peptide X_GFP_.

### Insertion of ncAAs into the P2X2 receptor extracellular domain

While standard PTS has been employed to splice numerous cytosolic proteins and peptides, extracellular targets are more challenging and have been rarely investigated using PTS^17^. We sought to test whether tPTS could be used to insert synthetic peptides into an extracellular protein domain. We chose the P2X2 receptor (P2X2R), a trimeric ATP-gated ion channel whose extracellular domain binds ATP released during synaptic transmission^32^.

While the location of the ATP-binding site in the extracellular domain is undisputed, the details of how conserved basic side chains coordinate the phosphate tail of ATP remain unclear^33^. However, ribosome-based non-sense suppression approaches, using e.g. ncAA analogs of lysine, have failed at position K71 in the P2X2R, likely due to nonspecific incorporation of endogenous amino acids (Fig S8). We therefore used tPTS to test whether the charge position of K71 is crucial for ATP recognition.

As an initial proof-of-concept of splicing within an extracellular domain of a membrane receptor, we used standard PTS to independently assess splicing at either side of K71 in P2X2R: S54, which was mutated to Cys to improve splicing (splice site A), and S76 (splice site B). Again, we chose *Xenopus leavis* oocytes as an expression system, as they allow facile peptide delivery. For splice site A, the N-terminal fragment contained amino acids 1-53 of P2X2 linked to Int^N^ of *Cfa*DnaE. However, the C-terminal construct contained a *faux* transmembrane domain (amino acids 1-74 from ASIC1a) followed by Int^C^ of *Cfa*DnaE and the C-terminal receptor fragment of P2X2 (amino acids 54-472) (Fig S9a). Introduction of the *faux* transmembrane segment was necessary to enforce the correct topology of the resulting construct. To demonstrate that successful splicing is necessary for assembly of full-length receptors, we also generated a version of the C-terminal construct containing the C54A mutation, which effectively removes the required +1 Cys side chain and renders the construct splicing incompetent (Fig S9b). Expression of the individual constructs alone (N or C) in *Xenopus* oocytes did not result in functional receptors, whereas co-expression of N+C (but not N+C (C54A)) resulted in receptors with WT-like ATP sensitivity (Fig S9c,d). Confirmation of correct splicing was provided by immunoblots showing that bands corresponding to WT P2X2 only occurred in the presence of N+C, but not any of the control constructs (Fig S9e). Of note, biochemical analysis confirmed that splicing was highly efficient, with near complete conversion of the N and C fragments to full-length receptors.

We proceeded to test splice site B, employing an analogous approach to that implemented for splice site A (Fig S10a-b), except we used the *Ssp*DnaB^M86^ split intein in this case together with amino acids 1-75 of P2X2 (N-terminal fragment) and amino acids 76-472 of P2X2 plus a *faux* transmembrane segment (C-terminal fragment). Similar to the results obtained at splice site A, full-length P2X2 receptors with WT-like ATP sensitivity were only present upon co-expression of N+C, but not N or C alone (Fig S10c-e). Although lower than in the case of splice site A, the splicing was still efficient, with over half of the N and C fragments being converted into full-length receptors (Fig S10d). In contrast to splice site A, co-expression of the N and splicing-deficient C (S76A) constructs resulted in small ATP-gated currents in response to high concentrations of ATP (> 1 mM) after long incubation times (> 48 h). Of note, the prevention of splicing favors side reactions, which will result in the accumulation of cleavage products. It is thus possible that the non-covalent assembly of the N and C cleavage products results in a receptor population with impaired function, as evident from the drastically reduced ATP sensitivity we observed (Fig S10e). However, this result likely overstates the likelihood of cleavage products occurring compared to when splicing-competent split inteins are used. Overall, these data confirm that splicing can be achieved within an extracellular domain of a membrane receptor.

In order to insert a peptide fragment into the extracellular domain of P2X2 using tPTS, we used an analogous approach to that described for Na_v_1.5 and GFP to generate three constructs: 1) an N-terminal construct (N) corresponding to amino acids 1-53 of P2X2 fused to Int^N^ of *Cfa*DnaE; 2) a C-terminal construct (C) containing a *faux* transmembrane domain linked to Int^C^ of *Ssp*DnaB^M86^ and amino acids 76-472 of P2X2; and 3) a peptide X fragment (termed X_P2X2_^REC^) containing amino acids 54 to 75 of P2X2 flanked N- and C-terminally by Int^C^ of *Cfa*DnaE and Int^N^ of *Ssp*DnaB^M86^, respectively (Fig 6a). To optimize splicing efficiency, we additionally tested a C-terminal construct with a cleavable *faux* transmembrane domain, which comprised an IgK N-term signal sequence and a signal peptidase cleavage site inserted between the *faux* transmembrane segment and Int^C^ of *Ssp*DnaB^M86^. The resultant current amplitudes confirmed superior performance compared to the non-cleavable *faux* domain (Fig S11), therefore further experiments proceeded with this optimized C-terminal construct. Following expression in *Xenopus laevis* oocytes, splicing of full-length, ATP-gated receptors was only apparent when all three fragments (N, C, and X_P2X2_^REC^) were present, and represented an estimated 7% of the total products/reactants detected by immunoblotting (Fig 6b,c). Importantly, introduction of the S76A mutation at the +1 extein position of the C construct did not result in detectable currents. Further, introduction of the K71Q mutation-bearing peptide X_P2X2_^REC^ into the spliced receptors shifted the ATP concentration-response curve to the right to a similar degree as the conventional K71Q mutant (Fig 6d). Together, these data demonstrate successful and splicing-dependent assembly of functional P2X2 receptors upon co-expression of N, C, and X_P2X2_^REC^ constructs.

**Figure 6.**
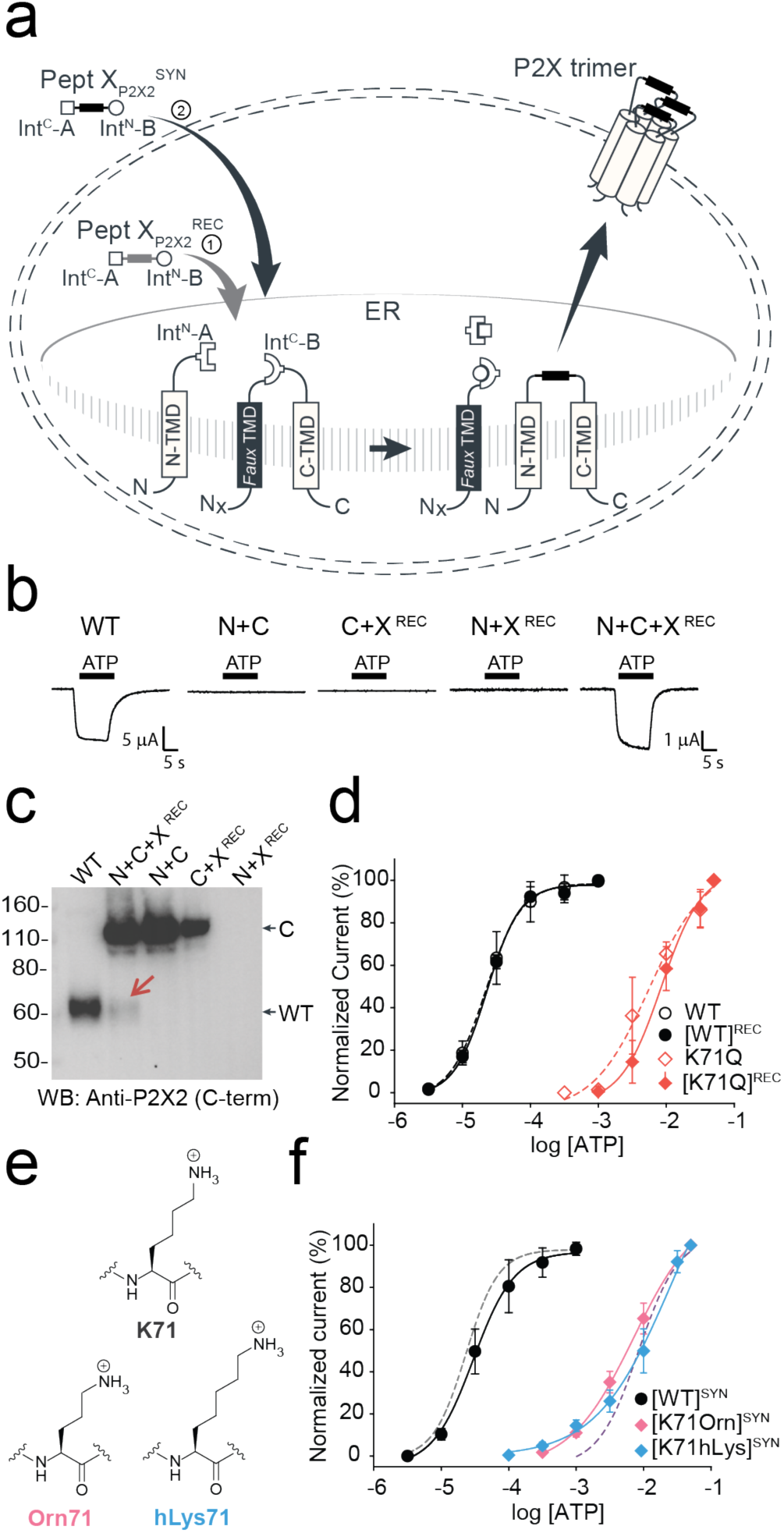
Insertion of recombinant and synthetic peptides into the P2X2R extracellular domain. (a) Schematic presentation of the strategy to reconstruct full-length P2X2Rs from recombinantly expressed N-/C-terminal fragments (N and C) and a recombinant (strategy 1, Pept X_P2X2_^REC^) or synthetic peptide (strategy 2, Pept X_P2X2_^SYN^) into the P2X2R extracellular domain in *Xenopus laevis* oocytes. Note that a *faux* transmembrane helix (*faux* TMD) was engineered into the C-terminal fragment to maintain its native membrane topology (see also Fig S9). Inteins A (*Cfa*DnaE) and B (*Ssp*DnaB^M86^) indicated by square and round symbols, respectively. Peptide X was designed to include a C-terminal ER-targeting KDEL signal sequence, which is excised during the splicing process. However, we cannot exclude the possibility that splicing takes place at a different subcellular location than depicted here. (b) Representative current traces obtained by application of 30 µM ATP, verifying functional expression only when all three components were recombinantly co-expressed (strategy 1 in (a)). (c) Immunoblot of surface-purified proteins. Black arrows on the right indicate band positions of the respective constructs (Actual MW of constructs: WT, 53 kDa; C-construct, 97 kDa). Red arrow indicates the band of the spliced full-length receptor. (d) ATP concentration-response curves (CRCs) for P2X2 WT (black) and K71Q (red) reconstituted from the three co-expressed constructs. Dashed lines represent the CRCs for the respective full-length proteins. (e) Structures of inserted side chains at position 71 (Lys, Orn, hLys) incorporated via synthetic peptide X (strategy 2 in (a)). (f) ATP CRCs indicate successful incorporation of synthetic WT peptide X_P2X2_ (black) with wild-type like ATP-sensitivity, while synthetic peptides with Orn (pink) or hLys (blue) resulted in a decrease in ATP sensitivity. Dashed lines indicate ATP CRCs for spliced P2X2R WT (grey) or K71Q (purple) obtained with peptide X_P2X2_. Values in (d) and (f) are displayed as mean +/- SD; n = 5-9). Source data are provided as a Source Data file.

Finally, to test whether the position of the charge at K71 is crucial for ATP recognition, we synthesized peptide X_P2X2_^SYN^ variants containing lysine and ncAA lysine derivatives (homolysine, hLys, and ornithine, Orn) at position 71 (Fig 6e), which differed only in the length of their side chains. Following recombinant expression of N and C in *Xenopus laevis* oocytes and injection of synthetic peptide, successful splicing was confirmed by functional responses to ATP application (Fig 6b). The ATP sensitivity of these responses demonstrated that the Lys-containing peptide X_P2X2_^SYN^ variant generated WT-like responses, whereas those containing hLys and Orn generated responses similar to those obtained with the conventional K71Q mutant (Fig 6f). We thus conclude that efficient recognition of ATP by P2X2Rs is highly dependent on the precise position of the charge at K71. However, ATP-generated currents were markedly smaller (<5%) than those recorded from full length protein (WT) or from the spliced product generated by co-expression of X_P2X2_^REC^ with N and C, even after attempts to increase the concentration of peptide X_P2X2_^SYN^ inserted into oocytes using multiple injections (see Methods). Additionally, functional currents took a longer time to manifest (3–5 days after peptide X_P2X2_^SYN^ injection, compared to 1 day after WT and 3 days after N+C+X_P2X2_^REC^ RNA injection), indicating slow formation of the fully spliced product. This low splicing efficiency was also evident from our inability to use immunoblotting to detect bands corresponding to full-length P2X2R. Application of the tPTS approach to incorporate hLys and Orn at position 69 (replacing a different lysine residue involved in ATP recognition) resulted in currents not distinguishable from background (Fig S12). This suggested that the modification resulted in an even larger right-shift in the ATP concentration-response curve, which cannot be accurately determined. However, despite this low overall splicing efficiency, we were able to use conventional PTS to reconstitute functional P2X2Rs from N- and C-terminal fragments expressed in HEK cells using only a single (*Cfa*DnaE) split intein (Fig S13), demonstrating that splicing within extracellular domains is feasible in both *Xenopus laevis* oocytes and mammalian cells.

## Discussion

We have demonstrated that tPTS can be employed to introduce single or multiple chemical modifications into soluble and membrane proteins in live cells. This includes combinations of ncAAs, PTMs or PTM mimics that cannot currently be incorporated into live cells using available methods. A key advantage of the tPTS approach in live cells is that the refolding step typically required with *in vitro* applications can be bypassed. This means the approach can be used for larger, more complex proteins, including those residing in the membrane. Additionally, the approach does not rely on the ribosomal machinery and thus delivers a homogenous protein population by avoiding the potential for non-specific incorporation, which can affect protein manipulation using non-sense suppression approaches^32–36^.

While tPTS offers new ways to manipulate proteins, several aspects require careful consideration for its applications in a broader context. First, the splicing efficiency in tPTS is sequence-dependent. In cases where the native sequence does not contain residues required for splicing, mutations may need to be introduced at the intended splice sites to fulfil the extein requirements for successful splicing (i.e. the need for a Cys or Ser at the +1 extein position of the extein, see Fig 1). Moreover, the protein fragment to be modified needs to be within the length limit of what is synthetically feasible. We also expect for example transmembrane sections of a protein to be less amenable to this method, as they are challenging to synthesize and insert post-translationally. Second, we note that numerous other split inteins^34^ with different extein requirements could alternatively be used for this approach and could potentially, depending on the context, yield higher splicing efficiency. Here, we chose the split inteins *Cfa*DnaE and *Ssp*DnaB^M86^, as they have been well characterized with fast kinetics and engineered to have increased tolerance to non-native extein sequences^10, 11^. Importantly, *Ssp*DnaB^M86^ can be split asymetrically with the Int^N^ segment only comprising 11 amino acids, making it an ideal split intein B in this approach (Fig 1). Finally, the means of introducing the synthetic peptide X needs to be optimized depending on the cell type in question. While synthetic peptides can be injected directly into *Xenopus laevis* oocytes, our approach requires potentially more challenging delivery techniques, such as cell squeezing, electroporation or the use of cell-penetrating peptides when implemented in mammalian cells.

Unsurprisingly, for all the proteins tested here, we note that the amount of fully spliced products generated using tPTS is generally lower, and their formation can take longer than when expressing full-length WT proteins. Factors such as molecular crowding or unfavourable spatial arrangements of protein fragments in the cell could contribute to these issues. Furthermore, it cannot be excluded that the recombinantly expressed protein fragments display different stabilities toward the proteasome or are differentially trafficked, resulting in unequal fragment ratios and thus potentially suboptimal conditions for splicing to occur. The length, proteolytic stability, and solubility of synthetic peptides, along with requirements for native-like flanking extein sequences, can also affect splicing efficiency and reaction rates^35^. Additionally, the amount of synthetic peptide that can be delivered into a cell is typically limited by the viability of the cell in response to delivery of the peptide and peptide concentration. Lastly, factors that contribute to optimal splicing conditions, such as pH or redox potential, which are controllable *in vitro*, are virtually impossible to manipulate in a live cell. Indeed, it is possible that the lower splicing efficiency we observed when using tPTS to modify the extracellular domain of the P2X2 receptor was due to unfavorable redox conditions in the endoplasmic reticulum and/or the low abundance of synthetic peptide in this subcellular compartment (or others that the splicing could take place in).

Nevertheless, it is important to appreciate that low protein yields are also not uncommon with ribosome-based approaches to genetically engineer proteins. This is particularly true for complex proteins expressed in eukaryotic cells. In fact, many groups have repeatedly observed yields of 10% or less with ncAA incorporation into transporters^36^, ion channels^37–39^, and G protein-coupled receptors^40, 41^. Although the generally low yields observed with tPTS likely restrict the approach to applications that do not require large amounts of protein, at least some of the above limitations can be addressed by engineering more promiscuous and efficient split inteins^10–12, 42^ or by adding affinity tags to promote split intein interactions^18^. Such improvements would allow the approach to be applied to a broader complement of target proteins.

The ability to apply this approach in eukaryotic cells has enabled us to use highly sensitive electrophysiology and imaging techniques to determine the presence and functionality of fully spliced products. tPTS will thus permit synthetic peptide insertion into different proteins, in particular those that are amenable to highly sensitive methods to study function or localization. Beyond the introduction of PTMs, PTM mimics and ncAAs, the approach can be used to insert virtually any chemical modification into a target protein, including backbone modifications, chemical handles, fluorescent or spectroscopic labels, and combinations thereof. This constitutes an important advantage over existing methodologies. Specifically, we anticipate that the approach will overcome some of the drawbacks associated with conventional genetic engineering in eukaryotic cells (non-specific incorporation, premature termination, dependence on ribosomal promiscuity^36, 43^) and semi-synthetic approaches that require protein refolding^7^. It will thus increase the number and type of functionalities that can be incorporated into proteins that prove amenable to tPTS.

## Methods

### Molecular biology

Plasmid DNAs were purchased from GeneArt (ThermoFisher scientific), General biosystems Inc. or Twist Bioscience. All gene constructs were sub-cloned into either the pUNIV or pcDNA3.1+ backbone. pUNIV backbone was a gift from Cynthia Czajkowski (Addgene plasmid # 24705; http://n2t.net/addgene:24705; RRID:Addgene_24705). Conventional site-directed mutagenesis was performed using standard PCR. Complementary RNA (cRNA) for oocyte microinjection was transcribed from respective linearized cDNA using the Ambion mMESSAGE mMACHINE T7 Transcription Kit (Thermo Fisher Scientific).

### Peptide synthesis

Peptides for GFP splicing were sourced from Proteogenix, France. Peptides for Na_v_1.5 and P2X2R splicing were synthesized by solid-phase peptide synthesis (details in Supplementary material). Peptide X variants were synthesized as 3 shorter fragments and assembled in a one-pot native chemical ligation procedure, as briefly outlined below. The split intein-mediated reconstitution of proteins developed here required the synthesis of a small collection of peptides between 69 and 77 amino acids in length. Conveniently, all peptide X variants needed for our work share identical Int^C^-A (35 amino acids) and Int^N^-B (11 amino acids) sequences, which flank the sequence corresponding to the protein of interest (POI). In order to reduce the synthesis demands, we took advantage of the sequences of Int^C^ and Int^N^ (i.e. Cys residues at +1 position in the exteins) by adopting a ‘one-pot’ chemical ligation strategy of three parts (Int^C^-A, POI segment and Int^N^-B), with the sequence from the POI being the only variable one. For this purpose, a C-to-N-directed ligation strategy based on Thz masking of cysteine^44^ was implemented for the assembly of the peptide X variants (Fig S2). For the assembly of peptide X variants containing a thio-acetylated lysine, a different ligation strategy (N-to-C directed) was adopted (Fig S2) in order to avoid the Thz-cysteine unmasking step (acidic pH at 37°C) of the C-to-N-directed ligation. Indeed our collaborators experienced partial conversion of similar thioamide-containing peptides into amides during the HPLC purification step, performed in water–MeCN containing 0.1% TFA (personal communication with Dr Christian A. Olsen, data not published).

### Expression in Xenopus laevis oocytes

cRNAs were injected into *Xenopus laevis* oocytes (prepared as previously described^38^) and incubated at 18 °C in OR-3 solution (50 % Leibovitz’s medium, 1 mM L-Glutamine, 250 mg/L Gentamycin, 15 mM HEPES, pH 7.6) for up to 7 days. For injection of synthetic peptides, lyophilized peptides were dissolved in Milli-Q H_2_O to a concentration of 750 µM and 18 nL of solubilized peptide was injected into cRNA pre-injected oocytes with the *Nanoliter 2010* micromanipulator (World Precision Instruments). For Na_V_1.5 constructs, synthetic peptides were injected 1 day after cRNAs were injected and recordings performed 12-20 hrs after peptide injection. For P2X2 constructs, synthetic peptides were injected consecutively on days 2, 3 and 4 following cRNA injection and recordings performed on day 7.

### Two-electrode voltage clamp (TEVC) recordings

Voltage or ATP-induced currents were recorded with two microelectrodes using an OC-725C voltage clamp amplifier (Warner Instruments). Oocytes were perfused in ND96 solution (in mM: 96 NaCl, 2 KCl, 1 MgCl_2_, 1.8 CaCl_2_ /BaCl_2_, 5 HEPES, pH 7.4) during recordings. Glass microelectrodes were backfilled with 3 M KCl and microelectrodes with resistances between 0.2 and 1 MΩ were used. Oocytes were held at −100 mV (for Na_V_1.5 constructs) or −40 mV (for P2X2 constructs). For Na_V_1.5, sodium currents were induced by +5 mV voltage steps from −80 mV to +40 mV. Steady-state inactivation was measured by delivering a 500 ms prepulse from −100 mV to −20 mV in +5 mV voltage steps followed by a 25 ms test pulse of −20 mV. For P2X2 recordings, ATP-induced currents were elicited through application of increasing concentrations of ATP (dissolved in ND96, pH7.4) supplied via an automated perfusion system operated by a ValveBank™ module (AutoMate Scientific).

### Immunoblots

Oocytes expressing full-length receptors or different combinations of the split-intein receptor fragment fusion proteins were isolated 3–4 days after RNA injection and washed twice with PBS. Total cell lysates were obtained by lysing the oocytes in Pierce™ IP lysis buffer with added Halt protease inhibitor cocktail (Thermo Fisher Scientific). Surface proteins were purified with the Pierce™ Cell Surface Protein Isolation Kit (Thermo Fisher Scientific). Purified surface proteins or total cell lysates were run on a 4–12 % BIS-TRIS gel (for P2X2) or 3-8 % Tris-acetate gel (for Na_V_1.5) and transferred to a PVDF membrane. Membranes were incubated with rabbit polyclonal anti-Na_V_1.5 (#ASC-005, Alomone labs; 1:2000), anti-Na_V_1.5 (#ASC-013, Alomone labs; 1:1500) or anti-P2X2 Antibody (#APR-003, Alomone labs; 1:2000) and the bound primary antibodies were detected by a HRP-conjugated goat anti-rabbit secondary antibody (W401B, Promega; 1:2000). Membranes were developed and visualized using the Pierce™ ECL immunoblotting substrate (ThermoFisher Scientific).

### Expression in HEK293 cells

HEK293 cells (American Type Culture Collection) were grown in Dulbecco’s modified Eagle’s Medium (DMEM) (Gibco) supplemented with 10 % Fetal Bovine Serum (Gibco), 100 units/mL penicillin and 100 μg/mL streptomycin (Gibco) and incubated at 37 °C with 5 % of CO_2_. Confluent cells growing in monolayers were washed with 10 mL phosphate-buffered saline (PBS) (in mM: 137 NaCl, 2.7 KCl, 4.3 Na_2_HPO_4_, 1.4 KH_2_PO_4_ (pH 7.3)), detached with trypsin-EDTA (Thermo Fisher Scientific) and re-suspended in DMEM. The re-suspended cells were seeded onto glass coverslips pre-treated with poly-L-Lysine in 35-mm dishes for patching or in 35-mm glass bottom dishes for imaging and incubated for 24 hrs. prior transfection. The plated HEK293 cells were transfected using TransIT DNA transfection reagent (Mirus) following the instructions supplied by the manufacturer and incubated until use. For imaging of reconstituted GFP in HEK293 cells, DNA coding for three GFP-split intein fusion fragments (N, X and C) was inserted into the pcDNA3.1+ vector by GeneArt (Thermo Fisher Scientific) and co-transfected in a 1:1:1 ratio using a total of 3 µg DNA and incubated for 48 hrs. before imaging. In parallel, a batch of cells was transfected with WT GFP as a positive control and in addition five batches of cells were transfected with DNA coding for two fragments of the GFP alone (N+X, N+C, X+C) or combined with a non-splicing GFP fragment (N+X-Cys65Ala+C or N-X+C-Ser85Ala) as negative controls. To keep the same amount of DNA for each combination pcDNA3.1 + empty vector was co-transfected for the control experiments. For P2X2R patch-clamp recordings, HEK293 cells were transfected in a 30 mm dish with 1.5 µg DNA of each construct, respectively (N+C, N+C_mut_, N, C) and incubated for 2 days at 37 °C.

### Imaging of reconstituted GFP

Imaging was performed using an inverted microscope *IX73* (Olympus) with 10X and 20X objectives mounted on a motorized nosepiece (Olympus) controlled by a *CMB U-HSCBM* switch and connected to a *DCC1545-M* camera (ThorLabs). GFP fluorescence was visualized using a LED light source (CoolLed pE-100, 470nm).

### Peptide transfer by cell squeezing

Squeezing was performed using a chip with constrictions of 6 µm in diameter and 10 µm in length (CellSqueeze 10-(6)x1, SQZbiotech). In all microfluidic experiments, a cell density of 1.5.10^6^ cells/mL in Opti-MeM was squeezed through the chip at a pressure of 40 psi. Transduction was conducted at 4 °C to block cargo uptake by endocytosis^45^. During squeezing, a peptide concentration of 10-20 µM in the surrounding buffer was used. After squeezing, cells were incubated for 5 min at 4 °C to reseal the plasma membrane. Squeezed cells were washed with DMEM containing 10 % FCS, seeded into 8-well on cover glass II slides (Sarstedt) coated with fibronectin (5 µg/mL) in DMEM containing 10% FCS, and cultured at 37 °C and 5 % CO_2_. As a control for endosomal uptake, cells were incubated with 10 µM of peptide at RT without microfluidic cell manipulation. Confocal imaging was performed 1, 2, 4, 8 h and 20 h after squeezing. Before imaging, cells were washed with PBS (Sigma-Aldrich), fixed with 4 % formaldehyde (Roth)/PBS for 20 min at 20 °C and quenched by the addition 50 mM glycine in PBS (10 min, 20 °C).

### Confocal laser scanning microscopy

Imaging was performed using the confocal laser scanning (LSM) microscope LSM880 (Zeiss) with Plan-Apochromat 20x/1.4 Oil DIC objective. The following laser lines were used for excitation: 405 nm for blue-shifted GFP and 488 nm for GFP. ImageJ^46^, Fiji^47^, and Zen 2.3 black (Carl Zeiss Jena GmbH, Germany) were used for image analysis.

### Patch-clamp electrophysiology

The cells were reseeded 1 to 4 hours before the patch-clamp experiments. Ionic currents were recorded with borosilicate patch pipettes (2-5 MΩ) filled with intracellular solution (in mM: 140 KCl, 5 MgCl_2_, 5 EGTA, 10 HEPES, pH 7.3) at −40 mV with the Axopatch 200B amplifier and the 1550A digitizer (Molecular Devices). Lifted cells were perfused with extracellular solution (in mM: 140 NaCl, 2.8 KCl, 2 CaCl_2_, 2 MgCl_2_, 10 HEPES, 10 Glucose, pH 7.3) and activated with 1 mM ATP via a piezo-actuated perfusion tool.

## Acknowledgements

We acknowledge the Lundbeck Foundation (R139-2012-12390 to SAP), the Carlsberg Foundation (CF16-0504 to SAP), the Independent Research Fund Denmark (7025-00097A to SAP), the University of Copenhagen, and the German Research Foundation (SFB 807, SPP 1623 and GRK 1986 to RT) for financial support. RT would like to acknowledge the support by an ERC Advanced Grant from the European Research Council. We thank Dr Christian A Olsen for support with the peptide chemistry, Janne Colding and Natasha Gray-Garney for technical support, and Drs Marlieke JM Jongsma and Huib Ovaa for help with the FACS experiments. We would also like to thank Drs Lesley Anson, Christian A Olsen and Kristian Strømgaard and members of the Pless lab for helpful comments on the manuscript.

## Author contributions

K.K.K., I.G. and S.A.P. designed the research. K.K.K., I.G., F.G., R.W., H.H., M.H.P., H.C.C, M.W. performed the experiments. K.K.K., I.G., F.G., R.W., H.H., M.H.P., H.C.C, M.W. analyzed the data. R.T. and S.A.P. supervised the project. K.K.K., I.G. and S.A.P. wrote the manuscript with input from all authors.

## Competing interests

The authors declare to have no competing interests.

## Data availability

The source data underlying Figs 2c-e, 3c,e-f, 6c,d,f, and Supplementary Figs S1c, S3a, S4c-e,h-j, S5c, S8b, S9d-e, S10d-e, S11d are provided as a Source Data file on Zenodo (DOI: 10.5281/zenodo.3712821).

## Supplementary Information

### Supplementary Figures

**Fig S1:**
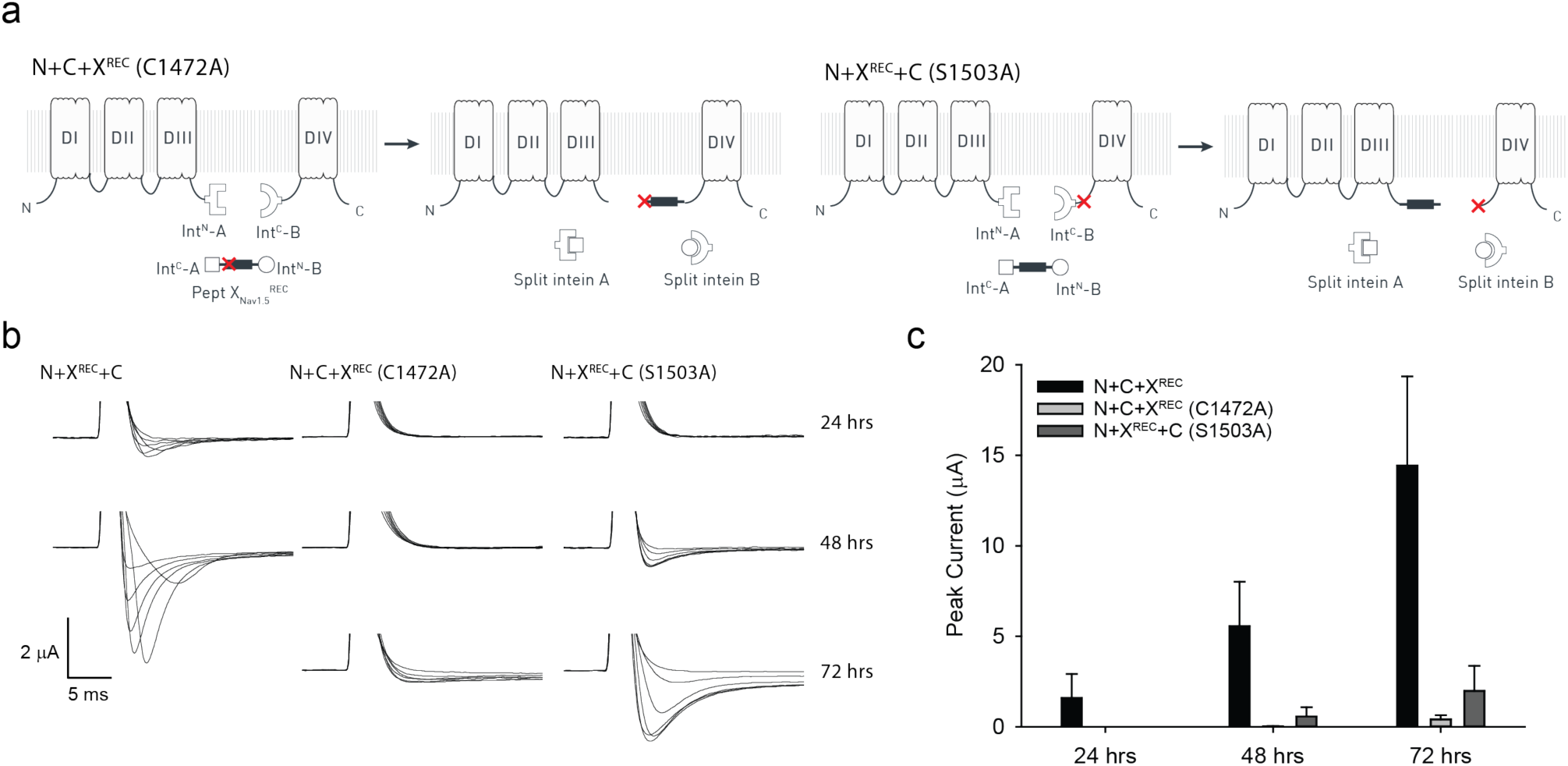
Control experiments for tPTS in Na_V_1.5 DIII-DIV linker (recombinant expression). (a) Schematic presentation of non-spliced Na_V_1.5 fragments expected for the Na_V_1.5 DIII-DIV linker splice sites tested when +1 extein residues of each split intein is mutated to alanine (indicated by red cross): C1472A in intein A and S1503A in intein B. This prevents splicing and favors side reactions, resulting in accumulation of cleavage products (note that this will likely overrepresent the occurrence of the side reactions compared to when splicing-competent split inteins are used, i.e. Fig 1). (b) Representative families of sodium current traces in response to voltage steps from −50 mV to +10 mV in 10 mV steps, recorded 24, 48, or 72 hrs after mRNA injection of N+X^REC^+C. Note that oocytes with peak currents exceeding 2 µA are typically not ideally voltage-clamped, which decreases the accuracy of the values obtained. As expected, functional non-spliced constructs (e.g. N+X^REC^+C [S1503A]) display currents with impaired (slower) inactivation. (c) Average peak currents recorded for each combination depicted as a bar plot (mean +/- SD; n = 3-7) for the respective time points (note that experiments shown in Fig 1 were conducted 24 hours after N+C mRNA injection or 12-20 hrs after injection of peptide X^SYN^). Source data are provided as a Source Data file.

**Fig S2:**
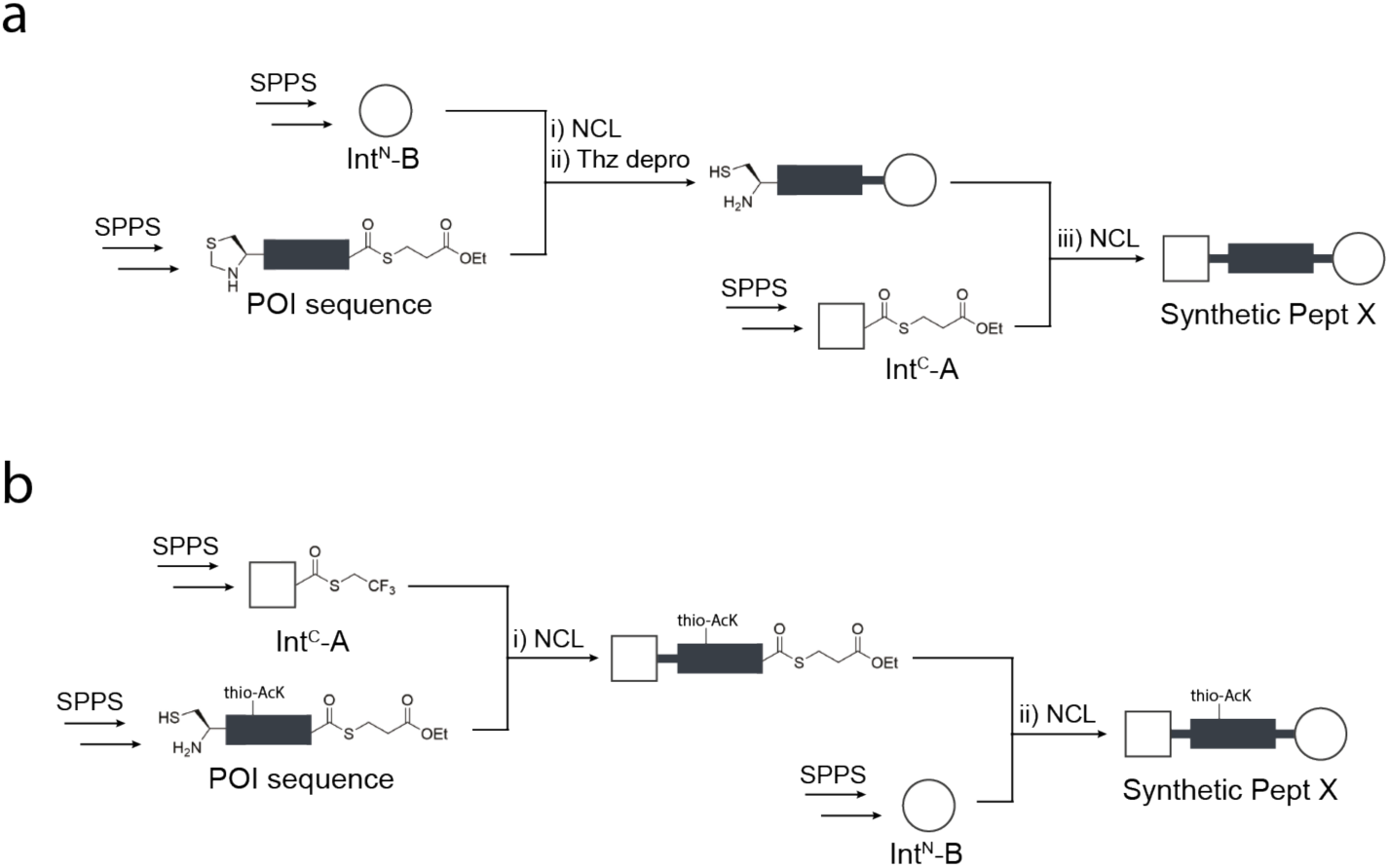
Peptide X synthesis strategies. (a) C-to-N ‘one-pot’ ligation of three parts, which were synthesized by standard SPPS. The two ligation steps were performed by exploiting the standard cysteine Thz-masking approach. (b) For peptides X containing a thio-acetylated lysine residue, an alternative N-to-C ‘one-pot’ ligation was adopted. This strategy exploited the faster kinetics of the TFET thioester of Int^C^-A compared to the alkyl thioester of the POI peptide, allowing us to avoid orthogonal protecting groups, e.g. Thz.

**Figure S3:**
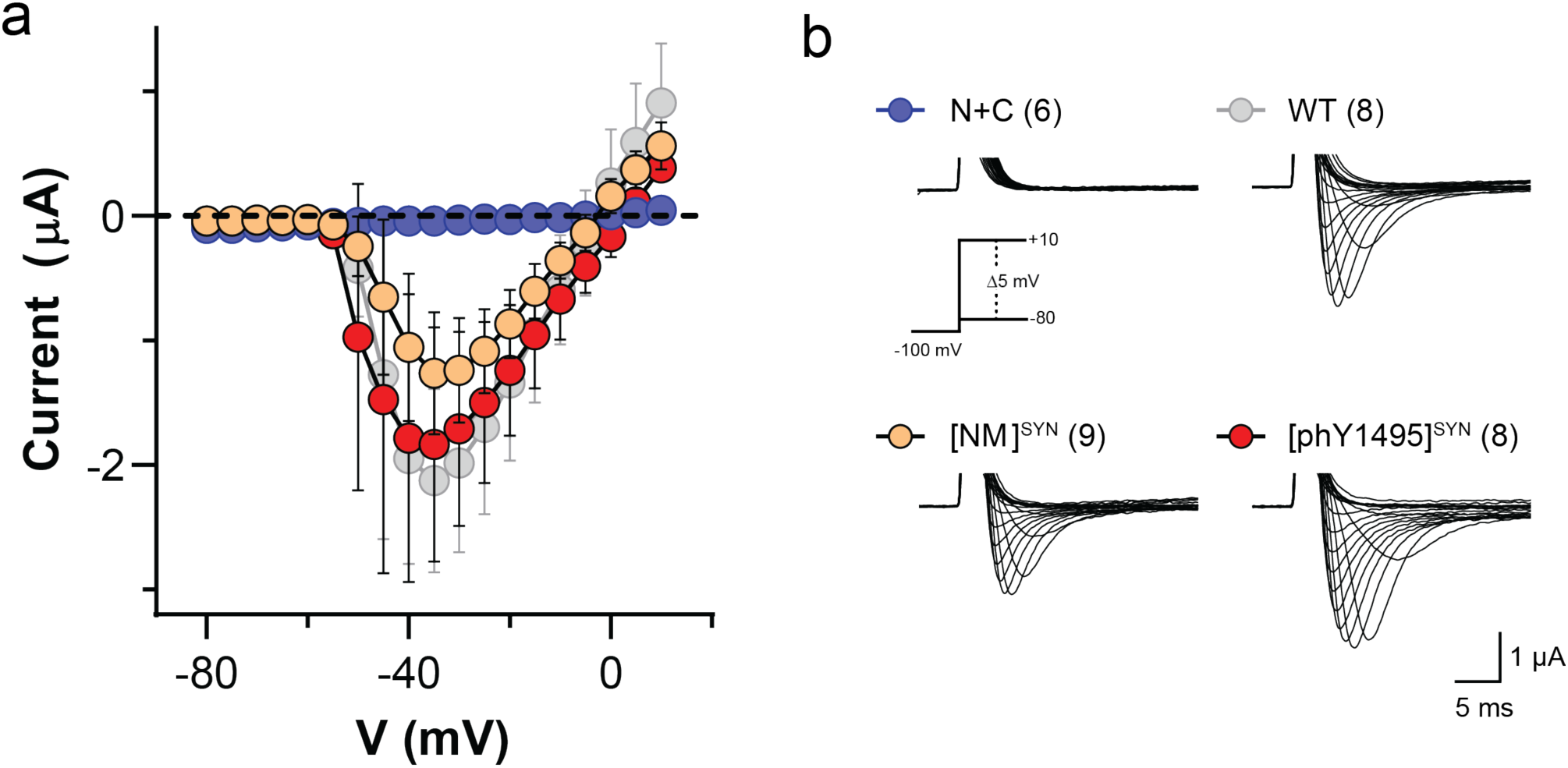
Spliced Na_V_1.5 channels containing synthetic peptide X showed depolarization-evoked currents of amplitudes comparable to WT Na_V_1.5. (a) Current-voltage relationships and (b) representative sodium currents of *Xenopus laevis* oocytes expressing N and C constructs only (N+C), Na_V_1.5 (WT), N+C+X^NM^ ([WT^NM^]^SYN^) and N+C+X^phY1495^ ([phY1495]^SYN^) in response to depolarizing voltage steps (−80 to +10 mV in 5 mV steps with a holding potential of −100 mV). *Xenopus laevis* oocytes were injected with RNAs of N+C or WT Na_V_1.5 48 hours before recording, whereas synthetic peptides X (X^NM^ or X^phY1495^) were injected into N+C RNA-injected oocytes 24 hours before recording. To ensure adequate control of voltage clamp, [Na^+^] in the extracellular recording solution was reduced, (in mM): 24 NaCl, 72 NMDG, 2 KCl, 1.8 CaCl_2_, 1 MgCl_2_ and 5 HEPES, pH 7.4 with HCl. Data in (a) are shown as mean ± SD; n=6-9. Numbers in parentheses (b) indicate number of individual cells used for recordings. Source data are provided as a Source Data file.

**Fig S4:**
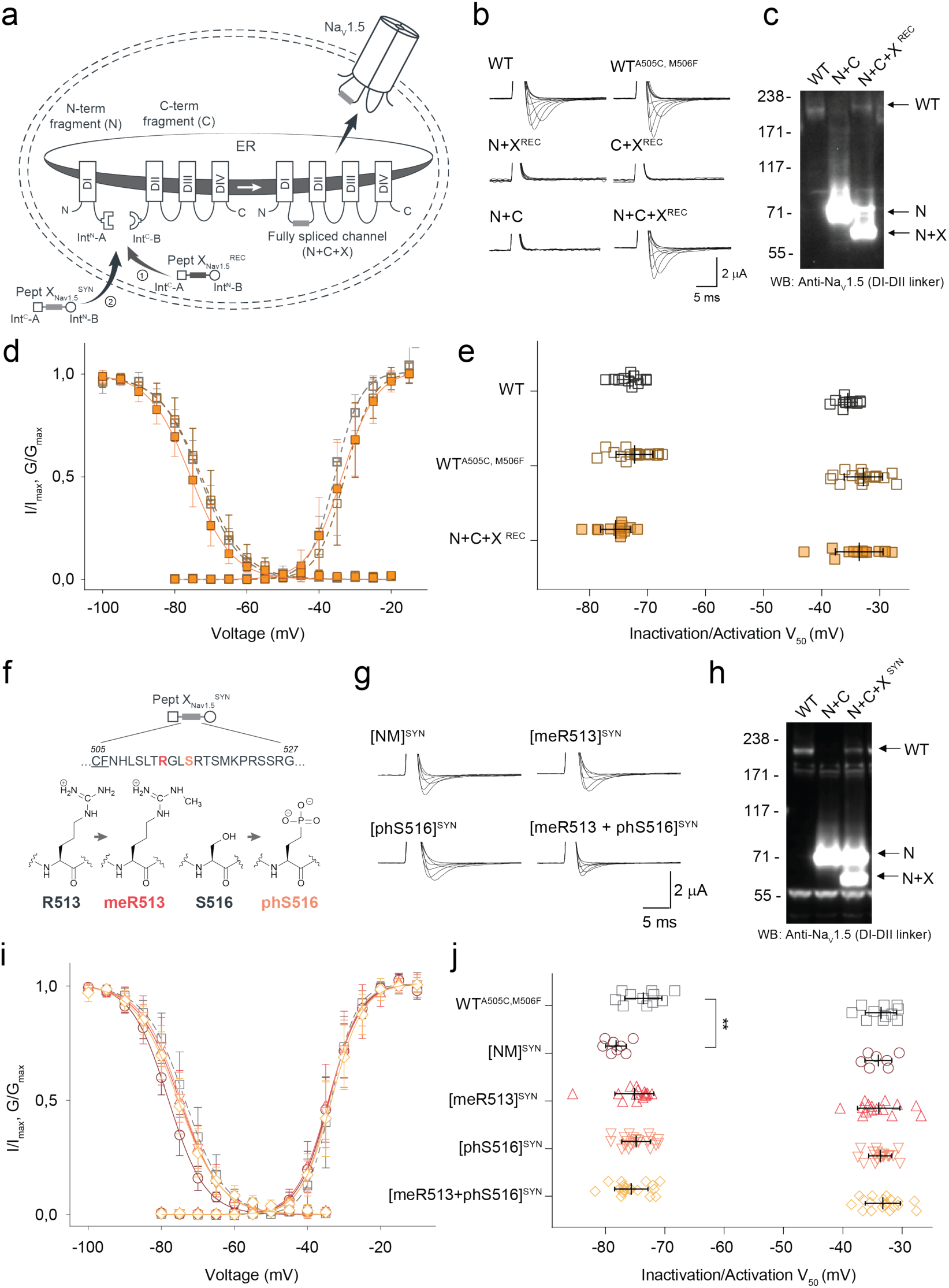
tPTS to insert recombinant or synthetic peptides into the Na_V_1.5.DI-DII linker. (a) Schematic overview of the applied approach using either recombinant (strategy 1) or synthetic peptide X_Nav1.5_^SYN^ (corresponding to amino acids 505 to 527 of the Na_V_1.5 DI-DII linker; strategy 2) to reconstruct full-length Na_V_1.5 from recombinantly expressed N-/C-terminal fragments (N and C) in *Xenopus laevis* oocytes. Inteins A (*Cfa*DnaE) and B (*Ssp*DnaB^M86^) are indicated by square and round symbols, respectively. (b) Representative sodium currents in response to sodium channel activation protocol (see methods; only voltage steps from −50 mv to +10 mV in 10 mV steps are displayed), demonstrating expression of functional Na_V_1.5 only in the presence of all three recombinantly expressed components (N+X^REC^+C), along with WT and A505C, M506F double mutant channels (introduced to create a functional splice site for intein A). (c) Western blot analysis verifying presence of fully spliced Na_V_1.5 only when all three components (N+X^REC^+C) were co-expressed (using antibody against Na_V_1.5 DI-DII linker residues 493-511). (d) Steady-state inactivation and conductance-voltage (G-V) relationships for respective full-length and spliced constructs. (e) Comparison of values for half-maximal (in)activation (V_50_) (values are displayed as mean +/- SD; n = 12-17). (f) Sequence of X_Nav1.5_ corresponding to the amino acids replaced in the Na_V_1.5 DI-DII linker and chemical structures of native amino acids and PTMs incorporated into the respective positions of the DI-DII intracellular linker via chemical synthesis of peptide X_Nav1.5_. Note that a non-hydrolysable phosphonylated serine was used (phS). Underlined residues indicate A505C, M506F mutations that were introduced to optimize the splicing reaction. (g) Representative sodium currents in response to voltage steps from −50 mV to +10 mV in 10 mV steps, demonstrating expression of functional Na_V_1.5 when *Xenopus laevis* oocytes expressing N and C constructs were injected with synthetic peptides containing non-modifiable side chains in positions 513 and 516 (R513K and S516V, NM), meR513 or phS516 or both PTMs together. (h) Immunoblot verifying presence of fully spliced Na_V_1.5 only when synthetic peptide X_Nav1.5_^SYN^ was injected into cells expressing N and C-terminal fragments (using antibody against Na_V_1.5 DI-DII linker residues 493-511). (i) Steady-state inactivation and conductance-voltage (G-V) relationships for PTM-modified/non-modified constructs. (j) Comparison of values for half-maximal (in)activation (V_50_) (values are displayed as mean +/- SD; n = 8-21). Significant differences were determined by one-way ANOVA with a Tukey post-hoc test. **, p<0.003. Source data are provided as a Source Data file.

**Fig S5:**
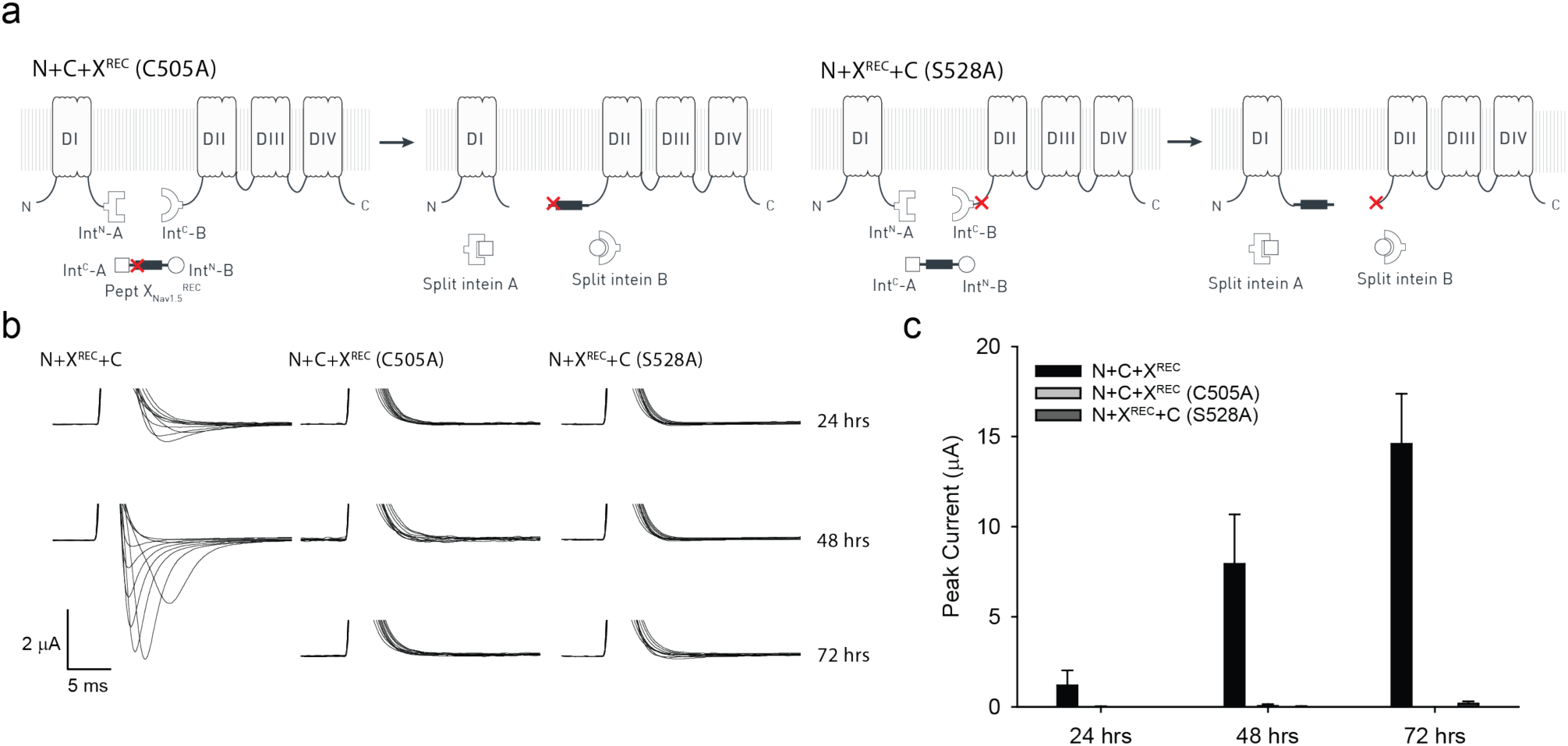
Control experiments for tPTS in Na_v_1.5 DI-DII linker (recombinant expression). (a) Schematic presentation of non-spliced Na_v_1.5 fragments expected for the Na_v_1.5 DI-DII linker splice sites tested when +1 extein residues of each split intein is mutated to alanine (indicated by red cross): C505A in intein A and S528A in intein B. This prevents splicing and favors side reactions, resulting in cleavage products to accumulate (note that this will likely overrepresent the occurrence of the side reactions compared to when splicing-competent split inteins are used, i.e. Fig S3). (b) Representative sodium current traces in response to voltage steps from −50 mV to +10 mV in 10 mV steps, recorded 24, 48 or 72 hours after mRNA injection of N+X^REC^+C. Note that oocytes with peak currents exceeding 2 µA are typically not ideally voltage-clamped, which decreases the accuracy of the values obtained. (c) Average peak currents recorded for each combination depicted as a bar plot (mean +/- SD; n= 3-11) for the respective time points (note that experiments shown in Fig S3 were conducted 24 hrs after mRNA injection or 12-20 hrs after injection of peptide X_Nav1.5_^SYN^). Source data are provided as a Source Data file.

**Fig S6:**
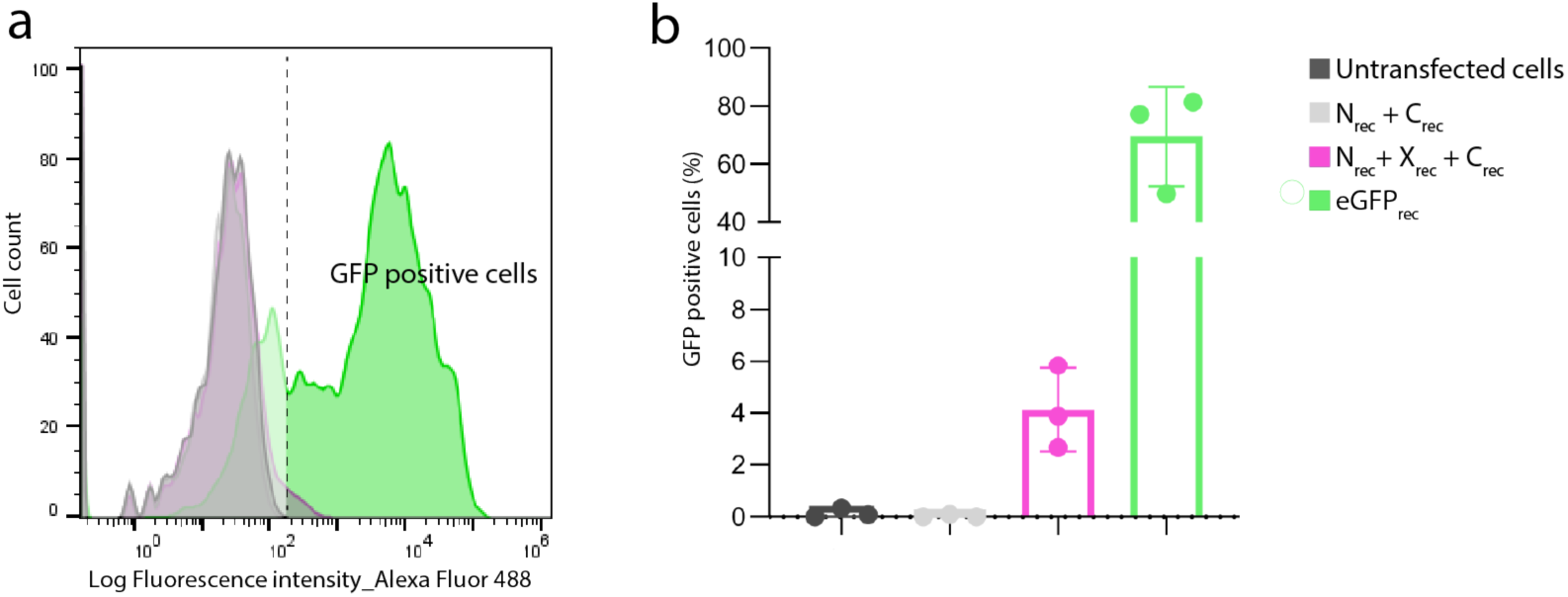
FACS analysis of reconstituted eGFP using cytosolic tPTS in mammalian cells. HEK293 cells were transfected with DNA coding for N- or C-terminal GFP fragments only (N_rec_ + C_rec_; grey) or together with peptide X_GFP_^REC^ (N_rec_ + X_rec_ + C_rec_; pink). Cellular fluorescence was analyzed and compared to that of untransfected cells (black) and eGFP transfected cells (green) using a BD™ LSR II flow cytometer. (a) Representative histogram plot of the fluorescence distribution from the analyzed cells. The fluorescence intensity cut off determining GFP positive cells (indicated by the dashed line) was set at the upper limit of fluorescence distribution from the untransfected cell population (<0.1% of untransfected cells fall in this area). (b) Bar graph showing % of GFP positive cells for each combination represented (mean +/- SD; n= 3).

**Fig S7:**
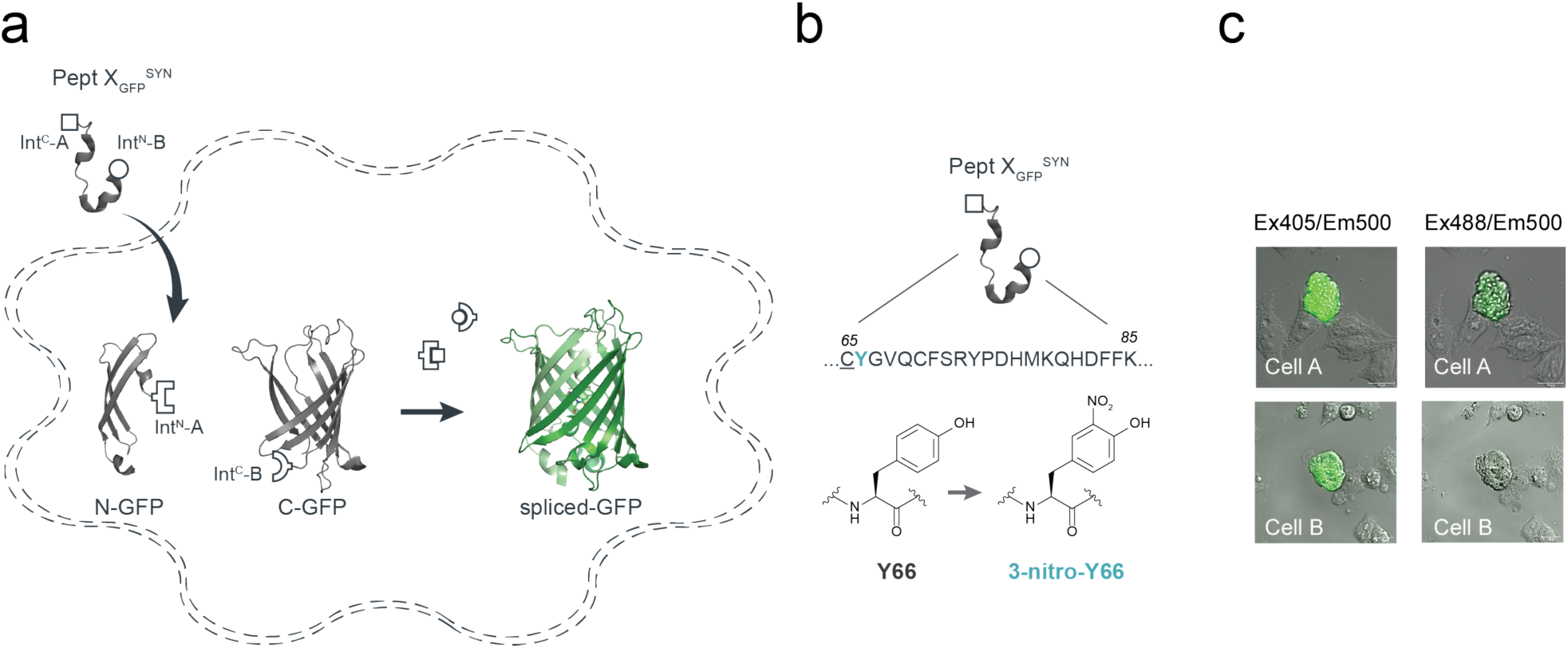
Insertion of synthetic peptide X_GFP_^SYN^ into GFP using cytosolic tPTS in mammalian cells. (a) Schematic presentation of the strategy applied to reconstitute GFP from recombinantly expressed N-/C-terminal fragments and synthetic peptide X_GFP_^SYN^ corresponding to amino acids 65-85 of GFP in HEK293 cells. Inteins A (*Cfa*DnaE) and B (*Ssp*DnaB^M86^) are indicated by square and round symbols, respectively. (b) Sequence of peptide X_GFP_^SYN^ corresponding to the amino acids replaced in GFP and chemical structures of tyrosine and its ncAA derivative (3-nitro-tyrosine) incorporated into position66 within the GFP chromophore via chemical synthesis of peptide X_GFP_^SYN^. (c) Two examples (cell A and B, respectively) of overlaid brightfield and fluorescence images of HEK293 cells transfected with N and C fragments and squeezed in the presence of peptide 20 µM X containing 3-nitro-Tyr in position 66. Note the brighter fluorescence emission when cells were excited with a 405-nm laser compared to excitation with a 488-nm laser, indicating a blue-shift in the spectral properties of the 3-nitro-tyrosine-containing GFP variant. Scale bars: 20 µm.

**Fig S8:**
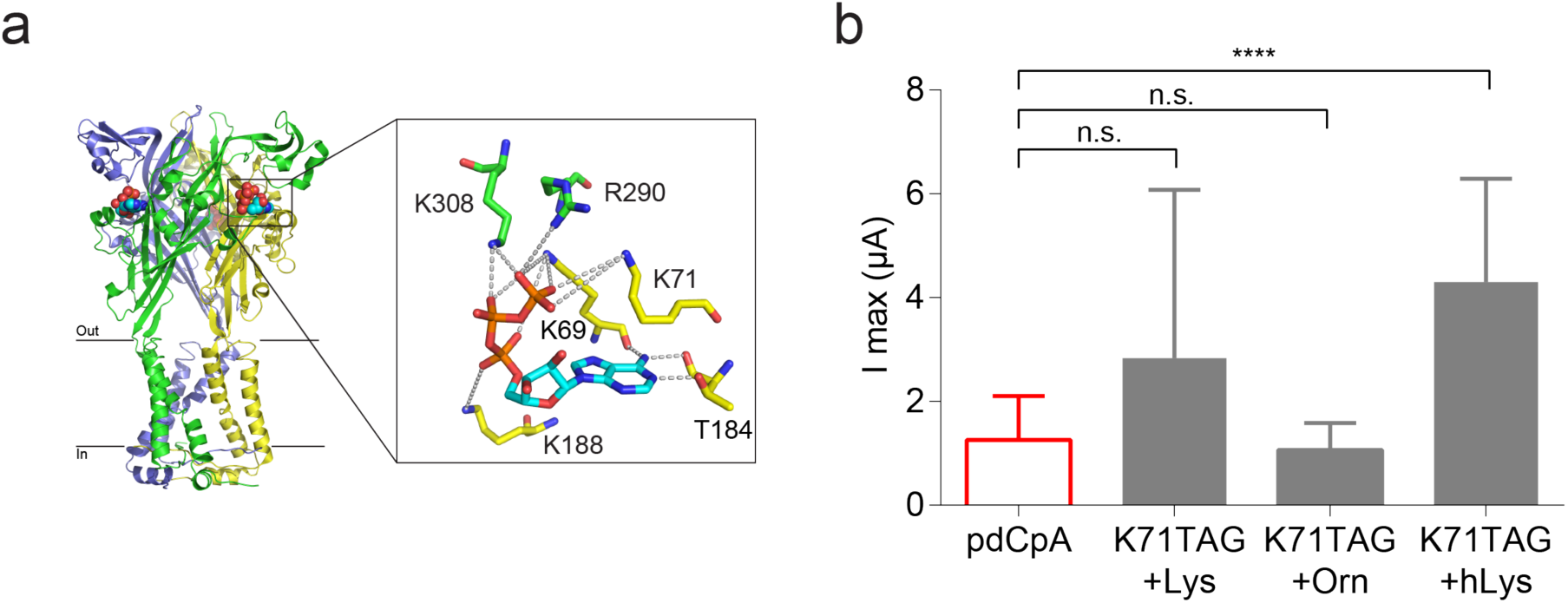
Incorporation of lysine derivatives in K71 position of the P2X2R using ribosome based non-sense suppression method in *Xenopus laevis* oocytes. (a) Crystal structure of ATP-bound hP2X3R (PDB: 5svk). Inset highlights interaction of K71 (P2X2 residue numbering) with the phosphate tail of ATP. (b) Peak currents elicited by 50 mM ATP at oocytes injected with uncharged tRNA (pdCpA, red bar) or mRNA coding for K71TAG with lysine, ornithine (Orn) or homolysine (hLys) charged tRNAs. Non-significant differences in peak current size recorded from oocytes injected with uncharged and charged tRNAs (with the exception of hLys) indicate possible non-specific incorporation of endogenous amino acids^1^ at the K71TAG site. Values are depicted as a bar plot (mean +/- SD, n = 5-27). Significant differences were determined by one-way ANOVA with a Tukey post-hoc test. n.s., p>0.03; ****, p<0.0001. Source data are provided as a Source Data file.

**Fig S9:**
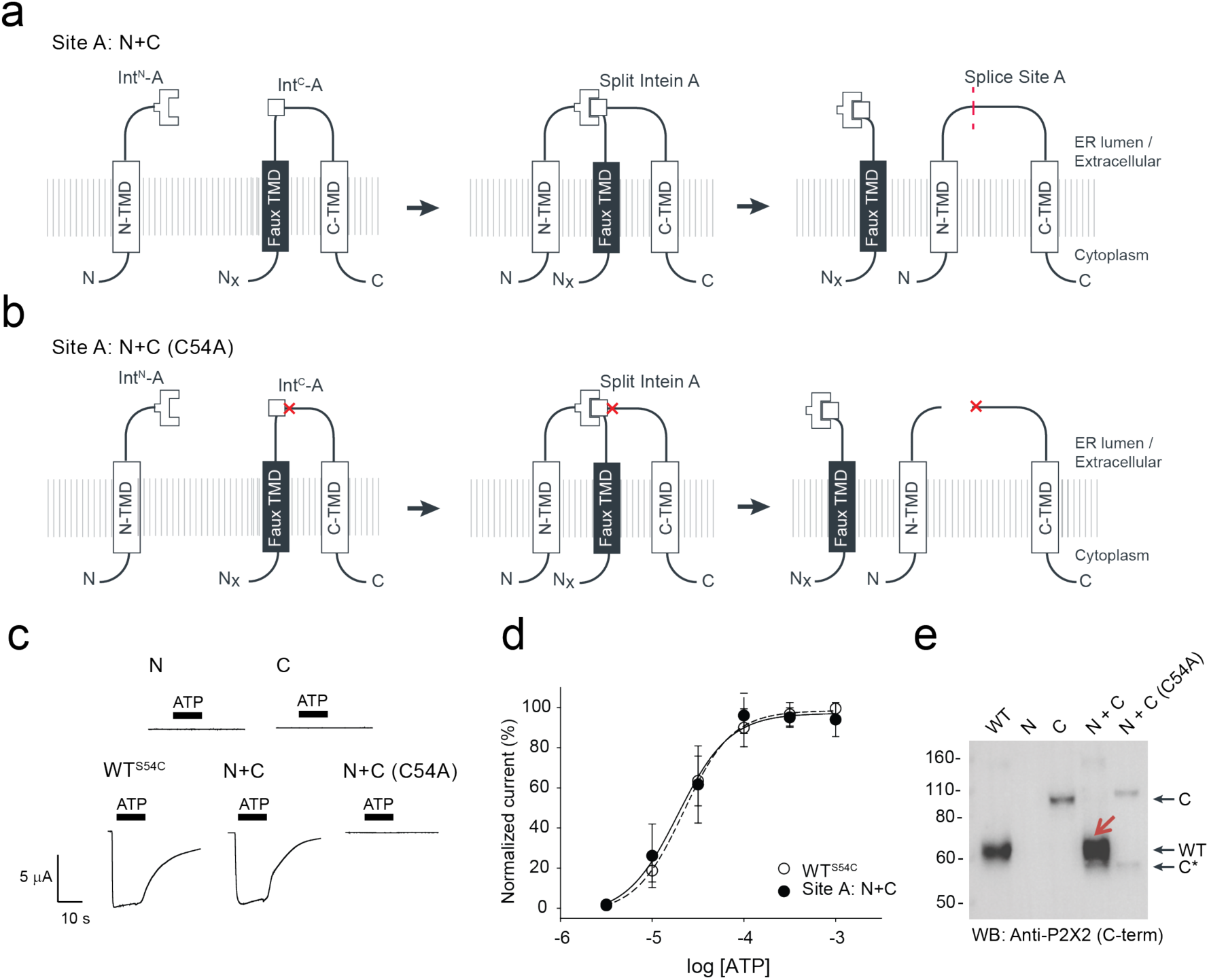
Single-intein PTS of the P2X2R extracellular domain at splice site A. (a) Schematic presentation of single-intein PTS at position 54 (site A) in the extracellular domain of P2X2Rs in *Xenopus laevis* oocytes. Note that a *faux* transmembrane helix was engineered into the C-terminal fragment to maintain its native membrane topology during protein expression. S54C mutation (at splice site A in P2X2) was introduced in the C-terminal fragment to create an optimized splice site for intein A. Intein A (*Cfa*DnaE) is indicated by square symbols. (b) Schematic presentation of non-spliced P2X2R fragments expected when +1 extein residue of split intein A (position 54 of P2X2) is mutated to alanine (indicated by red cross). This prevents splicing and favors side reactions, resulting in cleavage products to accumulate (note that this will likely overrepresent the occurrence of the side reactions compared to when splicing-competent split inteins are used). (c) Representative current traces during application of ATP (300 µM, indicated by black bars) to oocytes expressing respective constructs recorded one day after injection of mRNA. ATP-induced currents were absent under control conditions even after multiple days after injection. (d) Concentration-response curve (CRC) of reconstituted receptor indicates wild-type like functionality. Values are displayed as mean +/- SD; n = 5-7. (e) Western blot analysis of surface-purified proteins verifies splicing of the full-length receptor only when all required components were present (indicated with red arrow). Antibody targeting the C-terminus of P2X2 was used. Black arrows on the right indicate band positions of the respective constructs (Actual Mw of constructs: WT, 53 kDa; C-construct, 85 kDa; C-terminal cleavage product, C*, 46 kDa). Source data are provided as a Source Data file.

**Fig S10:**
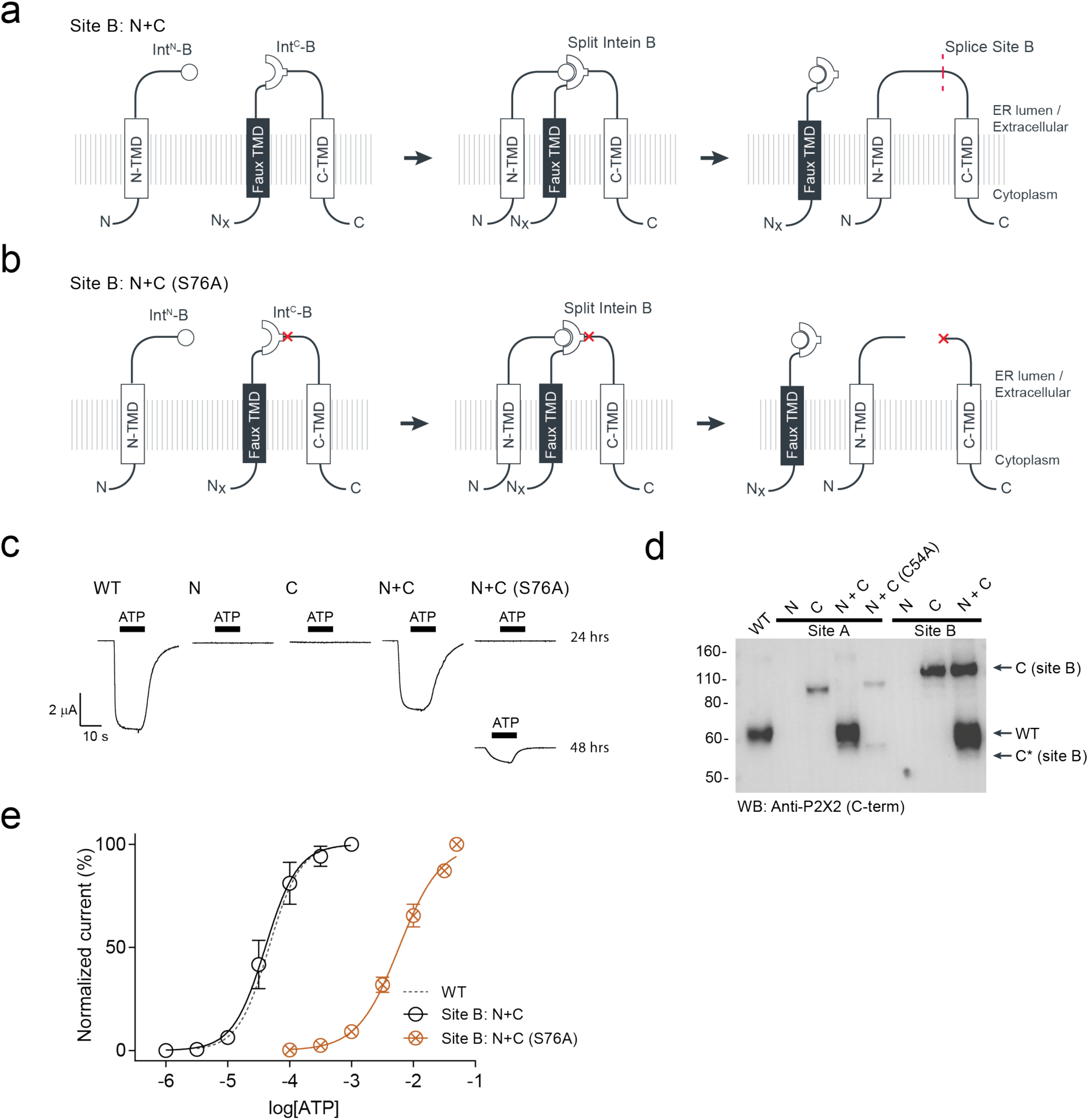
Single-intein PTS of the P2X2R extracellular domain at splice site B. (a) Schematic presentation of single-intein PTS at position 76 (site B) in the extracellular domain of P2X2Rs in *Xenopus laevis* oocytes. Note that a faux transmembrane helix was engineered into the C-terminal fragment to maintain its native membrane topology during protein expression. Intein B (*Ssp*DnaB^M86^) is indicated by round symbols. (b) Schematic presentation of non-spliced P2X2 fragments expected when +1 extein residue of split intein B (position 76 of P2X2) is mutated to alanine (indicated by red cross). This prevents splicing and favors side reactions, resulting in cleavage products to accumulate (note that this will likely overrepresent the occurrence of the side reactions compared to when splicing-competent split inteins are used). (c) Representative current traces during application of 1 mM ATP (indicated by black bars) to oocytes expressing respective constructs recorded one day after injection of mRNA. ATP-induced currents were absent when N and C constructs were expressed individually. ATP-induced currents were observed only 2 days after injection of N+C (S76A) mRNA. (d) Western blot analysis of surface-purified proteins verifies splicing of the full-length receptor only when all required components were present. Antibody targeting the C-terminus of P2X2 was used. Black arrows on the right indicate band positions of the respective site B constructs (Actual MW of constructs: WT, 53 kDa; C-construct, 97 kDa; C-terminal cleavage product, C*, 44 kDa). Note that band positions and MW of site A constructs (lanes 2-5 of the blot) are presented in Fig S8. (e) Concentration-response curve (CRC) of reconstituted receptor indicates wild-type like functionality. Functional non-spliced construct has a significantly right-shifted CRC compared to WT. Values are displayed as mean +/- SD; n = 4-7. Source data are provided as a Source Data file.

**Fig S11:**
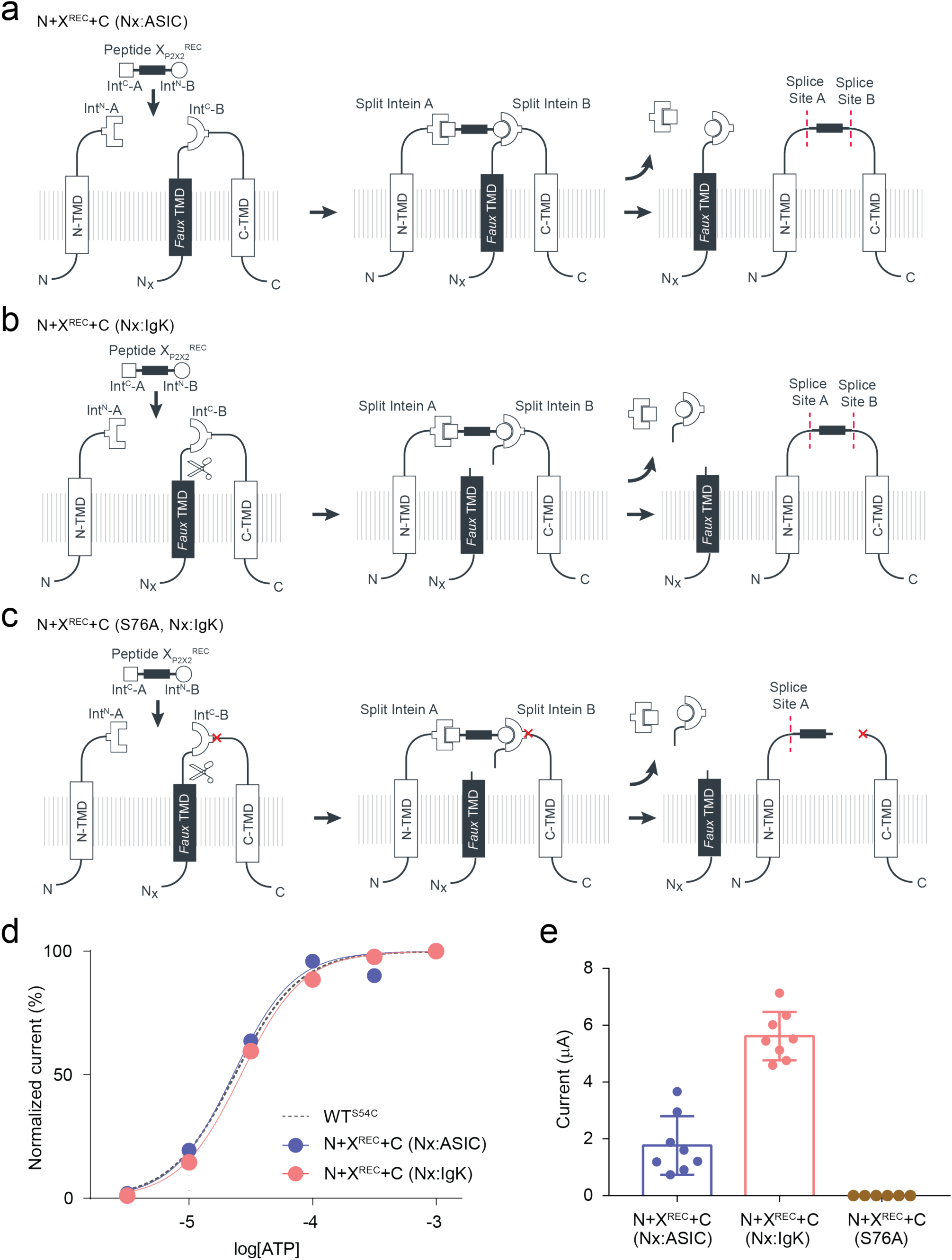
Alternative *faux* TMDs used for C-constructs in tPTS of the P2X2R extracellular domain. (a) Schematic presentation of tPTS applied to the P2X2R extracellular domain where a non-cleavable *faux*TMD (ASIC1a N-TMD sequence, Nx:ASIC) was used for the C-construct. *Faux* transmembrane helix was engineered into the C-terminal fragment to maintain its native membrane topology during protein expression. Inteins A (*Cfa*DnaE) and B (*Ssp*DnaB^M86^) are indicated by square and round symbols, respectively. Full-length P2X2Rs were reconstructed by recombinantly co-expressing N-/C-terminal fragments (N and C) and a recombinant peptide corresponding to amino acids 54 to 75 of the P2X2R extracellular domain (peptide X_P2X2_^REC^; X^REC^) in *Xenopus laevis* oocytes. (b) Schematic of tPTS applied to the P2X2R extracellular domain where a cleavable *faux* TMD (IgK N-term signal sequence, Nx:IgK) was used for the C-construct. (c) Schematic of non-spliced P2X2 fragments expected when +1 extein residue of split intein B (position 76 of P2X2) is mutated to alanine (indicated by red cross). This prevents splicing and favors side reactions, resulting in accumulation of cleavage products (note that this will likely overrepresent the occurrence of the side reactions compared to when splicing-competent split inteins are used). (d) Concentration-response curve (CRC) of reconstituted P2X2 receptors compared to WT. Values displayed as mean +/- SD; n = 4-5). (e) Maximal current size comparison of the tPTS reconstituted receptors using the non-cleavable/cleavable *faux*TMD recorded 3 days after injection of mRNA. No current was observed for the non-spliced constructs (produced by tPTS side reactions, N+X^REC^+C(S76A)). Values displayed as mean +/- SD; n = 6-8. Source data are provided as a Source Data file.

**Fig S12:**
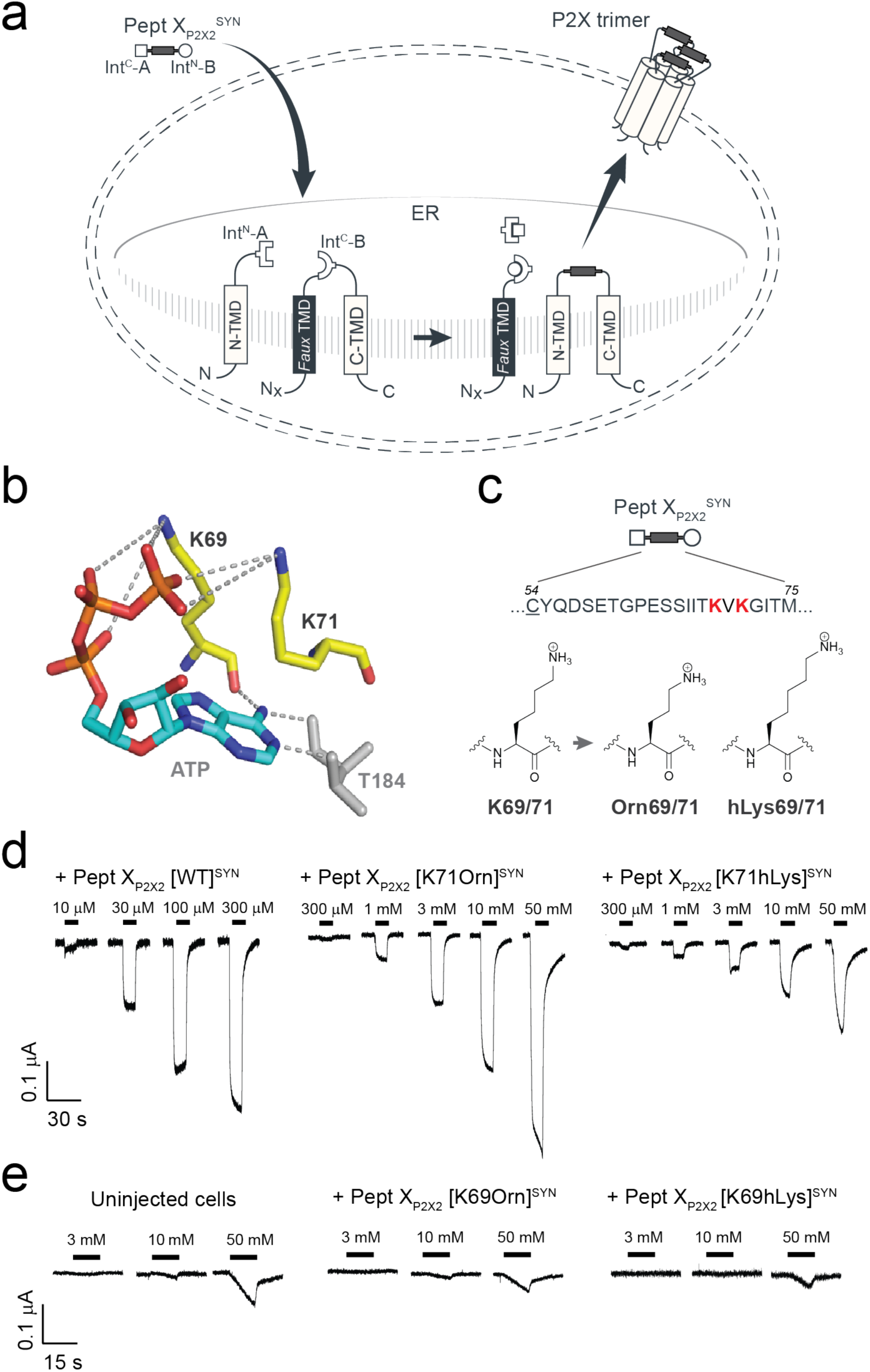
Incorporation of lysine derivatives at the K69 and K71 positions in the ATP binding site of P2X2 using tPTS. (a) Schematic presentation of the strategy to reconstruct full-length P2X2Rs from recombinantly expressed N-/C-terminal fragments (N and C) and a synthetic peptide (Pept X_P2X2_^SYN^) corresponding to amino acids 54 to 75 of the P2X2R extracellular domain in *Xenopus laevis* oocytes. Note that a *faux* transmembrane helix (*faux* TMD) was engineered into the C-terminal fragment to maintain its native membrane topology during protein expression. Inteins A (*Cfa*DnaE) and B (*Ssp*DnaB^M86^) are indicated by square and round symbols, respectively. Peptide X was designed to include a C-terminal ER targeting KDEL signal sequence (not depicted in the schematic presentation), which is excised during the splicing process. (b) Interaction of K69 and K71 (P2X2 residue numbering) with the phosphate tail of ATP (orange) as obtained from the crystal structure of ATP-bound hP2X3R (PDB: 5svk). (c) Sequence of peptide X_P2X2_^SYN^ corresponding to the amino acids replaced in the P2X2 extracellular ATP binding site. Underlined residue indicates S54C mutation that was introduced to optimize the splicing reaction. Lower panel displays the chemical structures of lysine derivatives incorporated at the K69 or K71 position via chemical synthesis of peptide X_P2X2_^SYN^. (d) Representative ATP-induced currents when synthetic peptide with WT sequence (i.e., lysines in positions 69 and 71; left panel) or synthetic peptide X variants with K71 modifications (K71Orn or K71hLys) was incorporated using tPTS. (e) Representative ATP-induced currents when synthetic peptide X variants with K69 modifications (K69Orn or K69hLys) was incorporated using tPTS. Small ATP-induced currents observed only at very high ATP concentrations (>10 mM) were similar to those observed in uninjected cells (left panel).

**Fig S13:**
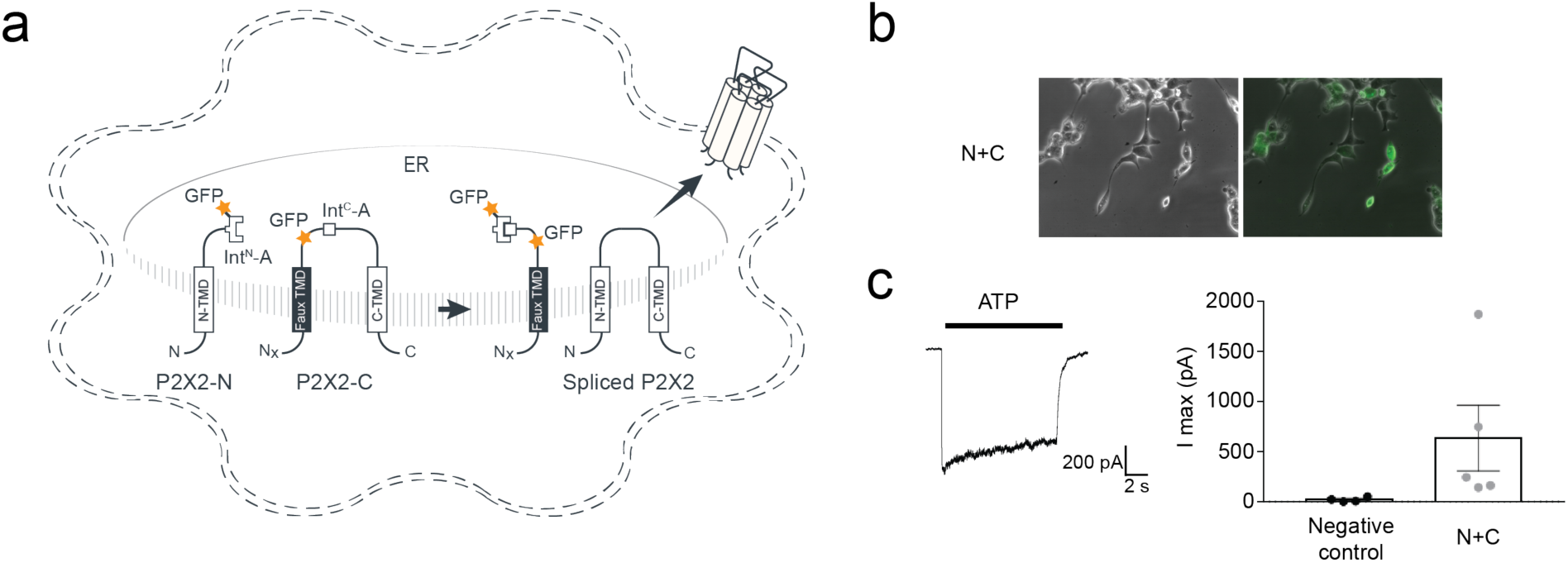
PTS of the P2X2R extracellular domain in HEK293 cells. (a) Schematic of constructs used for single PTS of P2X2Rs in HEK293 cells. Note that a faux transmembrane helix was engineered into the C-terminal fragment to maintain its native membrane topology during protein expression. Intein A (*Cfa*DnaE) is indicated by square symbols and GFP is indicated by an orange star. (b) GFP-fused P2X2-split intein constructs were successfully transfected into HEK293 cells. (c) ATP-induced current recorded during application of 1 mM ATP in HEK293 cells transfected with both P2X2 N and C constructs. Cells transfected with GFP only were used as a negative control. Values are displayed as mean +/- SD; n = 4-5.

**Supplementary Table 1:**
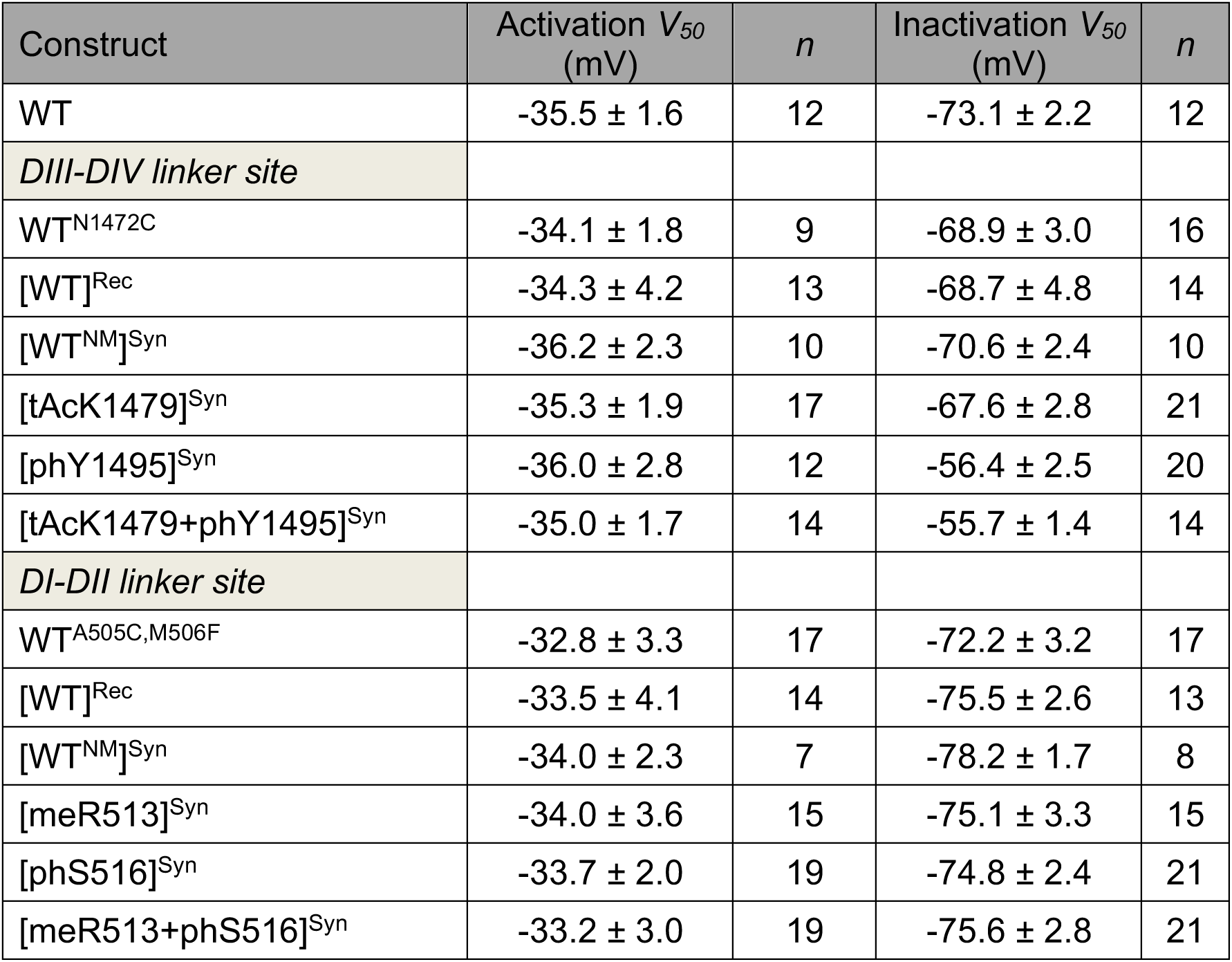
Activation and steady-state inactivation parameters of Na_V_1.5 constructs

## Supplementary methods

### Design of plasmid DNA constructs

Plasmid DNA constructs were designed to encode for the respective protein sequences shown below:

#### hNav1.5 DI-DII linker splicing constructs

N-construct pUNIV - hNav1.5(aa 1-504) - **CfaDnaE_N101_** - *HA tag linker* - *<u>ER retention signal</u>*

MANFLLPRGTSSFRRFTRESLAAIEKRMAEKQARGSTTLQESREGLPEEEAPRPQLDLQA SKKLPDLYGNPPQELIGEPLEDLDPFYSTQKTFIVLNKGKTIFRFSATNALYVLSPFHPIRR AAVKILVHSLFNMLIMCTILTNCVFMAQHDPPPWTKYVEYTFTAIYTFESLVKILARGFCLH AFTFLRDPWNWLDFSVIIMAYTTEFVDLGNVSALRTFRVLRALKTISVISGLKTIVGALIQSV KKLADVMVLTVFCLSVFALIGLQLFMGNLRHKCVRNFTALNGTNGSVEADGLVWESLDLY LSDPENYLLKNGTSDVLLCGNSSDAGTCPEGYRCLKAGENPDHGYTSFDSFAWAFLALF RLMTQDCWERLYQQTLRSAGKIYMIFFMLVIFLGSFYLVNLILAVVAMAYEEQNQATIAET EEKEKRFQEAMEMLKKEHEALTIRGVDTVSRSSLEMSPLAPVNSHERRSKRRKRMSSGT EECGEDRLPKSDSEDGPR**CLSYDTEILTVEYGFLPIGKIVEERIECTVYTVDKNGFVYTQP IAQWHNRGEQEVFEYCLEDGSIIRATKDHKFMTTDGQMLPIDEIFERGLDLKQVDGLP***YP YDVPDYAYPYDVPDY<u>LLDALTLASSRGPLRKRSVAVAKAKPKFSISPDSLS</u>*<u>PRKKFQ</u>**\***

X-construct ‘Rec’

pUNIV - **CfaDnaE_C35_** - hNav1.5(aa 505-527, <u>A505C, M506F</u>) - **SspDnaB^M86^ _N11_**

**VKIISRKSLGTQNVYDIGVEKDHNFLLKNGLVASN**<u>CF</u>NHLSLTRGLSRTSMKPRSSRG**CI SGDSLISLA**

C-construct

pUNIV – *<u>ER retention signal</u>* - *linker* - **SspDnaB^M86^_C143_**- hNav1.5(aa 528-2016)

*<u>MLLDALTLASSRGPLRKRSVAVAKAKPKFSISPDSLS</u>GSAGSAAGSGEF*STGKRVPIKDL LGEKDFEIWAINEQTMKLESAKVSRVFCTGKKLVYTLKTRLGRTIKATANHRFLTIDGWK RLDELSLKEHIALPRKLESSSLQLAPEIEKLPQSDIYWDPIVSITETGVEEVFDLTVPGLRN FVANDIIVHNSIFTFRRRDLG**S**EADFADDENSTAGESESHHTSLLVPWPLRRTSAQGQPS PGTSAPGHALHGKKNSTVDCNGVVSLLGAGDPEATSPGSHLLRPVMLEHPPDTTTPSEE PGGPQMLTSQAPCVDGFEEPGARQRALSAVSVLTSALEELEESRHKCPPCWNRLAQRY LIWECCPLWMSIKQGVKLVVMDPFTDLTITMCIVLNTLFMALEHYNMTSEFEEMLQVGNL VFTGIFTAEMTFKIIALDPYYYFQQGWNIFDSIIVILSLMELGLSRMSNLSVLRSFRLLRVFKL AKSWPTLNTLIKIIGNSVGALGNLTLVLAIIVFIFAVVGMQLFGKNYSELRDSDSGLLPRWH MMDFFHAFLIIFRILCGEWIETMWDCMEVSGQSLCLLVFLLVMVIGNLVVLNLFLALLLSSF SADNLTAPDEDREMNNLQLALARIQRGLRFVKRTTWDFCCGLLRQRPQKPAALAAQGQL PSCIATPYSPPPPETEKVPPTRKETRFEEGEQPGQGTPGDPEPVCVPIAVAESDTDDQEE DEENSLGTEEESSKQQESQPVSGGPEAPPDSRTWSQVSATASSEAEASASQADWRQQ WKAEPQAPGCGETPEDSCSEGSTADMTNTAELLEQIPDLGQDVKDPEDCFTEGCVRRC PCCAVDTTQAPGKVWWRLRKTCYHIVEHSWFETFIIFMILLSSGALAFEDIYLEERKTIKVL LEYADKMFTYVFVLEMLLKWVAYGFKKYFTNAWCWLDFLIVDVSLVSLVANTLGFAEMGP IKSLRTLRALRPLRALSRFEGMRVVVNALVGAIPSIMNVLLVCLIFWLIFSIMGVNLFAGKFG RCINQTEGDLPLNYTIVNNKSQCESLNLTGELYWTKVKVNFDNVGAGYLALLQVATFKGW MDIMYAAVDSRGYEEQPQWEYNLYMYIYFVIFIIFGSFFTLNLFIGVIIDNFNQQKKKLGGQ DIFMTEEQKKYYNAMKKLGSKKPQKPIPRPLNKYQGFIFDIVTKQAFDVTIMFLICLNMVTM MVETDDQSPEKINILAKINLLFVAIFTGECIVKLAALRHYYFTNSWNIFDFVVVILSIVGTVLS DIIQKYFFSPTLFRVIRLARIGRILRLIRGAKGIRTLLFALMMSLPALFNIGLLLFLVMFIYSIFG MANFAYVKWEAGIDDMFNFQTFANSMLCLFQITTSAGWDGLLSPILNTGPPYCDPTLPNS NGSRGDCGSPAVGILFFTTYIIISFLIVVNMYIAIILENFSVATEESTEPLSEDDFDMFYEIWE KFDPEATQFIEYSVLSDFADALSEPLRIAKPNQISLINMDLPMVSGDRIHCMDILFAFTKRVL GESGEMDALKIQMEEKFMAANPSKISYEPITTTLRRKHEEVSAMVIQRAFRRHLLQRSLKH ASFLFRQQAGSGLSEEDAPEREGLIAYVMSENFSRPLGPPSSSSISSTSFPPSYDSVTRA TSDNLQVRGSDYSHSEDLADFPPSPDRDRESIV*

#### Nav1.5 DIII-DIV linker splicing constructs

N-construct

pUNIV - hNav1.5(aa 1-1471) - **CfaDnaE_N101_** – *HA tag linker* - *<u>ER retention signal</u>*

MANFLLPRGTSSFRRFTRESLAAIEKRMAEKQARGSTTLQESREGLPEEEAPRPQLDLQASKKLPDLYGNPPQELIGEPLEDLDPFYSTQKTFIVLNKGKTIFRFSATNALYVLSPFHPIRRAAVKILVHSLFNMLIMCTILTNCVFMAQHDPPPWTKYVEYTFTAIYTFESLVKILARGFCLHAFTFLRDPWNWLDFSVIIMAYTTEFVDLGNVSALRTFRVLRALKTISVISGLKTIVGALIQSVKKLADVMVLTVFCLSVFALIGLQLFMGNLRHKCVRNFTALNGTNGSVEADGLVWESLDLYLSDPENYLLKNGTSDVLLCGNSSDAGTCPEGYRCLKAGENPDHGYTSFDSFAWAFLALFRLMTQDCWERLYQQTLRSAGKIYMIFFMLVIFLGSFYLVNLILAVVAMAYEEQNQATIAETEEKEKRFQEAMEMLKKEHEALTIRGVDTVSRSSLEMSPLAPVNSHERRSKRRKRMSSGTEECGEDRLPKSDSEDGPRAMNHLSLTRGLSRTSMKPRSSRGSIFTFRRRDLGSEADFADDENSTAGESESHHTSLLVPWPLRRTSAQGQPSPGTSAPGHALHGKKNSTVDCNGVVSLLGAGDPEATSPGSHLLRPVMLEHPPDTTTPSEEPGGPQMLTSQAPCVDGFEEPGARQRALSAVSVLTSALEELEESRHKCPPCWNRLAQRYLIWECCPLWMSIKQGVKLVVMDPFTDLTITMCIVLNTLFMALEHYNMTSEFEEMLQVGNLVFTGIFTAEMTFKIIALDPYYYFQQGWNIFDSIIVILSLMELGLSRMSNLSVLRSFRLLRVFKLAKSWPTLNTLIKIIGNSVGALGNLTLVLAIIVFIFAVVGMQLFGKNYSELRDSDSGLLPRWHMMDFFHAFLIIFRILCGEWIETMWDCMEVSGQSLCLLVFLLVMVIGNLVVLNLFLALLLSSFSADNLTAPDEDREMNNLQLALARIQRGLRFVKRTTWDFCCGLLRQRPQKPAALAAQGQLPSCIATPYSPPPPETEKVPPTRKETRFEEGEQPGQGTPGDPEPVCVPIAVAESDTDDQEEDEENSLGTEEESSKQQESQPVSGGPEAPPDSRTWSQVSATASSEAEASASQADWRQQWKAEPQAPGCGETPEDSCSEGSTADMTNTAELLEQIPDLGQDVKDPEDCFTEGCVRRCPCCAVDTTQAPGKVWWRLRKTCYHIVEHSWFETFIIFMILLSSGALAFEDIYLEERKTIKVLLEYADKMFTYVFVLEMLLKWVAYGFKKYFTNAWCWLDFLIVDVSLVSLVANTLGFAEMGPIKSLRTLRALRPLRALSRFEGMRVVVNALVGAIPSIMNVLLVCLIFWLIFSIMGVNLFAGKFGRCINQTEGDLPLNYTIVNNKSQCESLNLTGELYWTKVKVNFDNVGAGYLALLQVATFKGWMDIMYAAVDSRGYEEQPQWEYNLY MYIYFVIFIIFGSFFTLNLFIGVIID**CLSYDTEILTVEYGFLPIGKIVEERIECTVYTVDKNGFVY TQPIAQWHNRGEQEVFEYCLEDGSIIRATKDHKFMTTDGQMLPIDEIFERGLDLKQVDGL P***YPYDVPDYAYPYDVPDY<u>LLDALTLASSRGPLRKRSVAVAKAKPKFSISPDSLS</u>*<u>PRKKFQ</u>**\***

X-construct ‘Rec’

pUNIV - **CfaDnaE_C35_** - hNav1.5(aa 1472-1502, <u>N1472C</u>) - **SspDnaB^M86^ _N11_**

**VKIISRKSLGTQNVYDIGVEKDHNFLLKNGLVASN**<u>C</u>FNQQKKKLGGQDIFMTEEQKKYYN AMKKLG**CISGDSLISLA**

C-construct

pUNIV – *ER retention signal* - *linker* - **SspDnaB^M86^_C143_**- hNav1.5(aa 1503-2016)

*<u>MLLDALTLASSRGPLRKRSVAVAKAKPKFSISPDSLS</u>GSAGSAAGSGEF*STGKRVPIKDL LGEKDFEIWAINEQTMKLESAKVSRVFCTGKKLVYTLKTRLGRTIKATANHRFLTIDGWKRLDELSLKEHIALPRKLESSSLQLAPEIEKLPQSDIYWDPIVSITETGVEEVFDLTVPGLRNFVANDIIVHNSKKPQKPIPRPLNKYQGFIFDIVTKQAFDVTIMFLICLNMVTMMVETDDQSPEKINILAKINLLFVAIFTGECIVKLAALRHYYFTNSWNIFDFVVVILSIVGTVLSDIIQKYFFSPTLFRVIRLARIGRILRLIRGAKGIRTLLFALMMSLPALFNIGLLLFLVMFIYSIFGMANFAYVKWEAGIDDMFNFQTFANSMLCLFQITTSAGWDGLLSPILNTGPPYCDPTLPNSNGSRGDCGSPAVGILFFTTYIIISFLIVVNMYIAIILENFSVATEESTEPLSEDDFDMFYEIWEKFDPEATQFIEYSVLSDFADALSEPLRIAKPNQISLINMDLPMVSGDRIHCMDILFAFTKRVLGESGEMDALKIQMEEKFMAANPSKISYEPITTTLRRKHEEVSAMVIQRAFRRHLLQRSLKHASFLFRQQAGSGLSEEDAPEREGLIAYVMSENFSRPLGPPSSSSISSTSFPPSYDSVTRATSDNLQVRGSDYSHSEDLADFPPSPDRDRESIV*

rP2X2 extracellular site splicing constructs

### Single split intein A (CfaDnaE) splicing constructs

N-construct

pUNIV - rP2X2(aa 1-53) - ***CfaDnaE_N101_*** – *linker* - SEP

MVRRLARGCWSAFWDYETPKVIVVRNRRLGFVHRMVQLLILLYFVWYVFIVQK**CLSYDTE ILTVEYGFLPIGKIVEERIECTVYTVDKNGFVYTQPIAQWHNRGEQEVFEYCLEDGSIIRAT KDHKFMTTDGQMLPIDEIFERGLDLKQVDGLP***GSAGSAAGSGEF*SKGEELFTGVVPILVE LDGDVNGHKFSVSGEGEGDATYGKLTLKFICTTGKLPVPWPTLVTTLTYGVQCFSRYPDHMKRHDFFKSAMPEGYVQERTIFFKDDGNYKTRAEVKFEGDTLVNRIELKGIDFKEDGNILGHKLEYNYNDHQVYIMADKQKNGIKANFKIRHNIEDGGVQLADHYQQNTPIGDGPVLLPDNHYLFTTSTLSKDPNEKRDHMVLLEFVTAAGITHGMDELYK*

C-construct

pUNIV - Nx(rP2X2 aa 1-53) - *linker* - SEP - *linker* - **CfaDnaE_C35_** - rP2X2(aa 54-472, <u>S54C</u>)

MVRRLARGCWSAFWDYETPKVIVVRNRRLGFVHRMVQLLILLYFVWYVFIVQK*GSAGSA AGSGEF*SKGEELFTGVVPILVELDGDVNGHKFSVSGEGEGDATYGKLTLKFICTTGKLPV PWPTLVTTLTYGVQCFSRYPDHMKRHDFFKSAMPEGYVQERTIFFKDDGNYKTRAEVKFEGDTLVNRIELKGIDFKEDGNILGHKLEYNYNDHQVYIMADKQKNGIKANFKIRHNIEDGGVQLADHYQQNTPIGDGPVLLPDNHYLFTTSTLSKDPNEKRDHMVLLEFVTAAGITHGMDE LYK*GSAGSAAGSGEF***VKIISRKSLGTQNVYDIGVEKDHNFLLKNGLVASN**<u>C</u>YQDSETGP ESSIITKVKGITMSEDKVWDVEEYVKPPEGGSVVSIITRIEVTPSQTLGTCPESMRVHSSTCHSDDDCIAGQLDMQGNGIRTGHCVPYYHGDSKTCEVSAWCPVEDGTSDNHFLGKMAPNFTILIKNSIHYPKFKFSKGNIASQKSDYLKHCTFDQDSDPYCPIFRLGFIVEKAGENFTELAHKGGVIGVIINWNCDLDLSESECNPKYSFRRLDPKYDPASSGYNFRFAKYYKINGTTTTRTLIKAYGIRIDVIVHGQAGKFSLIPTIINLATALTSIGVGSFLCDWILLTFMNKNKLYSHKKFDKVRTPKHPSSRWPVTLALVLGQIPPPPSHYSQDQPPSPPSGEGPTLGEGAELPLAVQSPRPCSISALTEQVVDTLGQHMGQRPPVPEPSQQDSTSTDPKGLAQL*

### Double split intein splicing constructs

X-construct ‘Rec’

pUNIV - **CfaDnaE_C35_** - rP2X2(aa 54-75, <u>S54C</u>) - **SspDnaB^M86^ _N11_** – *linker* – *<u>ER targeting signal</u>*

**VKIISRKSLGTQNVYDIGVEKDHNFLLKNGLVASN**<u>C</u>YQDSETGPESSIITKVKGITMCISGD SLISLA*SSGES<u>KDEL</u>*\*

C-construct

pUNIV - Nx(IgK cleavable) – *HA tag linker* - **SspDnaB^M86^_C143_** - rP2X2(aa 76-472) - *myc tag*

METDTLLLWVLLLWVPGSTG^D*YPYDVPDYAGSAGSAAGSGEF*STGKRVPIKDLLGEKD FEIWAINEQTMKLESAKVSRVFCTGKKLVYTLKTRLGRTIKATANHRFLTIDGWKRLDELSLKEHIALPRKLESSSLQLAPEIEKLPQSDIYWDPIVSITETGVEEVFDLTVPGLRNFVANDIIVHNSEDKVWDVEEYVKPPEGGSVVSIITRIEVTPSQTLGTCPESMRVHSSTCHSDDDCIAGQLDMQGNGIRTGHCVPYYHGDSKTCEVSAWCPVEDGTSDNHFLGKMAPNFTILIKNSIHYPKFKFSKGNIASQKSDYLKHCTFDQDSDPYCPIFRLGFIVEKAGENFTELAHKGGVIGVIINWNCDLDLSESECNPKYSFRRLDPKYDPASSGYNFRFAKYYKINGTTTTRTLIKAYGIRIDVIVHGQAGKFSLIPTIINLATALTSIGVGSFLCDWILLTFMNKNKLYSHKKFDKVRTPKHPSSRWPVTLALVLGQIPPPPSHYSQDQPPSPPSGEGPTLGEGAELPLAVQSPRPCSISAL TEQVVDTLGQHMGQRPPVPEPSQQDSTSTDPKGLAQL*EQKLISEEDL*\*

#### eGFP splicing constructs

N-construct

Pcdna3.1 - eGFP(aa 1-64)**- CfaDnaE_N101_**

MVSKGEELFTGVVPILVELDGDVNGHKFSVSGEGEGDATYGKLTLKFICTTGKLPVPWPT LVTTL**CLSYDTEILTVEYGFLPIGKIVEERIECTVYTVDKNGFVYTQPIAQWHNRGEQEVF EYCLEDGSIIRATKDHKFMTTDGQMLPIDEIFERGLDLKQVDGLP\***

X-construct

Pcdna3.1 – *<u>TAT cpp</u>* **–** *linker* **– CfaDnaE_C35_** - eGFP (aa 65**-**85, <u>T65C</u>) **- SspDnaB^M86^ _N11_**

M*<u>GRKKRRQRRRPQ</u>GSAGSAAGSGEF*VKIISRKSLGTQNVYDIGVEKDHNFLLKNGLVA SN<u>C</u>YGVQCFSRYPDHMKQHDFFK**CISGDSLISLA\***

C-construct

Pcdna3.1 **–** <u>HA tag</u> **–** *linker* **- SspDnaB^M86^_C143_**- eGFP (aa 86-238)

M<u>YPYDVPDYA</u>*GSAGSAAGSGEF*STGKRVPIKDLLGEKDFEIWAINEQTMKLESAKVSRV FCTGKKLVYTLKTRLGRTIKATANHRFLTIDGWKRLDELSLKEHIALPRKLESSSLQLAPEIEKLPQSDIYWDPIVSITETGVEEVFDLTVPGLRNFVANDIIVHNSAMPEGYVQERTIFFKDDGNYKTRAEVKFEGDTLVNRIELKGIDFKEDGNILGHKLEYNYNSHNVYIMADKQKNGIKVNFKIRHNIEDGSVQLADHYQQNTPIGDGPVLLPDNHYLSTQSALSKDPNEKRDHMVLLEFVTAAGITLGMDELYK*

### Fluorescence-activated cell sorting (FACS)

HEK293T cells were grown in Dulbecco’s modified Eagle’s Medium (DMEM) (Gibco) supplemented with 10 % Fetal Bovine Serum (Biowest), and incubated at 37 °C with 5 % of CO2.

400.000 cells were seeded in 6-wells plates and incubated for 24 hrs. prior transfection.

DNA coding for three GFP-split intein fusion fragments (N, X and C) was co-transfected in a 1:1:1 ratio using a total of 4.5 µg DNA. To keep the same amount of DNA for each combination pcDNA3.1+ empty vector was co-transfected for the N+C and WT eGFP control experiments. Cells were transfected using 6 µg PEI and incubated for *circa* 44 hrs. The cells were then detached with trypsin-EDTA, spun down and resuspended in PBS containing formaldehyde 37% (1:40) and DAPI 200 µM (1:200). The cell suspension was then passed through a cell-strainer cap and analyzed with BD™ LSR II flow cytometer within 15 min.

### Peptide synthesis and purification General

All reagents and solvents were of analytical grade and used without further purification as obtained from commercial suppliers (Iris, Combi-Blocks, Rapp Polymere, Fluoro Chem, Sigma Aldrich). Anhydrous solvents were purchased from Sigma Aldrich. Reactions were conducted under an atmosphere of nitrogen whenever anhydrous solvents were used. Evaporation of solvents was carried out under reduced pressure at temperatures below 45 °C. Loading of resin during solid phase peptide synthesis was checked spectrophotometrically, quantifying the amount of Fmoc released upon cleavage of a small sample^2^.

Low resolution mass spectra were recorded on a MALDI-TOF Bruker Microflex LT/SH system, and samples were prepared using SA (sinapic acid) matrix dissolved in water– MeCN–TFA (50:50:0.1, v/v/v). The calculated mass reported is the most intense peak (100% relative intensity), predicted with mMass software.

High resolution mass spectra (HR-MS) were recorded on a SOLARIX ESI MALDI from Bruker Daltronik. Samples were dissolved in MeCN–water–FA (50:50:0.1, v/v/v) and were analyzed by ESI. The calculated mass reported is the most intense peak predicted with mMass software (100% relative intensity) in the isotopes pattern, which is compared to the most intense peak experimentally found in the isotopes pattern.

Unless otherwise stated, the amino acids used for solid phase peptide synthesis were:

Fmoc-Ala-OH; Fmoc-Cys(Trt)-OH; Fmoc-Phe-OH; Fmoc-Gly-OH; Fmoc-Ile-OH; Fmoc-Lys(Boc)-OH; Fmoc-Leu-OH; Fmoc-Pro-OH; Fmoc-His(Trt)-OH; Fmoc-Asn(Trt)-OH; Fmoc-Gln(Trt)-OH; Fmoc-Arg(Pbf)-OH; Fmoc-Ser(*^t^*Bu)-OH; Fmoc-Thr(*^t^*Bu)-OH; Fmoc-Tyr(*^t^*Bu)- OH; Fmoc-Asp(*^t^*Bu)-OH; Fmoc-Glu(*^t^*Bu)-OH; Fmoc-Met-OH; Fmoc-Val-OH.

Fmoc-tAcLys-OH was kindly donated by prof. Christian A. Olsen (University of Copenhagen).

Chemical ligations of peptide fragments were monitored by diluting 2.5 μL of ligation mixture in water–MeCN (8:2, v/v, 100 μL). The obtained solution was checked by MALDI-TOF and analytical HPLC. Illustrative chromatograms (λ 210 nm) and MALDI-TOF spectra are shown for every ligation.

### Analytical and preparative chromatography

Analytical reversed-phase HPLC was performed on an Agilent 1100 LC system equipped with a C8 Phenomenex Kinetex column [250 mm × 4.60, 5 μm, 100 Å] and a diode array UV detector, using a gradient and rising eluent II (0.1% TFA in MeCN) in eluent I (water–MeCN– TFA, 95:5:0.1, v/v/v) linearly from 0% to 40% over 40 min, with a flow rate of 1.2 mL/min at 40 °C.

Preparative reversed-phase HPLC was performed on an Agilent 1260 Infinity system equipped with a C18 Phenomenex Luna column [250 mm × 21.2 mm, 5 μm, 100 Å] or a C8 Phenomenex Luna column [250 mm × 21.2 mm, 5 μm, 100 Å] and a diode array UV detector, using a gradient of eluent I (water–MeCN–TFA, 95:5:0.1, v/v/v) and eluent II (0.1% TFA in MeCN) as specified for each compound, with a flow rate of 20 mL/min.

### Loading of 2-chlorotrityl chloride polystyrene resin

2-Chlorotrityl chloride resin (250 mg, 0.35 mmol) was transferred in a polypropylene syringe equipped with a fritted disk and swollen in anhydrous CH_2_Cl_2_ for 45 min, followed by washing with anhydrous CH_2_Cl_2_ (2×). *i*-Pr_2_NEt (61 µL, 0.35 mmol, 1.0 equiv) was added to a suspension of Fmoc-Ala-OH (44 mg, 0.14 mmol, 0.4 equiv) in anhydrous CH_2_Cl_2_ (1.5 mL) and the obtained solution was added to the resin. The suspension was agitated for 90 min, after which it was washed with DMF (4×) and CH_2_Cl_2_ (4×). After loading determination, the unreacted sites on resin were capped by incubating the resin with a mixture of CH_2_Cl_2_– MeOH–*i*-Pr_2_NEt (1.7:0.25:0.12, v/v/v, 2.1 mL) for 60 min, followed by washings with CH_2_Cl_2_ (4×).

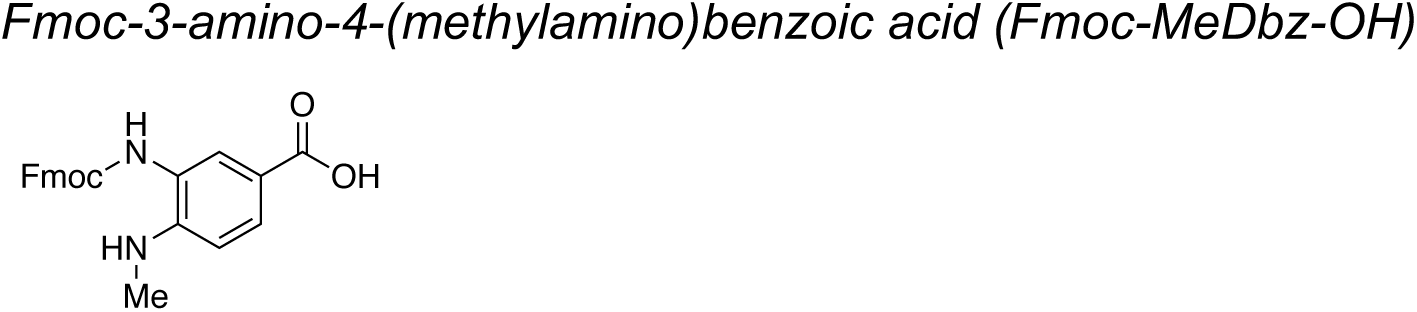

Fmoc-MeDbz-OH was synthesized essentially as previously described^3–5^.

Briefly, 4-fluoro-3-nitrobenzoic acid was dissolved in methanol, followed by MeNH_2_ (40% solution in water, 10 equiv). The reaction mixture turned bright orange and was stirred at room temperature for 20 h, after which the reaction was poured into water. The obtained solution was cooled with an ice bath and acidified with conc. HCl. The resulting bright yellow precipitate was isolated by filtration, washed with cold water and then dried under high vacuum overnight to give 4-methylamine-3-nitrobenzoic acid as a yellow solid.

4-Methylamine-3-nitrobenzoic acid obtained in the previous step was hydrogenated over Pd/C (10% wt) in methanol at atmospheric pressure and at room temperature. The reaction mixture was stirred overnight, during which it turned black. The catalyst was removed by filtration through Celite^®^ and the clear, black filtrate was evaporated under reduced pressure to obtain 3-amino-4-(methylamino)benzoic acid as a black solid.

3-Amino-4-(methylamino)benzoic acid obtained in the previous step was suspended in a mixture of MeCN–water (1:1, v/v). Upon addition of *i*-Pr_2_NEt (0.95 equiv) the reaction mixture turned into a black solution. Fmoc chloride (0.90 equiv) was dissolved in MeCN and was added dropwise to the reaction mixture at room temperature. Upon complete addition of Fmoc chloride, the reaction mixture was stirred for further 45 min, after which the MeCN was evaporated under reduced pressure. The obtained slurry was filtered and the isolated solid was washed several times with cold water and cold MeCN. Drying overnight under high vacuum gave Fmoc-3-amino-4-(methylamino)benzoic acid (Fmoc-MeDbz-OH) as a grey solid.

### Fmoc-MeDbz-Gly PHB TentaGel resin

PHB TentaGel resin (2.0 g, 0.4 mmol) was transferred in a polypropylene syringe equipped with a fritted disk and swollen in CH_2_Cl_2_ for 30 min, followed by washing with anhydrous CH_2_Cl_2_ (2×). In parallel, *N*-methylimidazole (120 µL, 1.5 mmol, 3.75 equiv) was added to a solution of Fmoc-Gly-OH (595 mg, 2 mmol, 5.0 equiv) in anhydrous DMF–CH_2_Cl_2_ (7:1, v/v, 8 mL), followed by MSNT (593 mg, 2 mmol, 5.0 equiv). The obtained solution was added to the resin and the suspension was agitated at room temperature for two hours. The resin was then washed with DMF (3×) and CH_2_Cl_2_ (3×). The loading procedure was repeated once. The unreacted sites were capped *via* treatment with a solution of acetic anhydride (151 µL, 1.6 mmol, 4.0 equiv to original resin loading) and *i*-Pr_2_NEt (418 µL, 2.4 mmol, 6.0 equiv to original resin loading) in CH_2_Cl_2_ (8 mL) for one hour. The resin was then washed with CH_2_Cl_2_ (5×) and Fmoc-deprotected *via* treatment with piperidine–DMF (1:4, v/v, 12 mL) for 2 min, followed by a second treatment for 15 min, after which the resin was washed with DMF (5×).

Fmoc-MeDbz-OH (388 mg, 1.0 mmol, 2.5 equiv) was dissolved in DMF (9 mL), followed by HATU (380 mg, 1.0 mmol, 2.5 equiv) and *i*-Pr_2_NEt (348 µL, 2.0 mmol, 5.0 equiv). The obtained solution was added to the resin and the suspension was agitated at room temperature for 2 hours. The resin was then washed with DMF (3×) and CH_2_Cl_2_ (3×) and the loading procedure repeated once. The resin was then dried under vacuum and the loading determined by Fmoc deprotection.

### SPPS general protocols

Automated peptide synthesis was carried out on a Biotage Syro Wave™ peptide synthesizer using standard Fmoc/*^t^*Bu SPPS chemistry. If not stated differently, SPPS was performed on 0.02 mmol scale using either MeDbz-Gly PHB TentaGel resin or preloaded trityl TentaGel resins (Rapp Polymere). Fmoc deprotection was performed in two stages: piperidine–DMF– formic acid (25:75:0.95, v/v/v) for 3 min, followed by a second treatment for 12 min. The deprotection step was followed by washings with DMF (5×1 min).

Coupling reactions were performed as double couplings using Fmoc-Xaa-OH (6.0 equiv to the resin loading, 0.5 M, dissolved in DMF), HCTU (6.0 equiv, 0.48 M, dissolved in DMF) and *i*-Pr_2_NEt (12 equiv, 2.0 M, dissolved in NMP) for 40 min for each coupling (final concentration of Fmoc-Xaa-OH and HCTU = 0.15 M). Couplings reactions for non-standard Fmoc-protected amino acids were performed as outlined for each peptide (see below). General cleavage and deprotection of the peptides was performed by incubating the resin, if not stated differently, with a mixture of TFA–DODT–TIPS (94:3.3:2.7, v/v/v) for 60–90 min. Upon full deprotection (monitored by MALDI-TOF), the reaction mixture was concentrated under a stream of nitrogen and the crude peptide was precipitated by addition of cold diethyl ether. The solid was spun down, washed with cold diethyl ether (2×) and subjected to preparative HPLC purification.

### General procedure for thioesterification of peptides from MeDbz-Gly PHB TentaGel resin

After automated peptide elongation, the resin (0.02 mmol, 1.0 equiv) was transferred into a polypropylene syringe equipped with a fritted disk where the resin was washed with CH_2_Cl_2_ (5×). Activation of the MeDbz linker was performed similarly to a previously reported procedure^3, 5^. A solution of 4-nitrophenyl-chloroformate (20 mg, 0.10 mmol, 5.0 equiv) in CH_2_Cl_2_ (1.0 mL) was added to the resin and the suspension was incubated for 30 min, after which the resin was washed with CH_2_Cl_2_ (2×). The procedure was repeated once. The resin was then washed with CH_2_Cl_2_ (5×) and DMF (3×) and a solution of *i*-Pr_2_NEt (87 μL, 0.50 mmol, 25.0 equiv) in DMF (1.0 mL) was added to the resin. After 25 min, the resin was washed with DMF (5×). The procedure was repeated once (this procedure was repeated four times for the Int^C^-A peptide). The resin was then washed with DMF (5×), *i*-Pr_2_NEt in DMF (5%, v/v, 3×) and DMF (5×). To cleave the peptide from the support, the resin was treated with a solution of 3-mercaptopropionic acid ethyl ester (25 μL, 0.20 mmol, 10.0 equiv) and *i*-Pr_2_NEt (35 μL, 0.20 mmol, 10.0 equiv) in DMF (1.5 mL). After overnight incubation, the resin was filtered off and washed twice with DMF (1.0 mL). The combined organic phase was concentrated under reduced pressure and then deprotected, if not stated differently, with a mixture of TFA–DODT–TIPS (94:3.3:2.7, v/v/v) for 60–90 min. Upon full deprotection (monitored by MALDI-TOF), the reaction mixture was concentrated under a stream of nitrogen and the crude peptide was precipitated by addition of cold diethyl ether. The solid was spun down, washed with cold diethyl ether (2×) and subjected to preparative HPLC purification.

### General procedure for thioesterification of peptides from trityl TentaGel resins

After automated peptide elongation, the resin (0.02 mmol, 1.0 equiv) was transferred into a polypropylene syringe equipped with a fritted disk where the resin was washed with CH_2_Cl_2_ (5×). The resin was incubated with a solution of HFIP–CH_2_Cl_2_ (1:4 v/v, 2 mL) for 20 min. The supernatant was collected and the procedure repeated once. The resin was then washed with CH_2_Cl_2_ (2×) and the combined organic fractions were evaporated under reduced pressure to give the protected peptide as an off-white residue. The thioesterification procedure was performed as previously described^6^. Briefly, the protected peptide was dissolved in anhydrous DMF (1.0 mL) and the obtained solution was cooled to *circa* −30 °C. The thiol of interest (30 equiv) was then added, followed by *i*-Pr_2_NEt (5 equiv) and PyBOP (5 equiv). The reaction mixture was stirred at *circa* −30 °C for 3 h, after which it was warmed to room temperature and then concentrated under reduced pressure. The obtained residue was deprotected, if not stated differently, with a mixture of TFA–DODT–TIPS (94:3.3:2.7, v/v/v) for 90 min. Upon full deprotection (monitored by MALDI-TOF), the reaction mixture was concentrated under a stream of nitrogen and the crude peptide was precipitated by addition of cold diethyl ether. The solid was spun down, washed with cold diethyl ether (2×) and subjected to preparative HPLC purification.

### General procedure for reduction of oxidized Met containing peptides

Reduction of oxidized methionine residues was performed similarly to a previously described procedure^7^. Briefly, DODT (65 μL, 0.2 M) was added to a solution of the crude peptide in TFA (2.0 mL for a 20 μmol scale), followed by trimethylsilyl bromide (26.4 μL, 0.1 M). The solution was incubated at room temperature for 20 min, after which it was concentrated under a stream of nitrogen. The crude peptide solid was precipitated by addition of cold diethyl ether. The precipitate was spun down, washed with cold diethyl ether and subsequently subjected to preparative HPLC purification.

Alternatively, the reduction could be performed during peptide deprotection under similar conditions: after incubation of full protected peptide with a mixture of TFA–DODT–TIPS (94:3.3:2.7, v/v/v, 4.0 mL) for 90 min, trimethylsilyl bromide (52.8 μL, final concentration 0.1 M) was added and the mixture further incubated for 20 min.

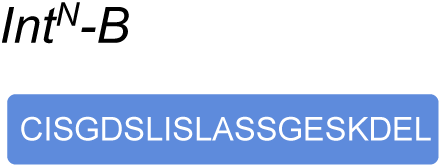

The peptide was synthesized according to the general SPPS protocol outlined above, on Fmoc-Leu PHB TentaGel preloaded resin (0.2 mmol/g). Preparative HPLC purification followed by lyophilization yielded the peptide as a fluffy solid (8.9 mg as a TFA salt; yield 20%).

Prep-HPLC purification conditions (C18 column): 0–10% eluent II in eluent I (5 min gradient) followed by 10–38% eluent II in eluent I (35 min gradient).

Low resolution MS (MALDI-TOF): calc. [C_82_ H_140_N_21_ O_35_ S]^+^ [M + H]^+^: 2010.95 Da; found: 2011.67 Da

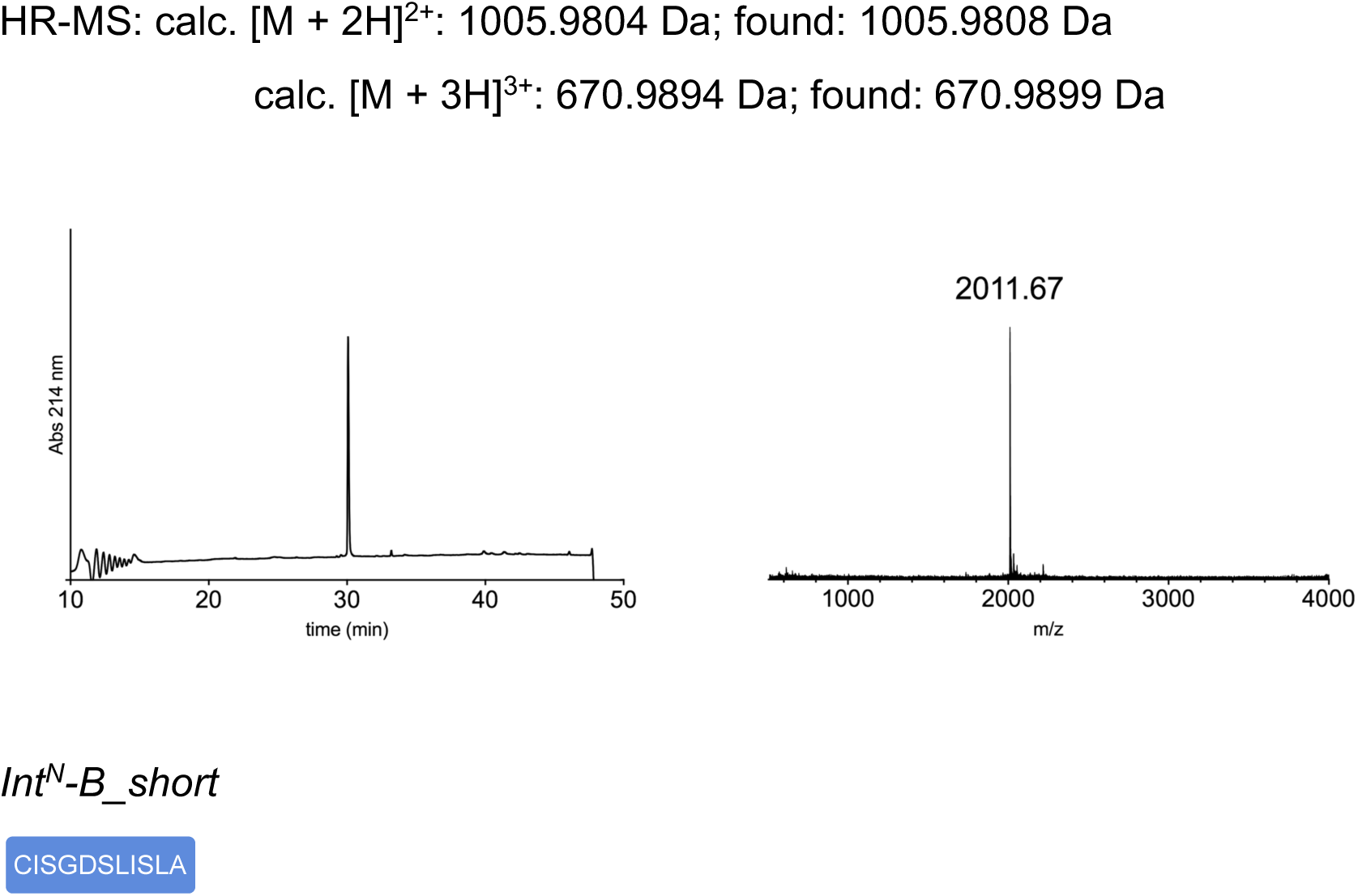

The peptide was synthesized (2 × 20 μmol scale) according to the general SPPS protocol outlined above, on 2-clorotrityl chloride resin (0.39 mmol/g) that was previously loaded with Fmoc-Ala-OH according to the general procedure outlined above. Deprotection and cleavage of the peptide was performed by incubating the resin with a mixture of TFA– DODT–TIPS (95:2.5:2.5, v/v/v) for 60 min. Preparative HPLC purification followed by lyophilization yielded the peptide as a fluffy solid (13.3 mg as a TFA salt; yield 28%).

Prep-HPLC purification conditions (C18 column): 0–35% eluent II in eluent I (35 min gradient).

Low resolution MS (MALDI-TOF): calc. [C_45_H_80_N_11_O_17_S]^+^ [M + H]^+^: 1078.54 Da; found: 1078.06 Da

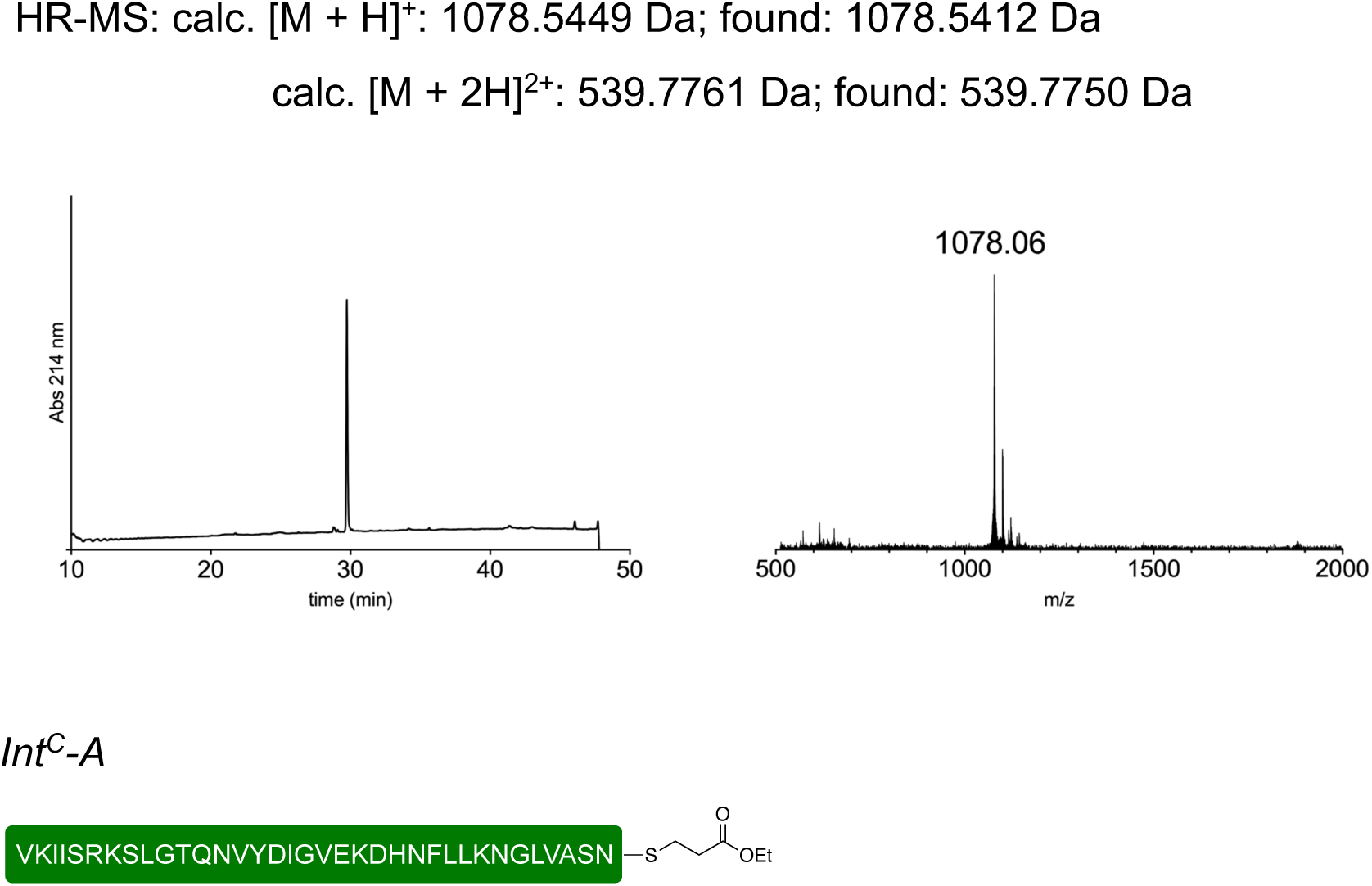

The peptide was synthesized on both preloaded Fmoc-Asn(Trt) trityl TentaGel resin (40 μmol scale, 0.19 mmol/g) and Fmoc-MeDbz-Gly PHB TentaGel resin (20 μmol scale, 0.16 mmol/g) according to the general SPPS protocol outlined above.

When MeDbz-Gly TentaGel resin was used, the loading of the first residue was performed as double coupling using HATU (6.0 equiv) as the coupling reagent and incubating the resin 90 min for each coupling.

Fmoc-DmbGly-OH was incorporated instead of regular Gly at the positions underlined in the sequence (VKIISRKSLGTQNVYDI<u>G</u>VEKDHNFLLKN<u>G</u>LVASN). The coupling was performed as a single coupling similarly to the other coupling steps, but using Fmoc-DmbGly-OH (2.5 equiv), HATU (2.5 equiv) and *i*-Pr_2_NEt (5.0 equiv) and incubating the resin for 90 min. The residue coming after the DmbGly was coupled similarly to the other coupling steps, but using HATU (6.0 equiv) as the coupling reagent.

When MeDbz-Gly TentaGel resin was used, DmbGly was incorporated at the positions underlined in the sequence (VKIISRKSL<u>G</u>TQNVYDI<u>G</u>VEKDHNFLLKN<u>G</u>LVASN).

Boc-Val-OH was coupled to the growing peptide as the N-terminal residue.

After peptide elongation and thioesterification, preparative HPLC purification followed by lyophilization yielded the peptide as a fluffy solid [preloaded Fmoc-Asn(Trt) trityl TentaGel resin (40 μmol): 17.8 mg as a TFA salt; yield 9%. Fmoc-MeDbz-Gly PHB TentaGel resin (20 μmol): 3.4 mg as a TFA salt; yield 3%]

Prep-HPLC purification conditions (C8 column): 0–12% eluent II in eluent I (5 min gradient) followed by 12–33% eluent II in eluent I (35 min gradient).

Low resolution MS (MALDI-TOF): calc. [C_177_H_294_N_49_O_53_S]^+^ [M + H]^+^: 3987.16 Da; found: 3988.34 Da

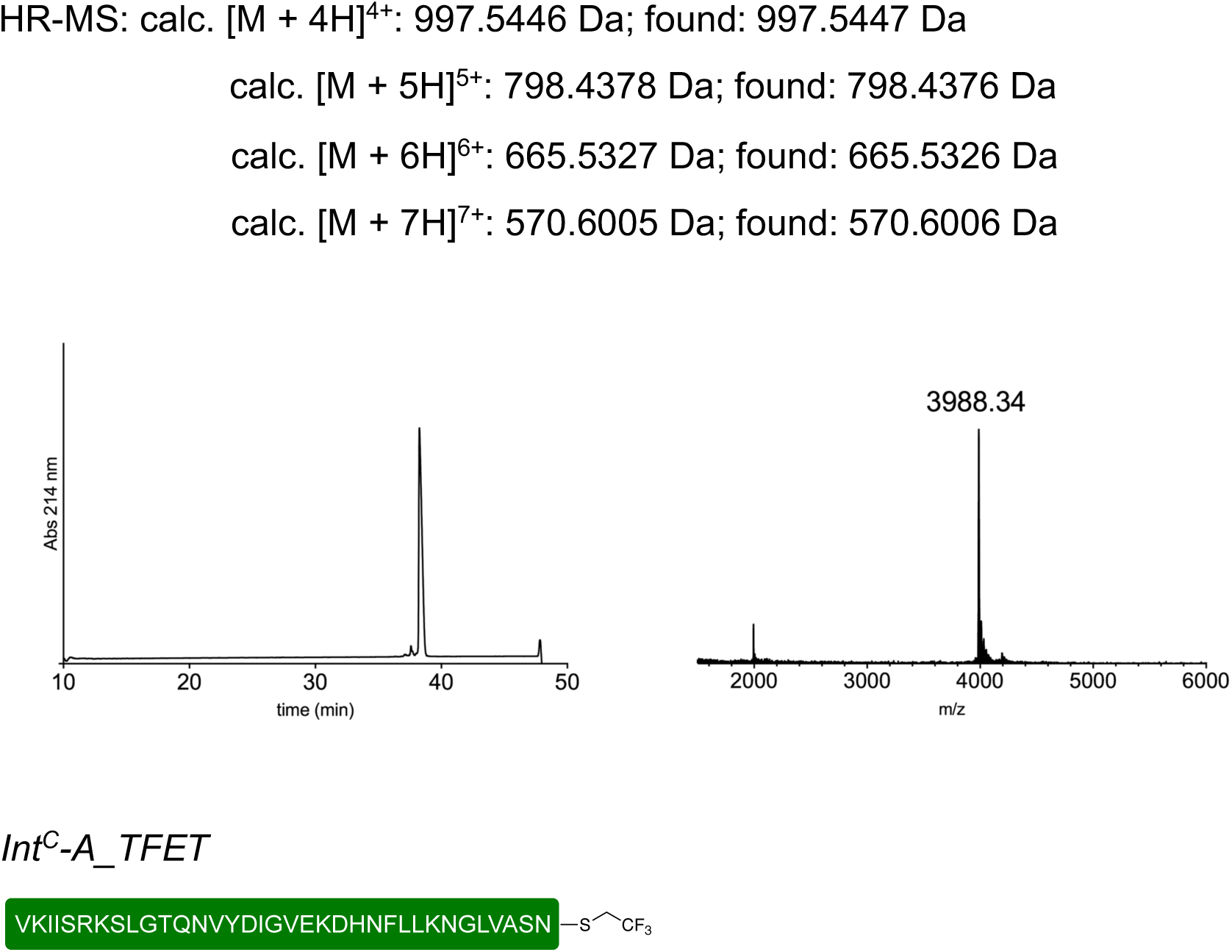

The peptide was synthesized on preloaded Fmoc-Asn(Trt) trityl TentaGel resin (0.19 mmol/g) according to the general SPPS protocol outlined above.

The coupling of the first residue was performed as double coupling using HATU (6.0 equiv) as the coupling reagent for 60 min for each coupling.

Fmoc-DmbGly-OH was incorporated instead of regular Gly at the positions underlined in the sequence (VKIISRKSLGTQNVYDI<u>G</u>VEKDHNFLLKN<u>G</u>LVASN). The coupling was performed as a single coupling similarly to the other coupling steps, but using Fmoc-DmbGly-OH (2.5 equiv), HATU (2.5 equiv) and *i*-Pr_2_NEt (5.0 equiv) and incubating for 90 min. The residue coming after the DmbGly was coupled similarly to the other coupling steps, but using HATU (6.0 equiv) as the coupling reagent.

Boc-Val-OH was coupled to the growing peptide as the N-terminal residue.

After peptide elongation and thioesterification, preparative HPLC purification followed by lyophilization yielded the peptide as a fluffy solid (8.8 mg as a TFA salt; yield 9%).

Prep-HPLC purification conditions (C8 column): 0–10% eluent II in eluent I (5 min gradient) followed by 10–35% eluent II in eluent I (35 min gradient).

Low resolution MS (MALDI-TOF): calc. [C_174_H_287_F_3_N_49_O_51_S]^+^ [M + H]^+^: 3969.11 Da; found: 3969.03 Da

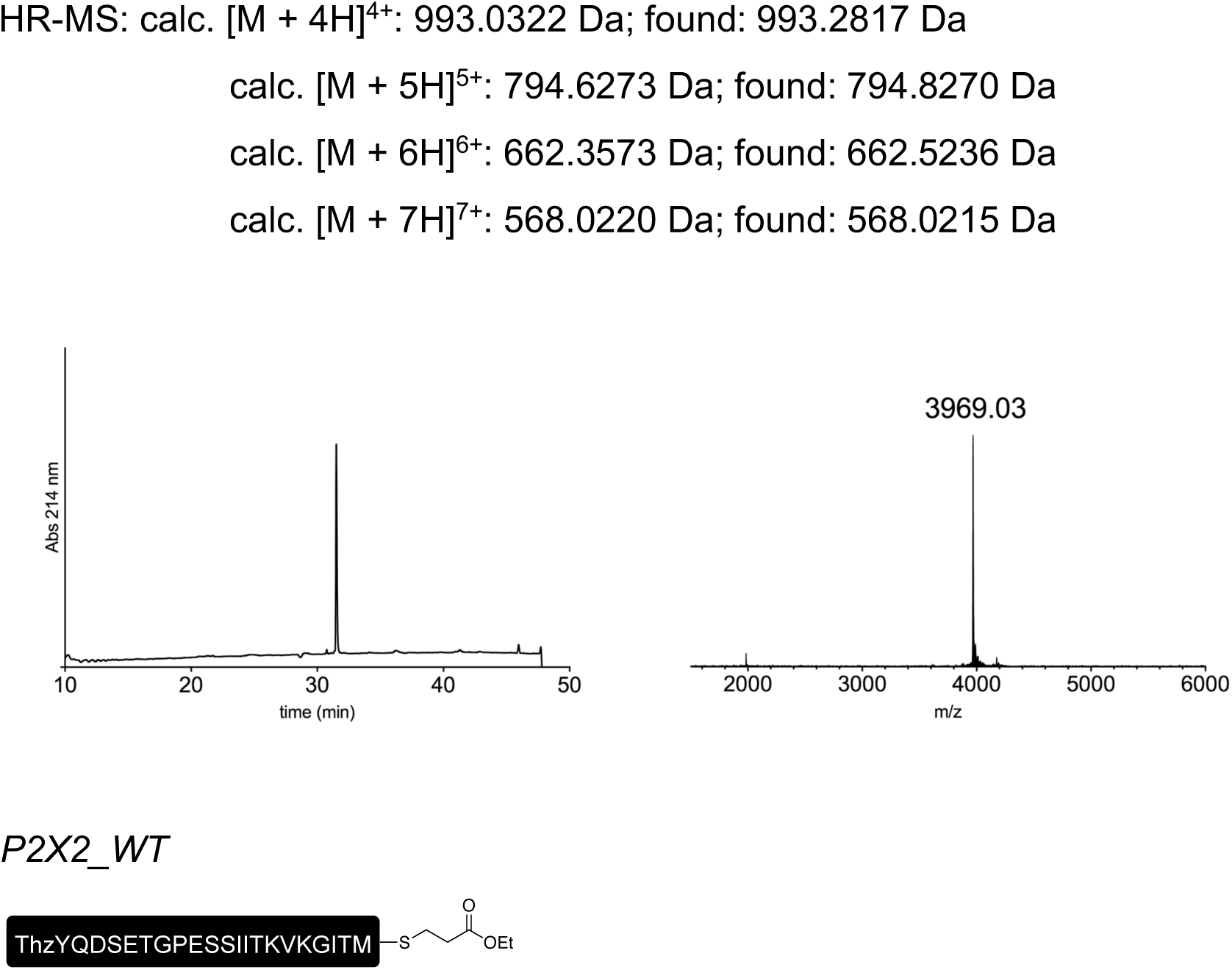

The peptide was synthesized on Fmoc-MeDbz-Gly PHB TentaGel resin (0.16 mmol/g) according to the general SPPS protocol outlined above.

Fmoc-DmbGly-OH was incorporated instead of regular Gly at the position underlined in the sequence (ThzYQDSETGPESSIITKVK<u>G</u>ITM). The coupling was performed as a single coupling similarly to the other coupling steps, but using Fmoc-DmbGly-OH (3.0 equiv), HATU (3.0 equiv) and *i*-Pr_2_NEt (6.0 equiv) and incubating for 90 min. The residue coming after the DmbGly was coupled similarly to the other coupling steps, but using HATU (6.0 equiv) as the coupling reagent.

Boc-Thz-OH was coupled to the growing peptide as the N-terminal residue.

After peptide elongation, thioesterification and deprotection, the crude peptide was reduced as described above. Preparative HPLC purification followed by lyophilization yielded the peptide as a fluffy solid (4.7 mg as a TFA salt; yield 8%).

Prep-HPLC purification conditions (C8 column): 0–10% eluent II in eluent I (5 min gradient) followed by 10–38% eluent II in eluent I (35 min gradient).

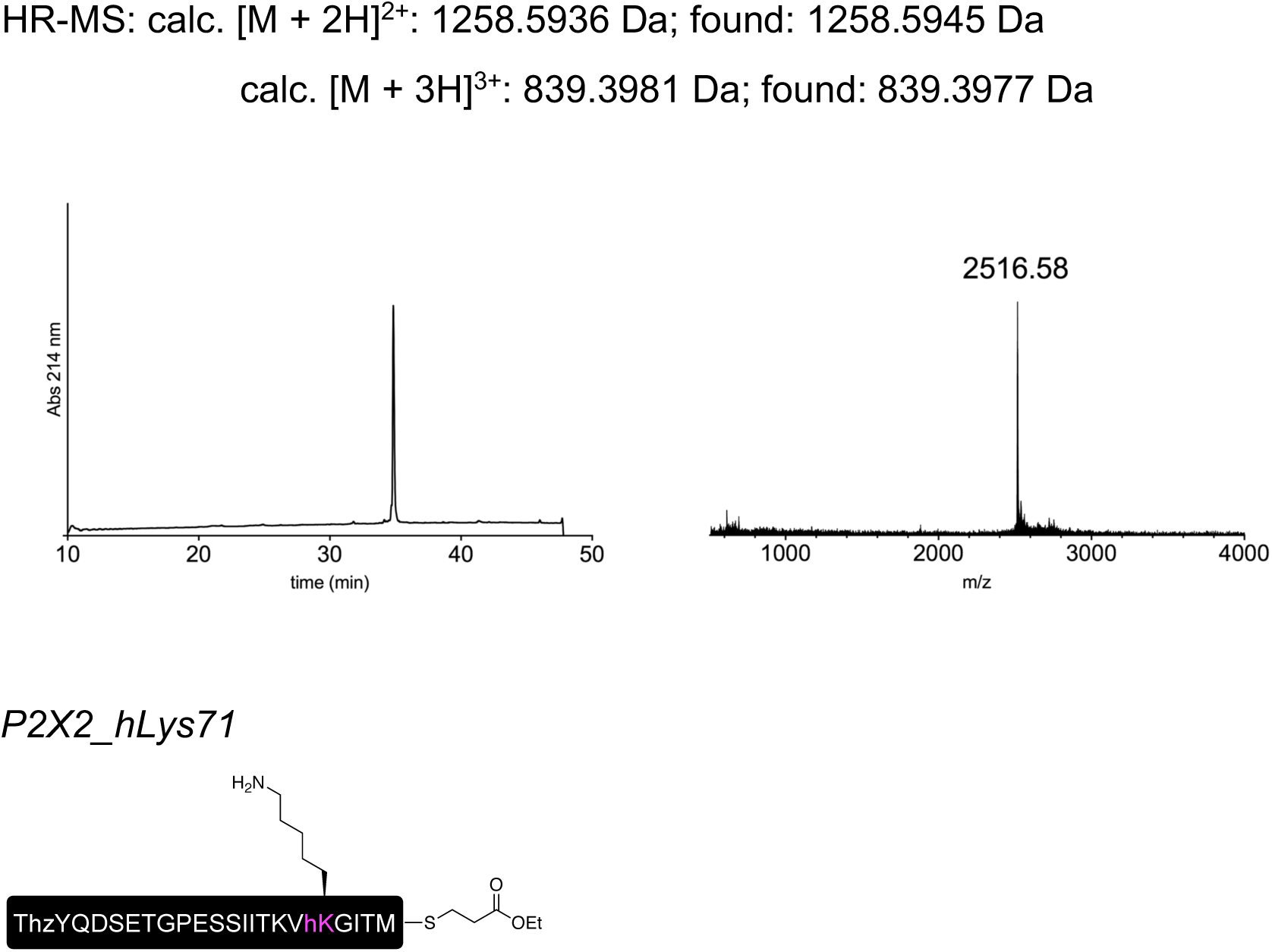

Low resolution MS (MALDI-TOF): calc. [C_107_H_176_N_25_O_38_S_3_]^+^ [M + H]^+^: 2516.18 Da; found: 2516.58 Da.

The peptide was synthesized on Fmoc-MeDbz-Gly PHB TentaGel resin (0.16 mmol/g) according to the general SPPS protocol outlined above.

The loading of the first residue was performed as double coupling using HATU (6.0 equiv) as the coupling reagent for 60 min for each coupling. Fmoc-DmbGly-OH was incorporated instead of regular Gly at the position underlined in the sequence (ThzYQDSETGPESSIITKVhK<u>G</u>ITM). Homolysine (denoted hK or hLys) was incorporated through a Fmoc/Boc protected amino acid building block. The coupling of Fmoc-DmbGly-OH and Fmoc-hLys(Boc)-OH was performed as a single coupling similarly to the other coupling steps, but using the Fmoc protected amino acid (2.5 equiv), HATU (2.5 equiv) and *i*-Pr_2_NEt (5.0 equiv) and incubating for 90 min. The residue coming after the DmbGly was coupled similarly to the other coupling steps, but using HATU (6.0 equiv) as the coupling reagent.

Boc-Thz-OH was coupled to the growing peptide as the N-terminal residue.

After peptide elongation, thioesterification and deprotection, the crude peptide was reduced as described above. Preparative HPLC purification followed by lyophilization yielded the peptide as a fluffy solid (14.4 mg as a TFA salt; yield 12%).

Prep-HPLC purification conditions (C8 column): 0–15% eluent II in eluent I (5 min gradient) followed by 15–35% eluent II in eluent I (32 min gradient).

Low resolution MS (MALDI-TOF): calc. [C_108_H_178_N_25_O_38_S_3_]^+^ [M + H]^+^: 2530.19 Da; found: 2529.58 Da.

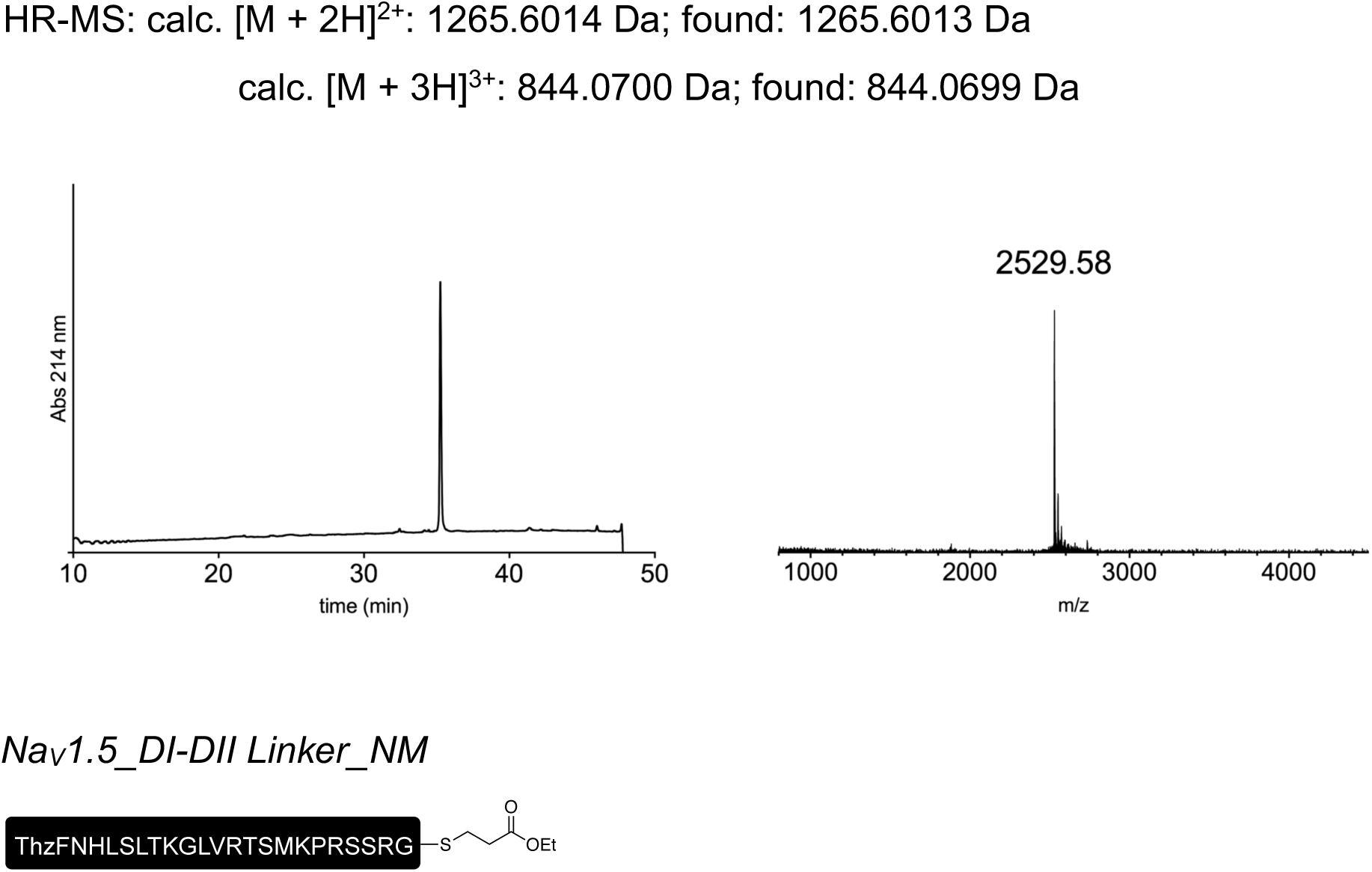

The peptide was synthesized on Fmoc-MeDbz-Gly PHB TentaGel resin (0.16 mmol/g) according to the general SPPS protocol outlined above.

The loading of the first residue was performed as double coupling using HATU (6.0 equiv) as the coupling reagent for 60 min for each coupling.

Boc-Thz-OH was coupled to the growing peptide as the N-terminal residue.

After peptide elongation, thioesterification and deprotection, the crude peptide was reduced as described above. Preparative HPLC purification followed by lyophilization yielded the peptide as a fluffy solid (4.6 mg as a TFA salt; 7% yield).

Prep-HPLC purification conditions (C8 column): 0–10% eluent II in eluent I (5 min gradient) followed by 10–30% eluent II in eluent I (30 min gradient).

Low resolution MS (MALDI-TOF): calc. [C_115_H_196_N_37_O_32_S_3_]^+^ [M + H]^+^: 2704.40 Da; found: 2703.35 Da.

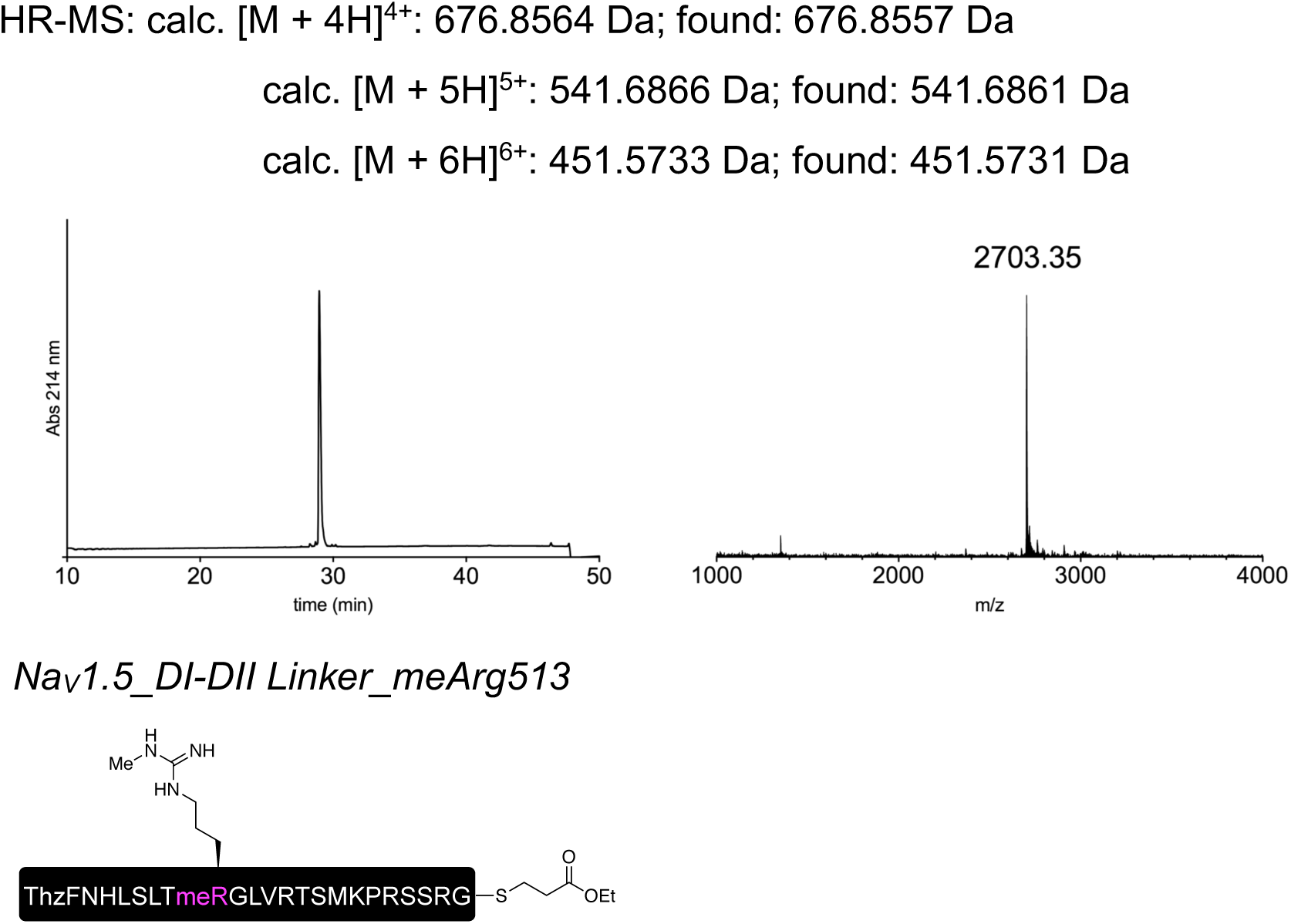

The peptide was synthesized on pre-loaded Fmoc-Gly Trityl TentaGel resin (0.22 mmol/g) according to the general SPPS protocol outlined above.

Methylated arginine (denoted meR or meArg) was incorporated through a Fmoc/Pbf protected amino acid building block. The coupling of Fmoc-Arg(Me,Pbf)-OH was performed as a single coupling similarly to the other coupling steps, but using Fmoc-Arg(Me,Pbf)-OH (2.5 equiv), HATU (2.5 equiv), *i*-Pr_2_NEt (5.0 equiv) and incubating for 2 hours.

Boc-Thz-OH was coupled to the growing peptide as the N-terminal residue.

After peptide elongation, thioesterification and deprotection, the crude peptide was reduced as described above. Preparative HPLC purification followed by lyophilization yielded the peptide as a fluffy solid (7.3 mg as a TFA salt; yield 10%).

Prep-HPLC purification conditions (C8 column): 0–10% eluent II in eluent I (5 min gradient) followed by 10–30% eluent II in eluent I (30 min gradient).

Low resolution MS (MALDI-TOF): calc. [C_116_H_198_N_39_O_32_S_3_]^+^ [M + H]^+^: 2746.42 Da; found: 2746.66 Da.

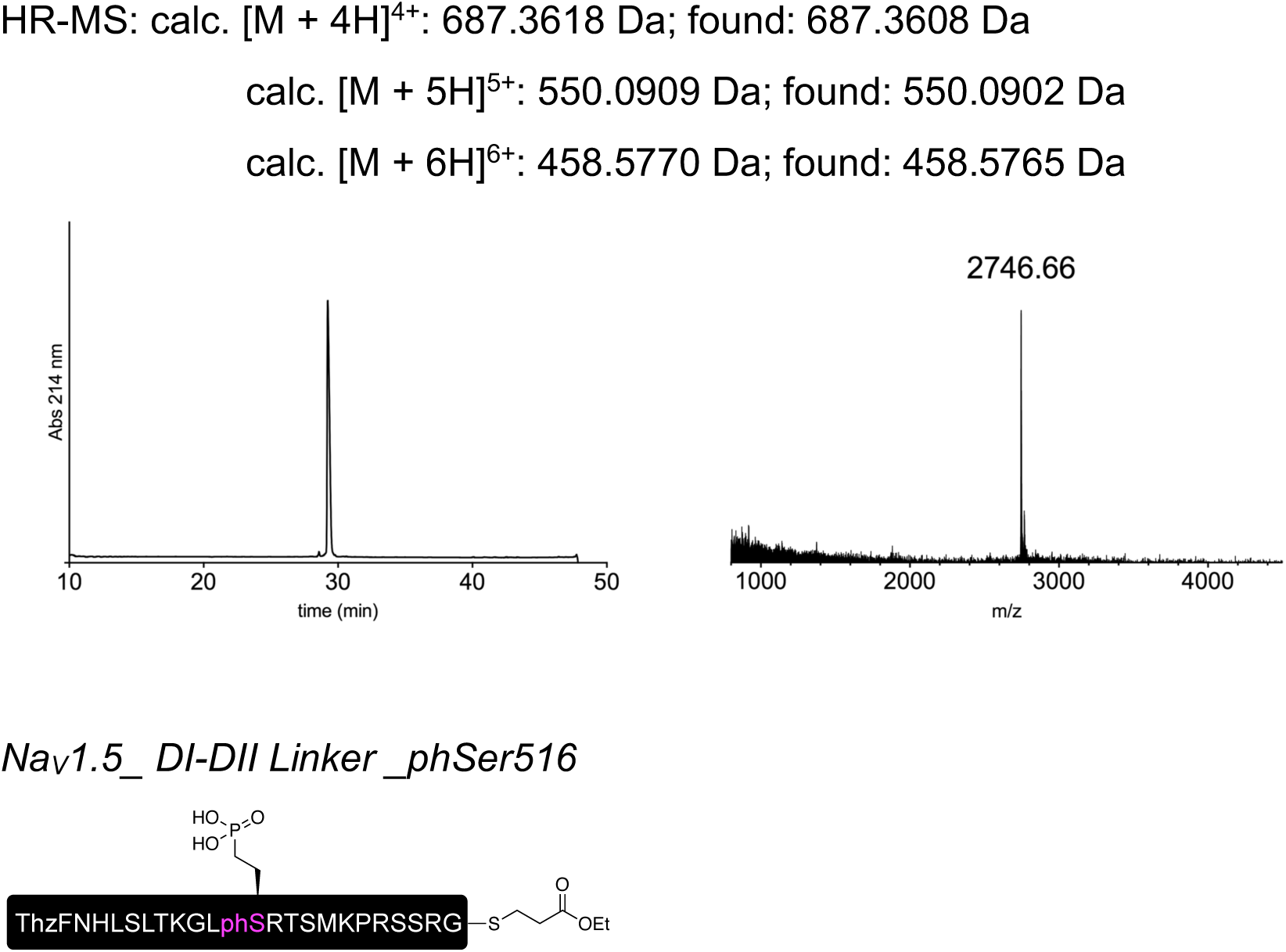

The peptide was synthesized on pre-loaded Fmoc-Gly Trityl TentaGel resin (0.22 mmol/g) according to the general SPPS protocol outlined above.

Phosphonylated serine (denoted phS or phSer) was incorporated through a Fmoc/*^t^*Bu protected amino acid building block. The coupling of Fmoc-Pma(*^t^*Bu)_2_-OH was performed as a single coupling similarly to the other coupling steps, but using Fmoc-Pma(*^t^*Bu)_2_-OH (2.5 equiv), HATU (2.5 equiv) and *i*-Pr_2_NEt (5.0 equiv) and incubating for 2 hours.

Boc-Thz-OH was coupled to the growing peptide as the N-terminal residue.

After peptide elongation, thioesterification and deprotection, the crude peptide was reduced as described above. Preparative HPLC purification followed by lyophilization yielded the peptide as a fluffy solid (7.3 mg as a TFA salt; yield 10%).

Prep-HPLC purification conditions (C8 column): 0–10% eluent II in eluent I (5 min gradient) followed by 10–30% eluent II in eluent I (30 min gradient).

Low resolution MS (MALDI-TOF): calc. [C_114_H_195_N_37_O_35_PS_3_]^+^ [M + H]^+^: 2770.35 Da; found: 2771.07 Da.

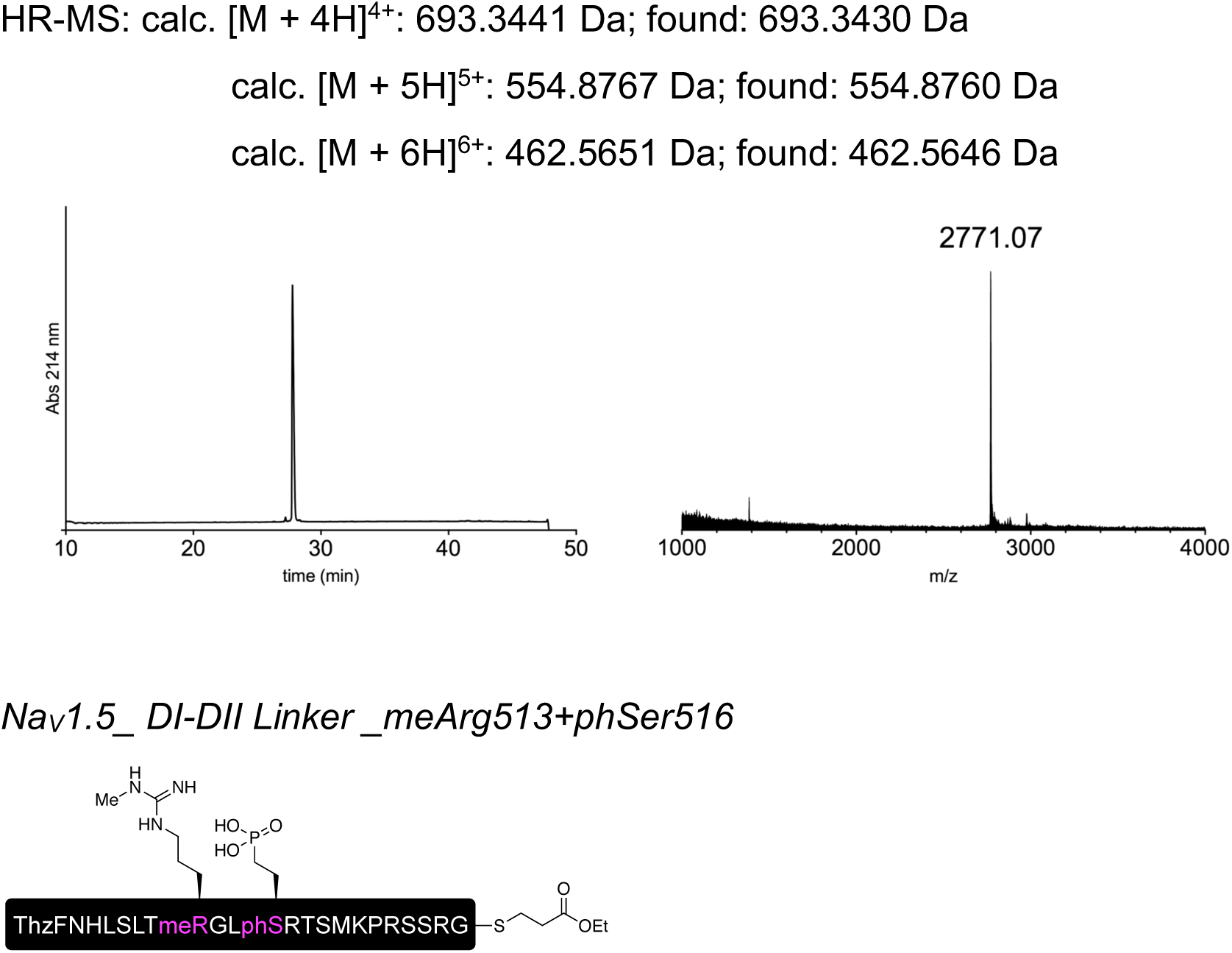

The peptide was synthesized on pre-loaded Fmoc-Gly Trityl TentaGel resin (0.22 mmol/g) according to the general SPPS protocol outlined above.

Phosphonylated serine (denoted phS or phSer) and methylated arginine (denoted meR or meArg) were incorporated through a Fmoc/*^t^*Bu and a Fmoc/Pbf protected amino acid building blocks, respectively. The coupling of Fmoc-Arg(Me,Pbf)-OH and Fmoc-Pma(*^t^*Bu)_2_-

OH was performed as a single coupling similarly to the other coupling steps, but using the Fmoc protected amino acid (2.5 equiv), HATU (2.5 equiv) and *i*-Pr_2_NEt (5.0 equiv) and incubating for 2 hours.

Boc-Thz-OH was coupled to the growing peptide as the N-terminal residue.

After peptide elongation, thioesterification and deprotection, the crude peptide was reduced as described above. Preparative HPLC purification followed by lyophilization yielded the peptide as a fluffy solid (9.0 mg as a TFA salt; yield 12%).

Prep-HPLC purification conditions (C8 column): 0–10% eluent II in eluent I (5 min gradient) followed by 10–30% eluent II in eluent I (30 min gradient).

Low resolution MS (MALDI-TOF): calc. [C_115_H_197_N_39_O_35_PS_3_]^+^ [M + H]^+^: 2812.38 Da; found: 2812.11 Da.

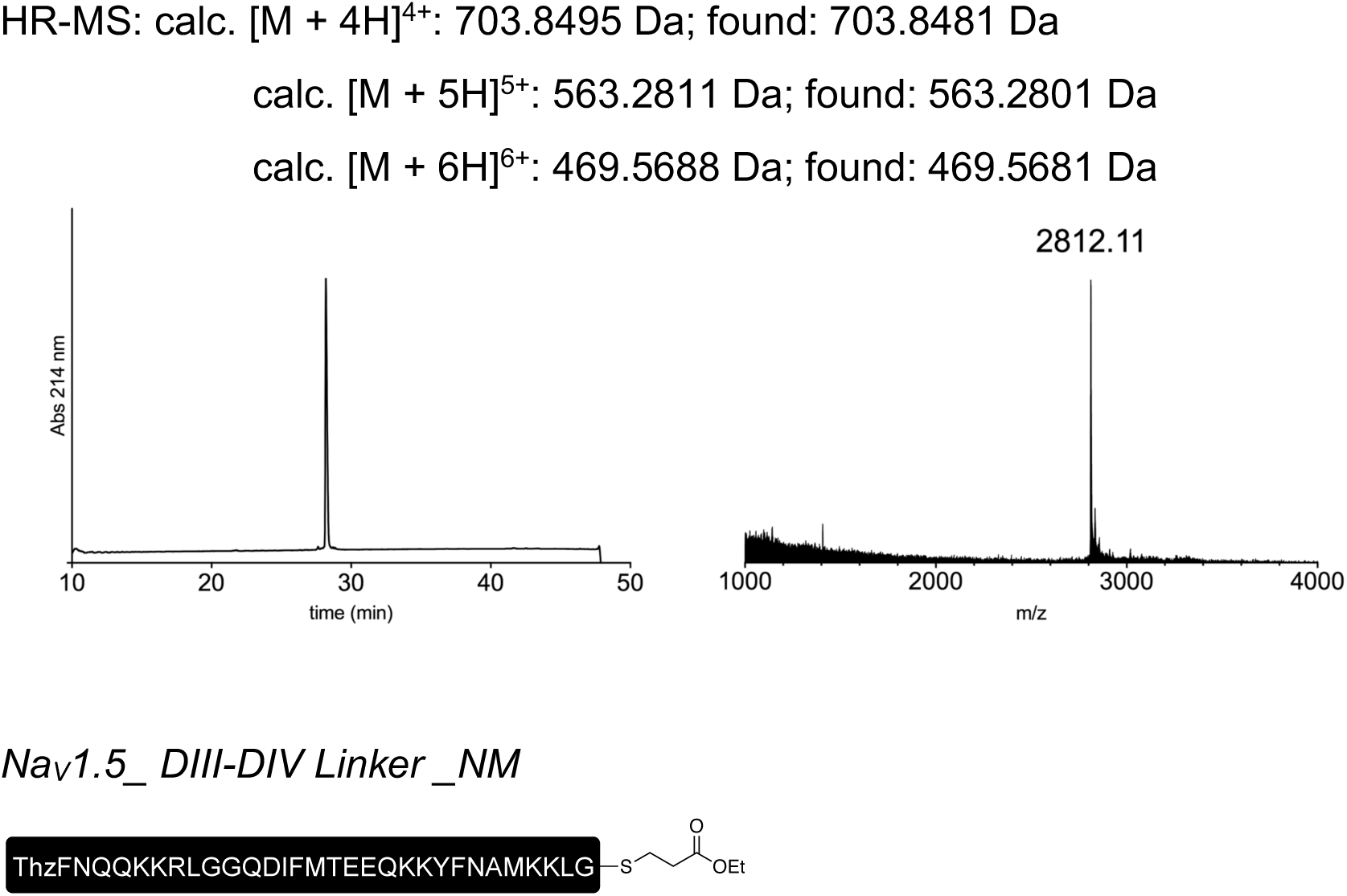

The peptide was synthesized on Fmoc-MeDbz-Gly PHB TentaGel resin (0.16 mmol/g) according to the general SPPS protocol outlined above.

The loading of the first residue was performed as double coupling using HATU (6.0 equiv) as the coupling reagent and incubating for 60 min (first coupling) + 90 min (second coupling).

Fmoc-DmbGly-OH was incorporated instead of regular Gly at the position underlined in the sequence (ThzFNQQKKRL<u>G</u>GQDIFMTEEQKKYFNAMKKLG). The coupling was performed as a single coupling similarly to the other coupling steps, but using Fmoc-DmbGly-OH (3.0 equiv), HATU (3.0 equiv) and *i*-Pr_2_NEt (6.0 equiv) and incubating for 90 min. The residue coming after the DmbGly was coupled similarly to the other coupling steps, but using HATU (6.0 equiv) as the coupling reagent.

Boc-Thz-OH was coupled to the growing peptide as the N-terminal residue.

After peptide elongation, thioesterification and deprotection, the crude peptide was subjected to preparative HPLC purification. Lyophilization of the collected fractions yielded the peptide as a fluffy solid (4.0 mg as a TFA salt; yield 4%).

Prep-HPLC purification conditions (C8 column): 0–15% eluent II in eluent I (5 min gradient) followed by 15–36% eluent II in eluent I (30 min gradient).

Low resolution MS (MALDI-TOF): calc. [C_170_H_271_N_46_O_47_S_4_]^+^ [M + H]^+^: 3837.91 Da; found: 3839.30 Da.

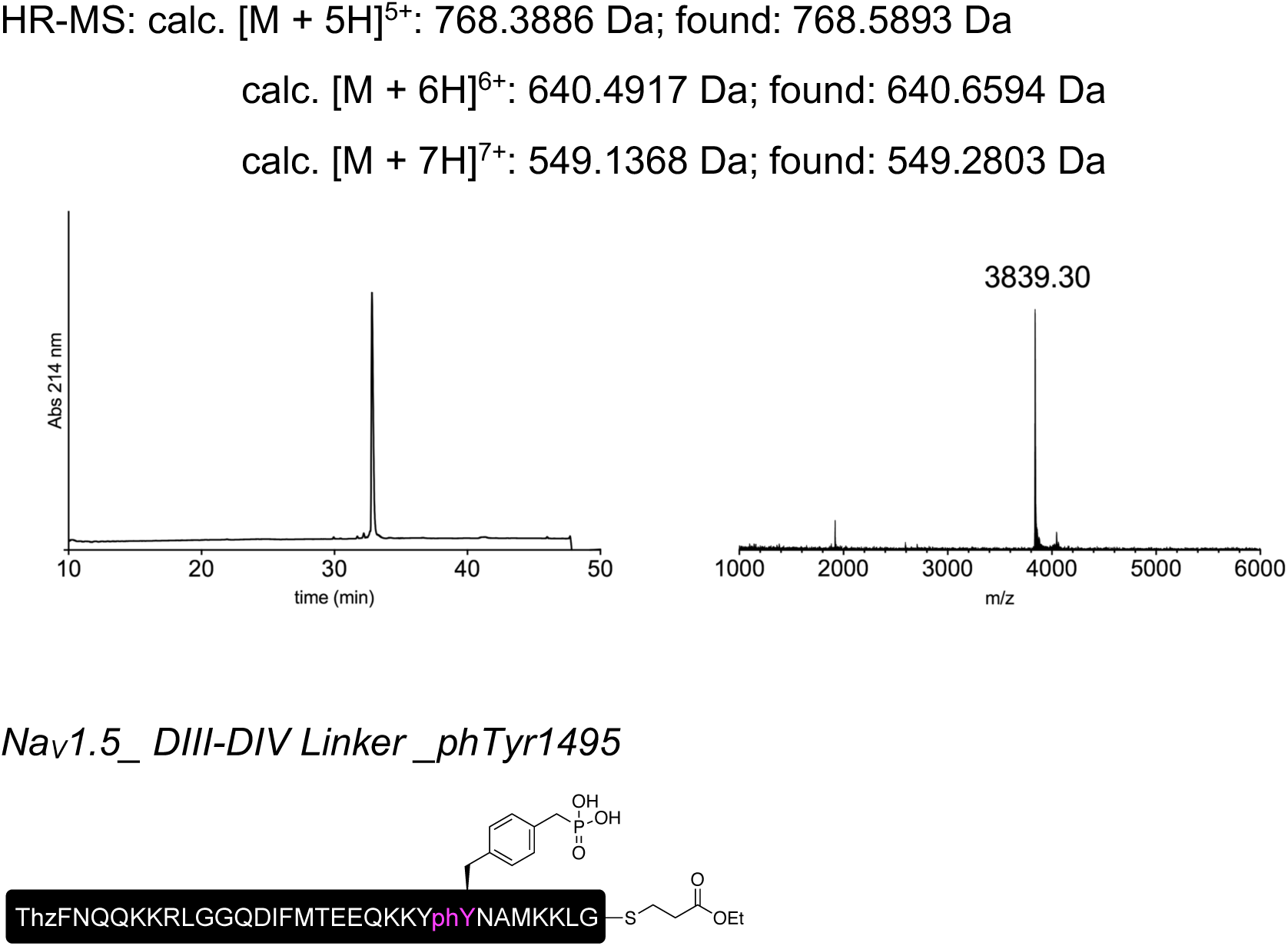

The peptide was synthesized on pre-loaded Fmoc-Gly Trityl TentaGel resin (0.22 mmol/g) according to the general SPPS protocol outlined above.

The loading of the first residue was performed as double coupling using HATU (6.0 equiv) as the coupling reagent and incubating for 60 min.

Phosphonylated tyrosine (denoted phY or phTyr) was incorporated through a Fmoc/*^t^*Bu protected amino acid building block. The coupling of Fmoc-Pmp(*^t^*Bu)_2_-OH was performed as a single coupling similarly to the other coupling steps, but using Fmoc-Pmp(*^t^*Bu)_2_-OH (2.5 equiv), HATU (2.5 equiv) and *i*-Pr_2_NEt (5.0 equiv) and incubating for 2 hours.

Boc-Thz-OH was coupled to the growing peptide as the N-terminal residue.

After peptide elongation, thioesterification and deprotection, the crude peptide was reduced as described above. Preparative HPLC purification followed by lyophilization yielded the peptide as a fluffy solid (9.0 mg as a TFA salt; yield 9%).

Prep-HPLC purification conditions (C8 column): 0–15% eluent II in eluent I (5 min gradient) followed by 15–36% eluent II in eluent I (30 min gradient).

Low resolution MS (MALDI-TOF): calc. [C_171_H_274_N_46_O_50_PS_4_]^+^ [M + H]^+^: 3931.90 Da; found: 3932.64 Da.

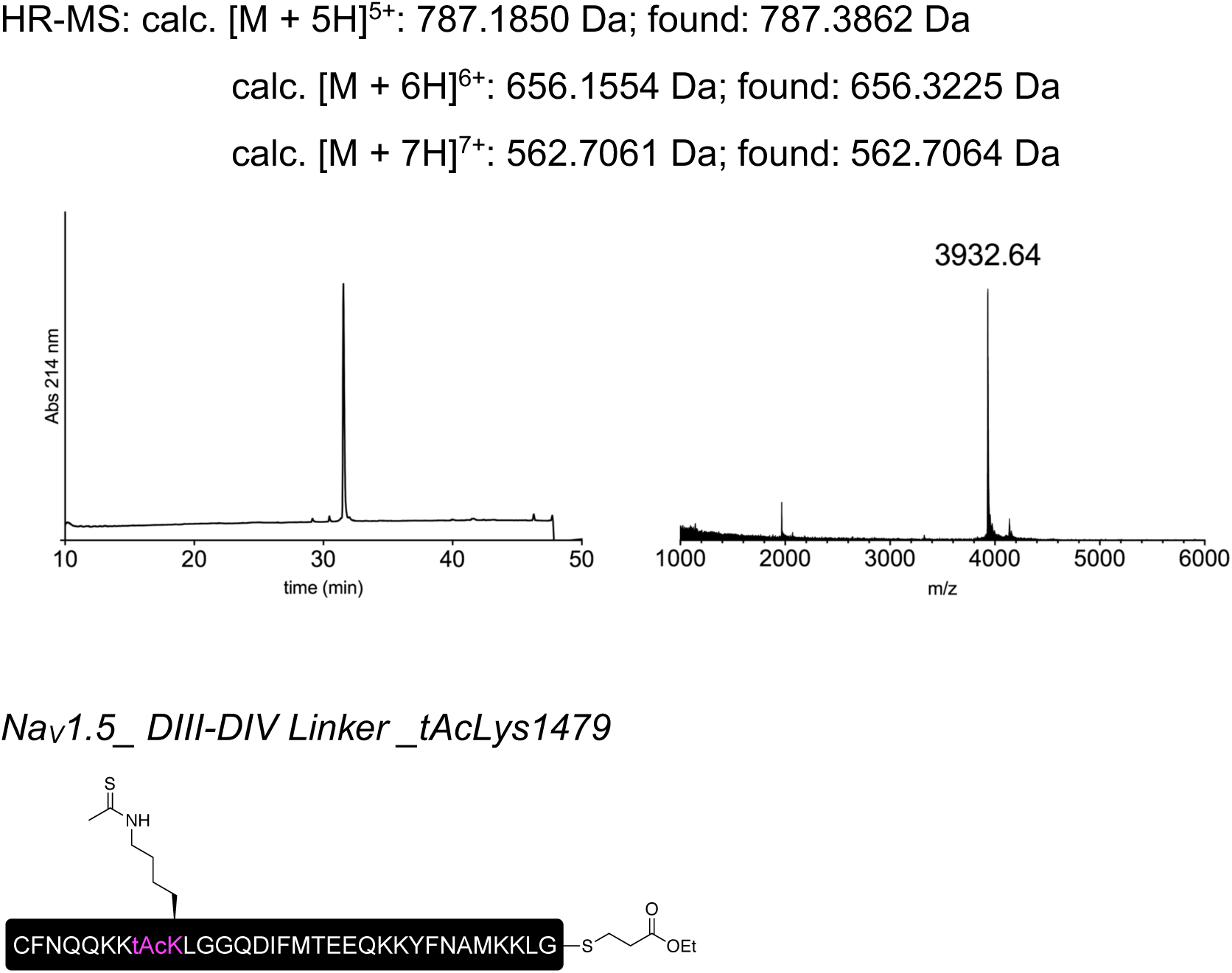

The peptide was synthesized on pre-loaded Fmoc-Gly Trityl TentaGel resin (0.21 mmol/g) according to the general SPPS protocol outlined above.

The loading of the first residue was performed as double coupling using HATU (6.0 equiv) as the coupling reagent and incubating for 60 min.

Thioacetylated lysine (denoted tAcK or tAcLys) was incorporated through a Fmoc protected amino acid building block. The coupling of Fmoc-tAcLys-OH was performed as a single coupling similarly to the other coupling steps, but by using Fmoc-tAcLys-OH (2.5 equiv), HATU (2.5 equiv) and *i*-Pr_2_NEt (5.0 equiv) and incubating for 2 hours.

Boc-Cys(Trt)-OH was coupled to the growing peptide as the N-terminal residue.

After peptide elongation, thioesterification and deprotection, the crude peptide was reduced as described above. Preparative HPLC purification followed by lyophilization yielded the peptide as a fluffy solid (7.0 mg as a TFA salt; yield 7%).

Prep-HPLC purification conditions (C8 column): 0–15% eluent II in eluent I (5 min gradient) followed by 15–35% eluent II in eluent I (30 min gradient).

Low resolution MS (MALDI-TOF): calc. [C_171_H_273_N_44_O_47_S_5_]^+^ [M + H]^+^: 3855.90 Da; found: 3856.23 Da.

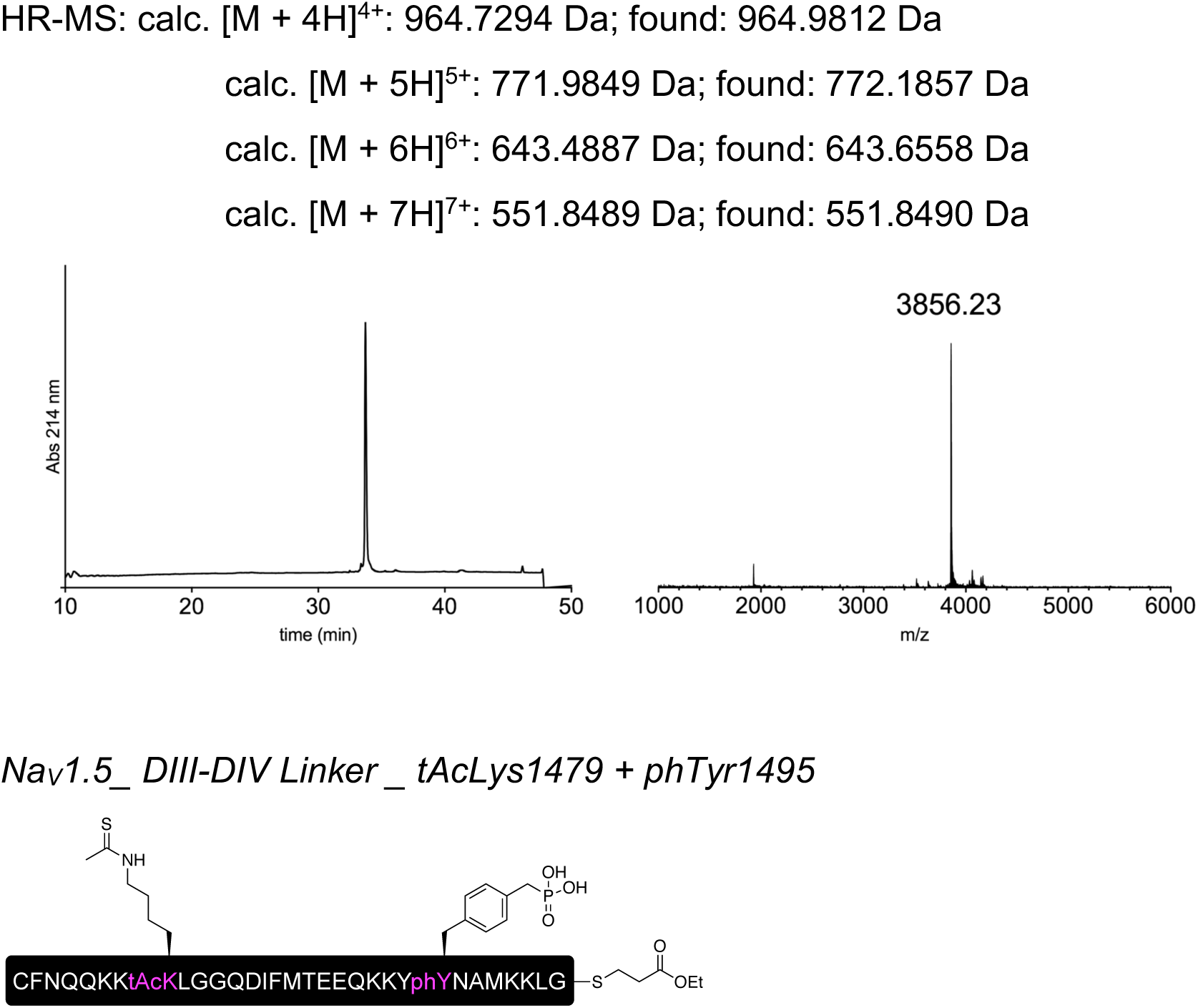

The peptide was synthesized on pre-loaded Fmoc-Gly Trityl TentaGel resin (0.21 mmol/g) according to the general SPPS protocol outlined above.

Phosphonylated tyrosine (denoted phY or phTyr) and thioacetylated lysine (denoted tAcK or tAcLys) were incorporated through a Fmoc/*^t^*Bu and a Fmoc protected amino acid building blocks, respectively. The coupling of Fmoc-Pmp(*^t^*Bu)_2_-OH and Fmoc-tAcLys-OH was performed as a single coupling similarly to the other coupling steps, but by using the Fmoc protected amino acid (2.5 equiv), HATU (2.5 equiv) and *i*-Pr_2_NEt (5.0 equiv) and incubating for 2 hours.

Boc-Cys(Trt)-OH was coupled to the growing peptide as the N-terminal residue.

After peptide elongation, thioesterification and deprotection, the crude peptide was reduced as described above. Preparative HPLC purification followed by lyophilization yielded the peptide as a fluffy solid (4.3 mg as a TFA salt; yield 4%).

Prep-HPLC purification conditions (C8 column): 0–15% eluent II in eluent I (5 min gradient) followed by 15–35% eluent II in eluent I (30 min gradient).

Low resolution MS (MALDI-TOF): calc. [C_172_H_276_N_44_O_50_PS_5_]^+^ [M + H]^+^: 3949.88 Da; found: 3951.86 Da.

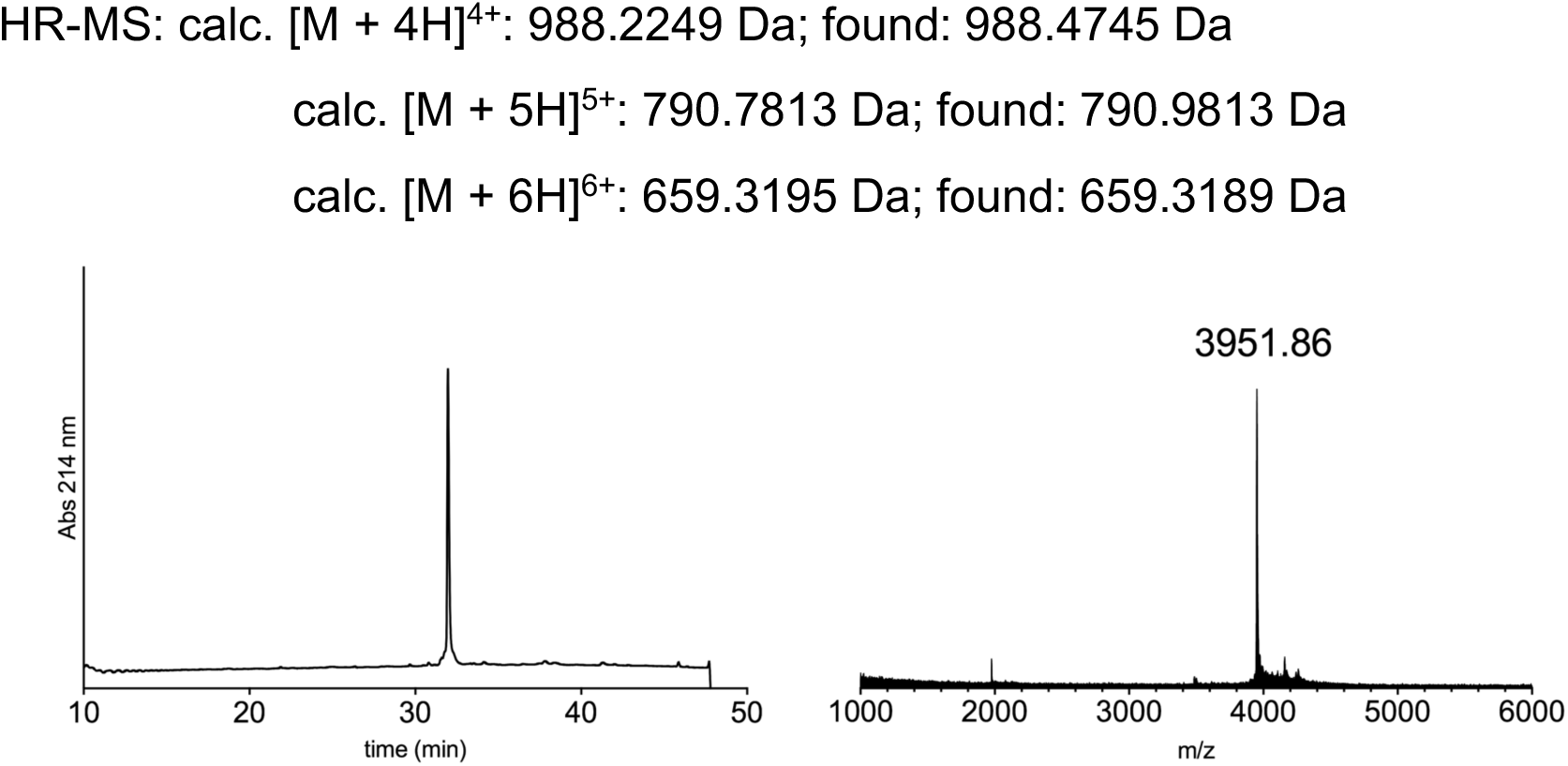

### General procedure for one-pot ligation of peptide fragments_C to N directed ligations

Before performing the ligation, the buffer (guanidinium chloride 6M, Na_2_HPO_4_ 100mM) was sparged with nitrogen. TCEP·HCl was then dissolved in buffer (20 mM), followed by addition of appropriate thiol (40 mM MPAA or 2% v/v TFET^8^, see below for details). The solution was adjusted to pH ∼7 with NaOH 5M and the ion channel peptide fragment (0.57 μmol, 1.0 equiv, 2 mM) was then added, followed by either the Int^N^-B or Int^N^-B_short peptide fragment (0.57 μmol, 1.0 equiv, see details below). The pH was readjusted to ∼7 and incubated at 37 °C. After conversion to the desired ligated product (monitored by HPLC and MALDI-TOF), TCEP·HCl (to reach 40 mM final concentration) and MeONH_2_·HCl (to reach 200 mM final concentration) were dissolved in buffer (15 µL) and added to the ligation mixture, which was then incubated at 37 °C. After conversion to the desired unmasked N-terminal cysteine peptide (monitored by HPLC and MALDI-TOF), the pH was readjusted to ∼7. Activating thiol (40 mM MPAA or 1% v/v TFET, see details below) and the Int^C^-A peptide fragment (0.57 μmol, 1.0 equiv if not stated differently, see below for details) were then added. The pH was adjusted to ∼7 and the reaction was incubated at either 37 °C or at room temperature (see details below). After conversion to the desired ligated product, the ligation mixture was subjected to preparative HPLC purification.

### General procedure for one-pot ligation of peptide fragments_N to C directed ligations

Before performing the ligation, the buffer (guanidinium chloride 6M, Na_2_HPO_4_ 100mM) was sparged with nitrogen. TCEP·HCl was then dissolved in buffer (20 mM) and the pH adjusted to ∼6.4 with NaOH 1M. The ion channel peptide fragment (0.57 μmol, 1.0 equiv, 2 mM) was then added, followed by Int^C^-A_TFET peptide fragment (0.57 μmol, 1.0 equiv). The pH was readjusted to pH ∼6.4 and the reaction mixture was incubated at room temperature. After conversion to the desired ligated product (monitored by HPLC and MALDI-TOF), Int^N^- B_short peptide fragment (0.57 μmol, 1.0 equiv) was added, followed by TFET (1% v/v).

The pH was adjusted to ∼7 and the reaction was incubated at room temperature. After conversion to the desired ligated product, the ligation mixture was subjected to preparative HPLC purification.

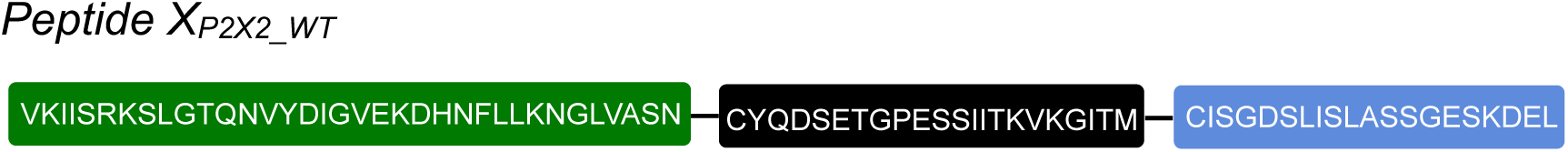

The full peptide was assembled according to the general procedure “C to N directed ligations” described above, using MPAA as the activating thiol. For the second ligation the N-terminal thioester fragment was used in slight excess (1.1 equiv) and the reaction mixture was incubated at 37 °C. Preparative HPLC purification followed by lyophilization yielded the peptide as a fluffy solid (1.7 mg as a TFA salt; yield 32%).

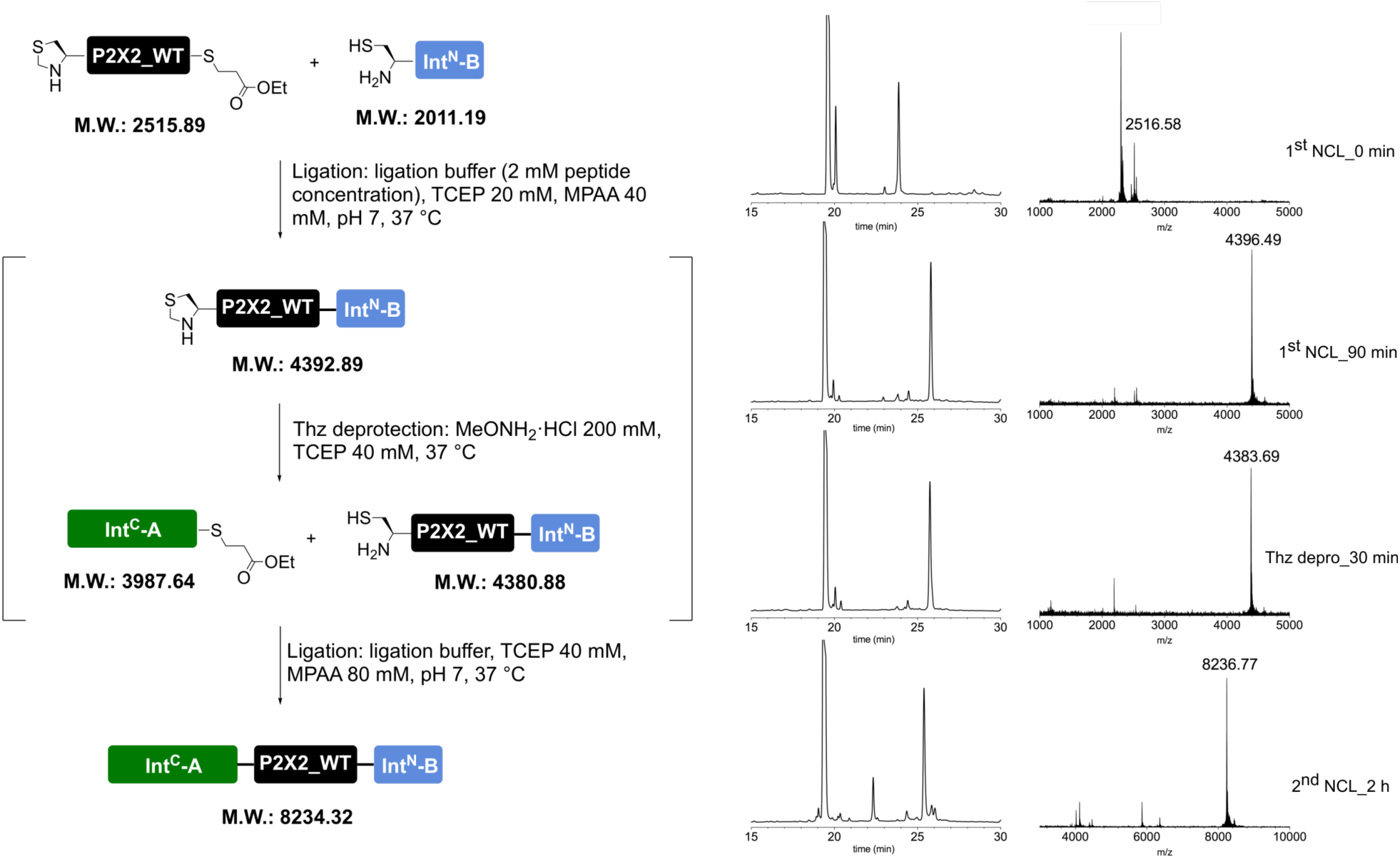

Prep-HPLC purification conditions (C8 column): 0–15% eluent II in eluent I (5 min gradient) followed by 15–45% eluent II in eluent I (45 min gradient).

Low resolution MS (MALDI-TOF): calc. [C_355_H_588_N_95_O_122_S_3_]^+^ [M + H]^+^: 8233.20 Da; found: 8239.33 Da.

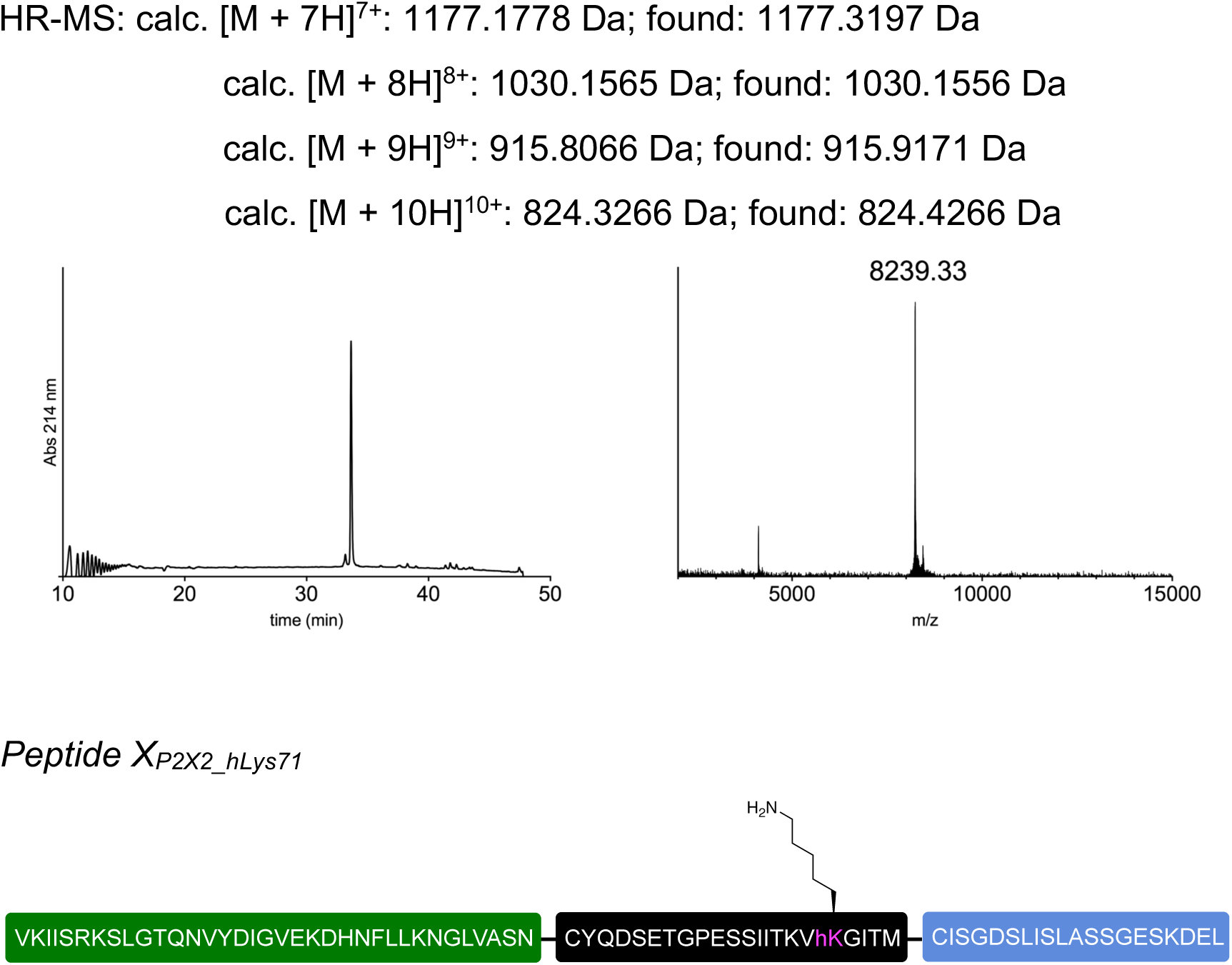

The full peptide was assembled according to the general procedure “C to N directed ligations” described above, using MPAA as the activating thiol. For the second ligation the Int^C^-A fragment was used in slight excess (1.1 equiv) and the reaction mixture was incubated at 37 °C. Preparative HPLC purification followed by lyophilization yielded the peptide as a fluffy solid (1.9 mg as a TFA salt; yield 35%).

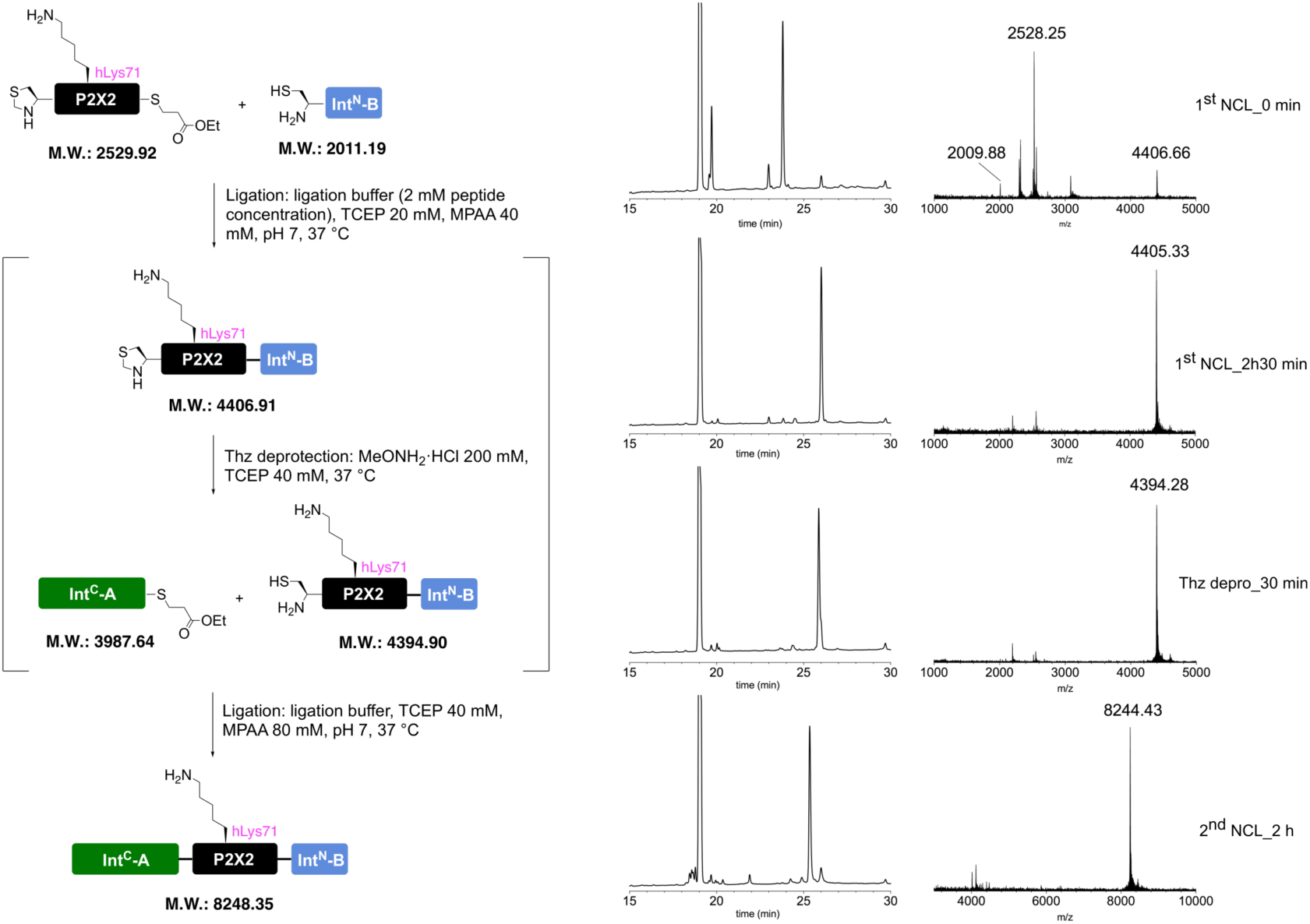

Prep-HPLC purification conditions (C8 column): 0–15% eluent II in eluent I (5 min gradient) followed by 15–45% eluent II in eluent I (45 min gradient).

Low resolution MS (MALDI-TOF): calc. [C_356_H_590_N_95_O_122_S_3_]^+^ [M + H]^+^: 8247.21 Da; found: 8251.23 Da.

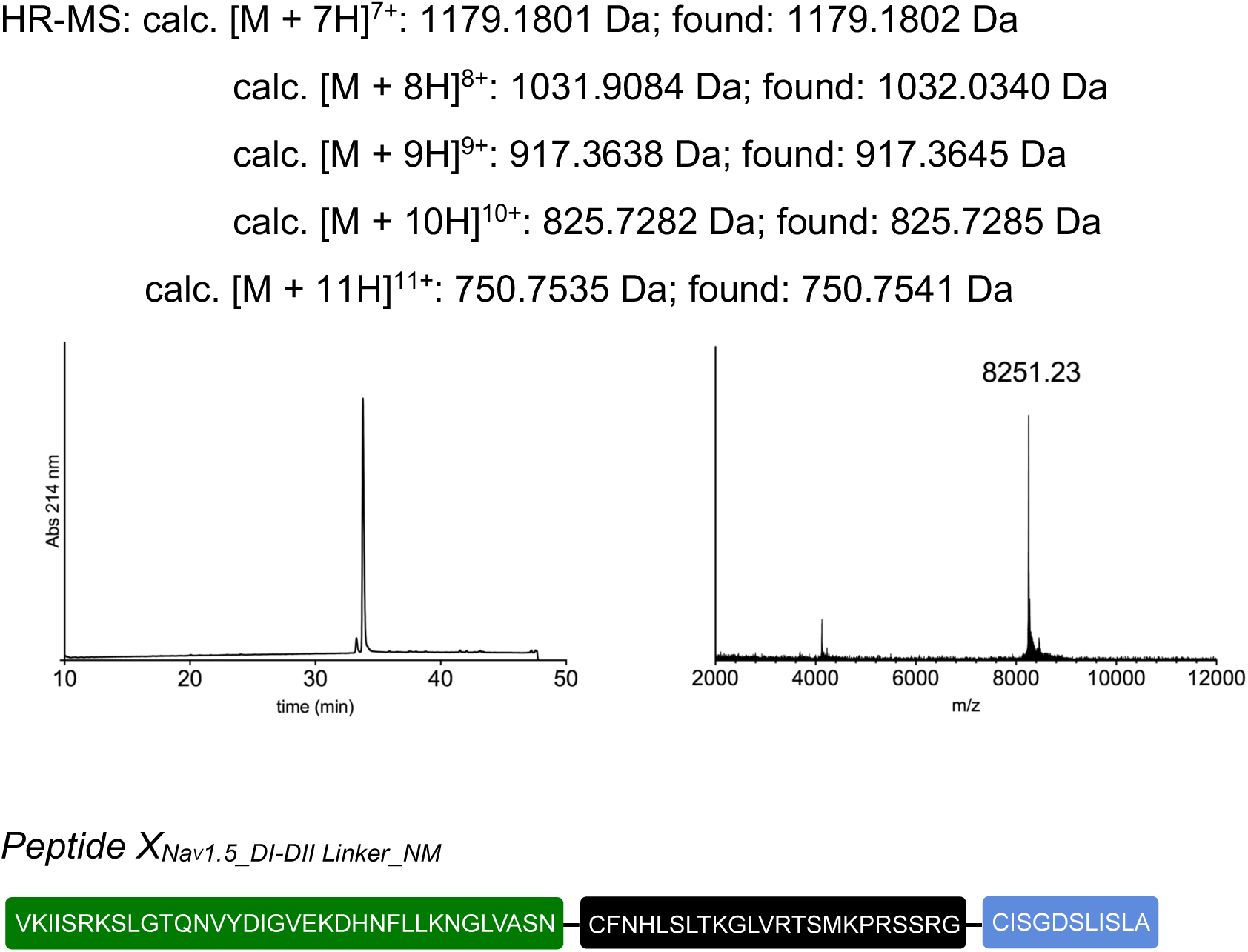

The full peptide was assembled according to the general procedure “C to N directed ligations” described above, using TFET as the activating thiol. For the second ligation, the reaction mixture was incubated at room temperature. Preparative HPLC purification followed by lyophilization yielded the peptide as a fluffy solid (1.8 mg as a TFA salt; yield 35%).

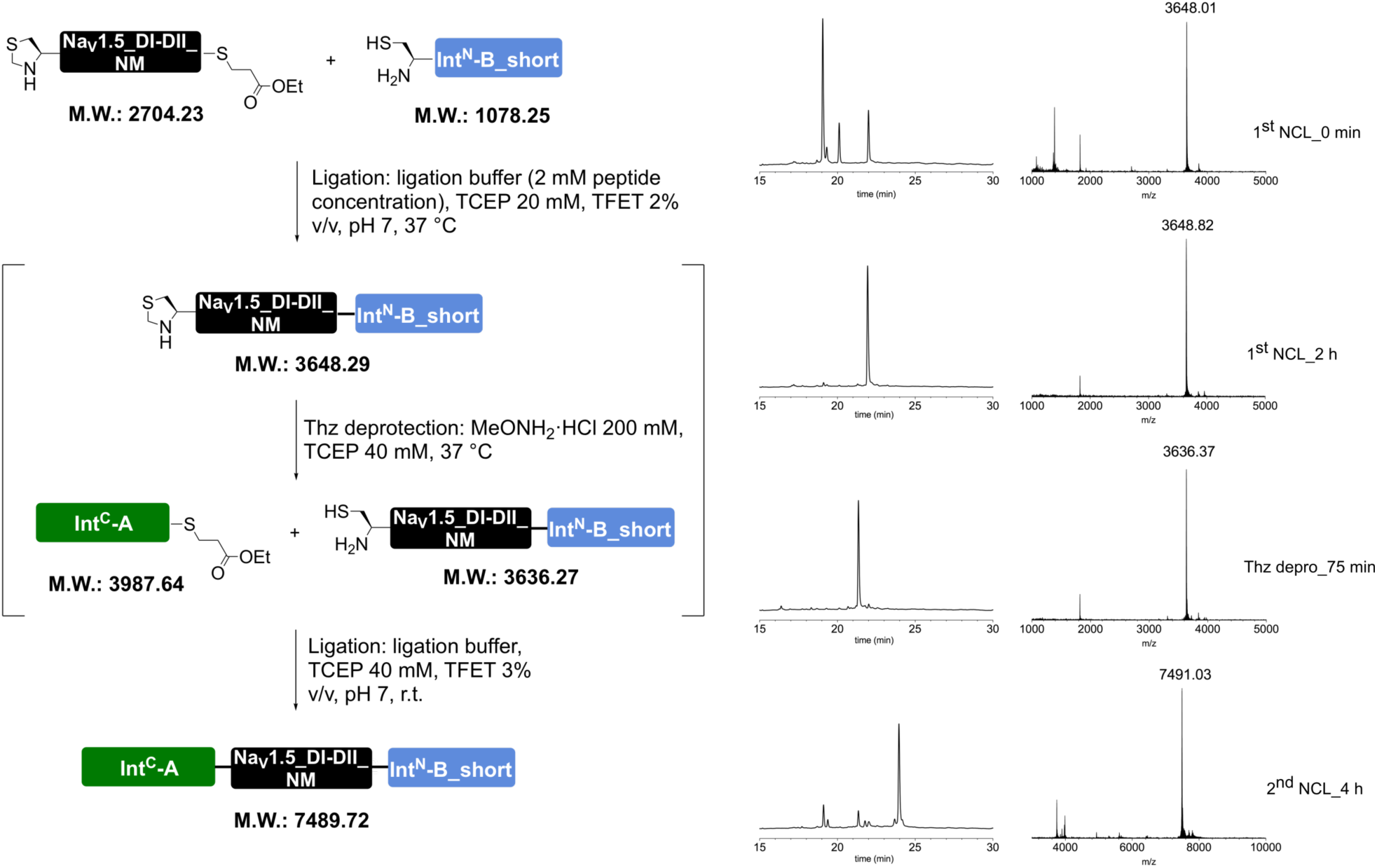

Prep-HPLC purification conditions (C8 column): 0–10% eluent II in eluent I (5 min gradient) followed by 10–45% eluent II in eluent I (45 min gradient).

Low resolution MS (MALDI-TOF): calc. [C_326_H_548_N_97_O_98_S_3_]^+^ [M + H]^+^: 7489.01 Da; found: 7487.79 Da.

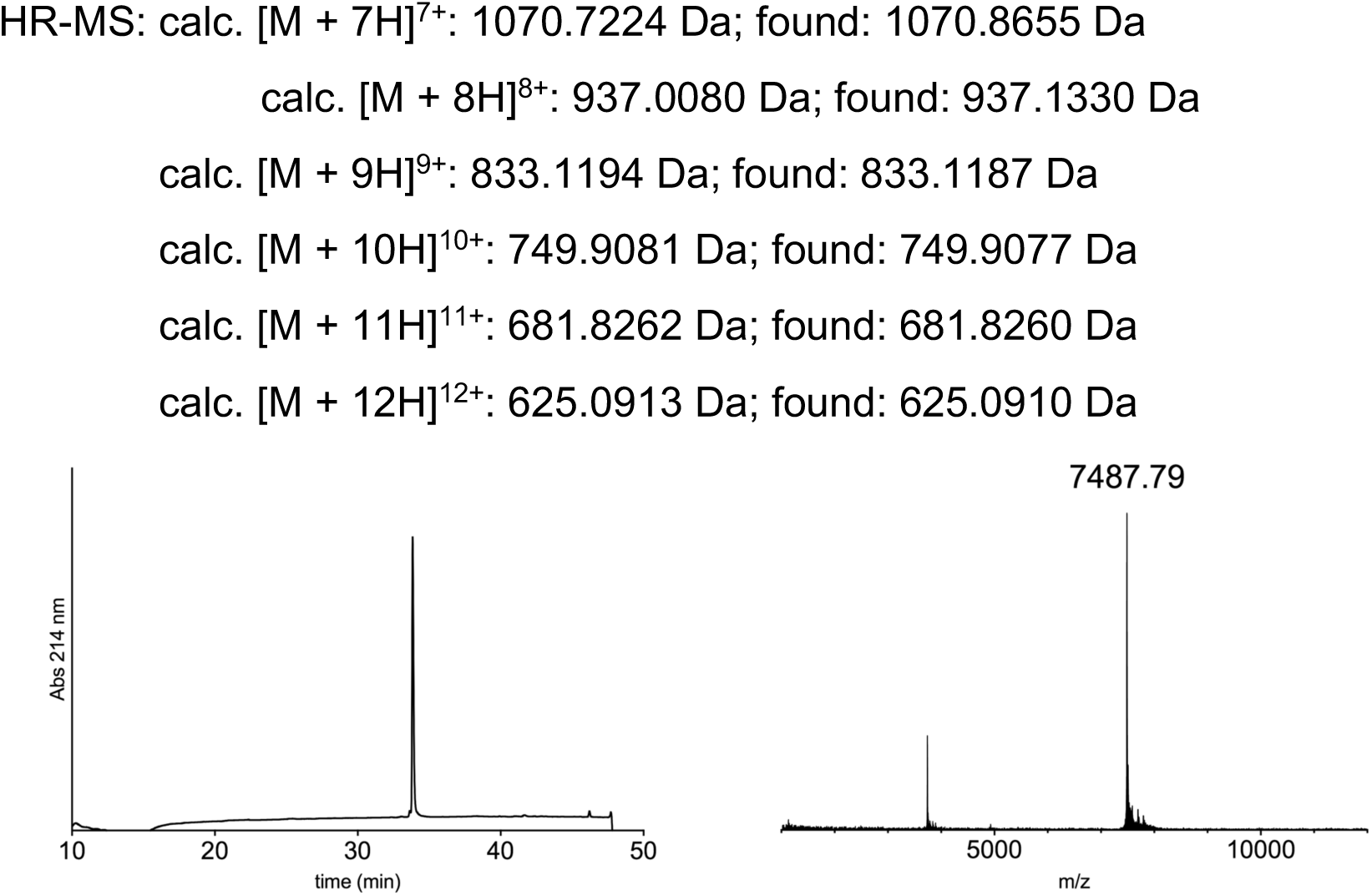

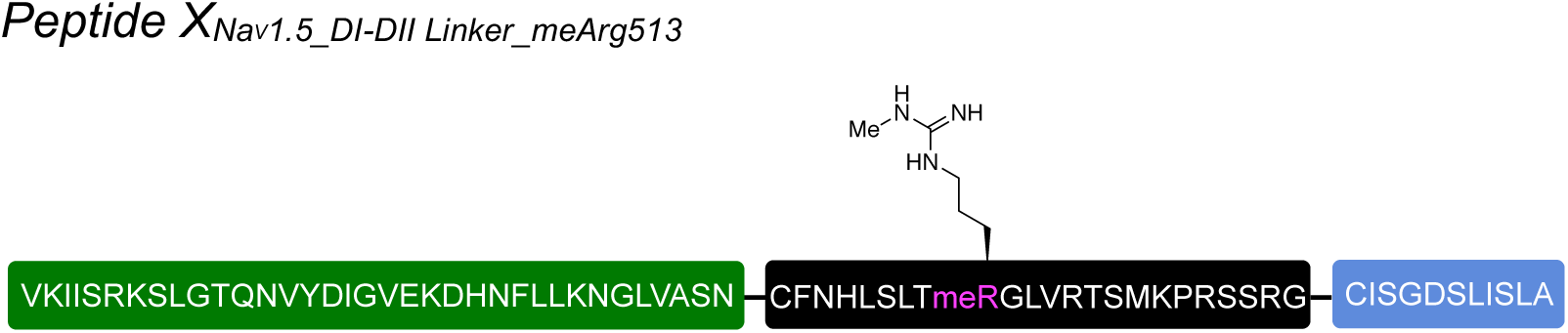

The full peptide was assembled according to the general procedure “C to N directed ligations” described above, using MPAA as the activating thiol. For the second ligation, the reaction mixture was incubated at room temperature. Since the desired product co-eluted with MPAA during preparative HPLC purification, the isolated fraction containing the peptide was dialyzed in multiple steps (2h + 2h + over night at 5 °C) in water using a cellulose membrane with a cutoff of 2 kDa. The dialyzed fraction was then diluted with eluent I and lyophilized to obtain the peptide as a fluffy solid (1.0 mg as a TFA salt; yield 19%).

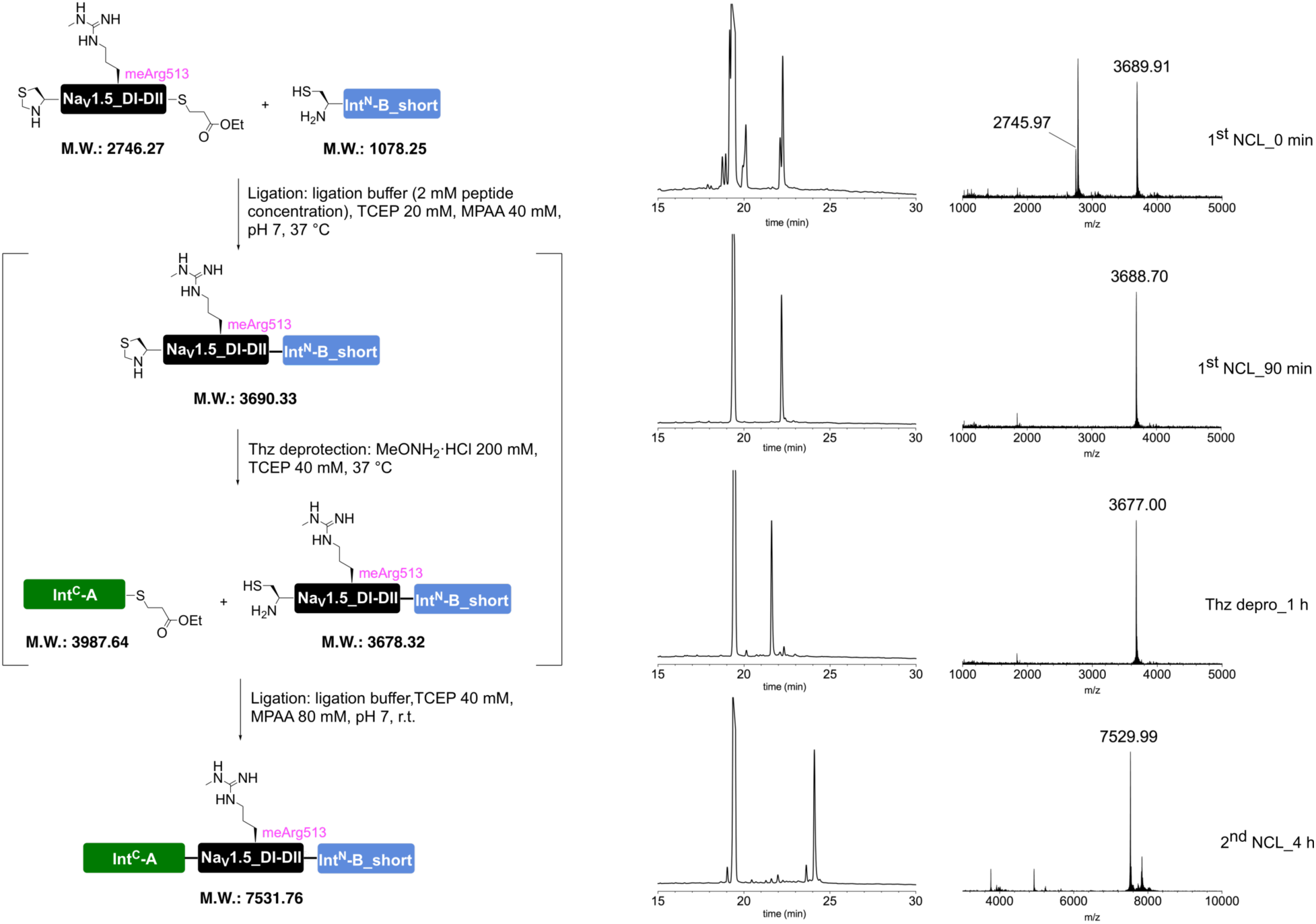

Prep-HPLC purification conditions (C8 column): 0–10% eluent II in eluent I (5 min gradient) followed by 10–45% eluent II in eluent I (45 min gradient).

Low resolution MS (MALDI-TOF): calc. [C_327_H_550_N_99_O_98_S_3_]^+^ [M + H]^+^: 7531.03 Da; found: 7529.99 Da.

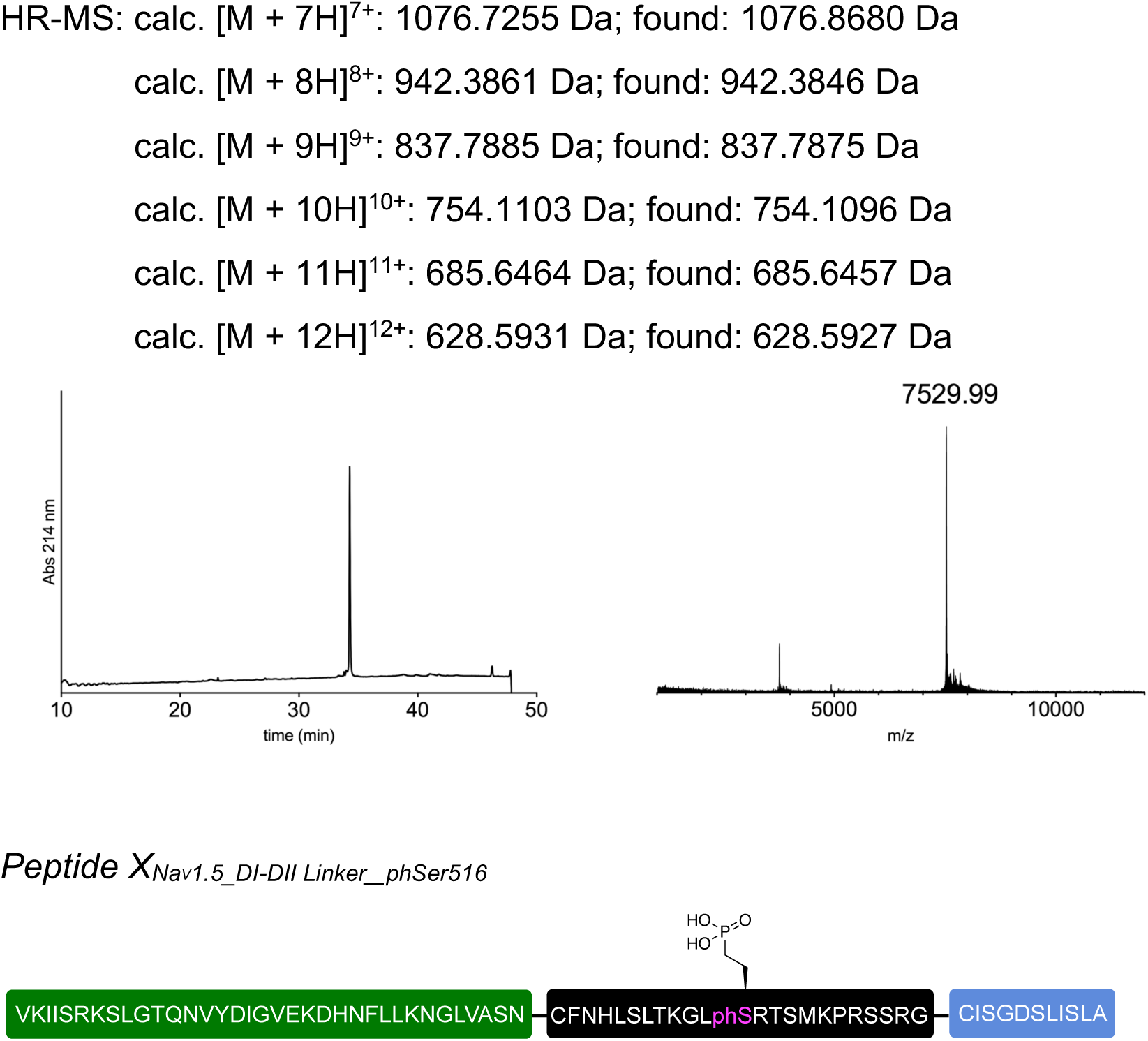

The full peptide was assembled according to the general procedure “C to N directed ligations” described above, using TFET as the activating thiol. For the second ligation, the reaction mixture was incubated at room temperature. Preparative HPLC purification followed by lyophilization yielded the peptide as a fluffy solid (1.7 mg as a TFA salt; yield 33%).

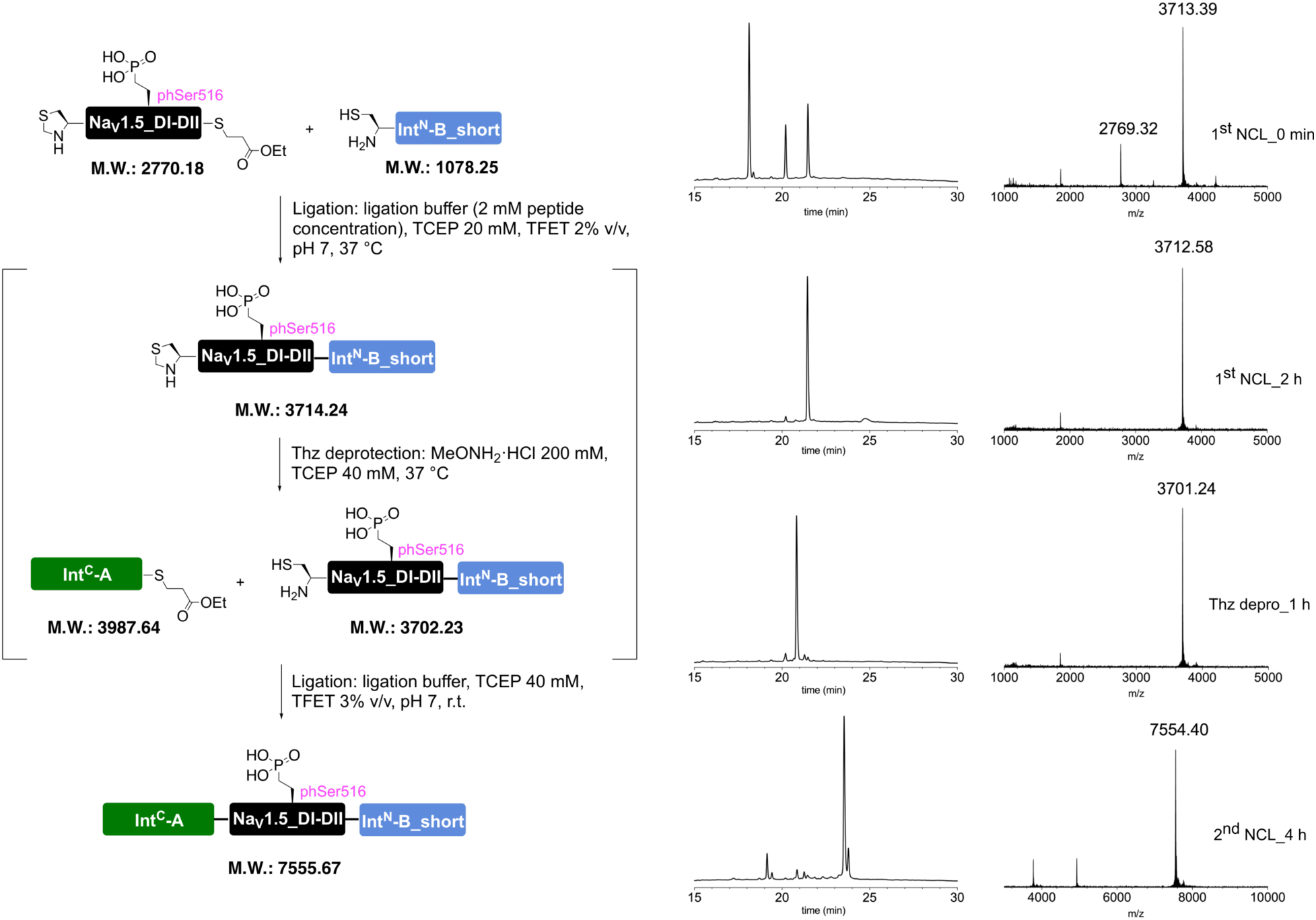

Prep-HPLC purification conditions (C8 column): 0–10% eluent II in eluent I (5 min gradient) followed by 10–45% eluent II in eluent I (45 min gradient).

Low resolution MS (MALDI-TOF): calc. [C_325_H_547_N_97_O_101_PS_3_]^+^ [M + H]^+^: 7554.96 Da; found: 7553.58 Da.

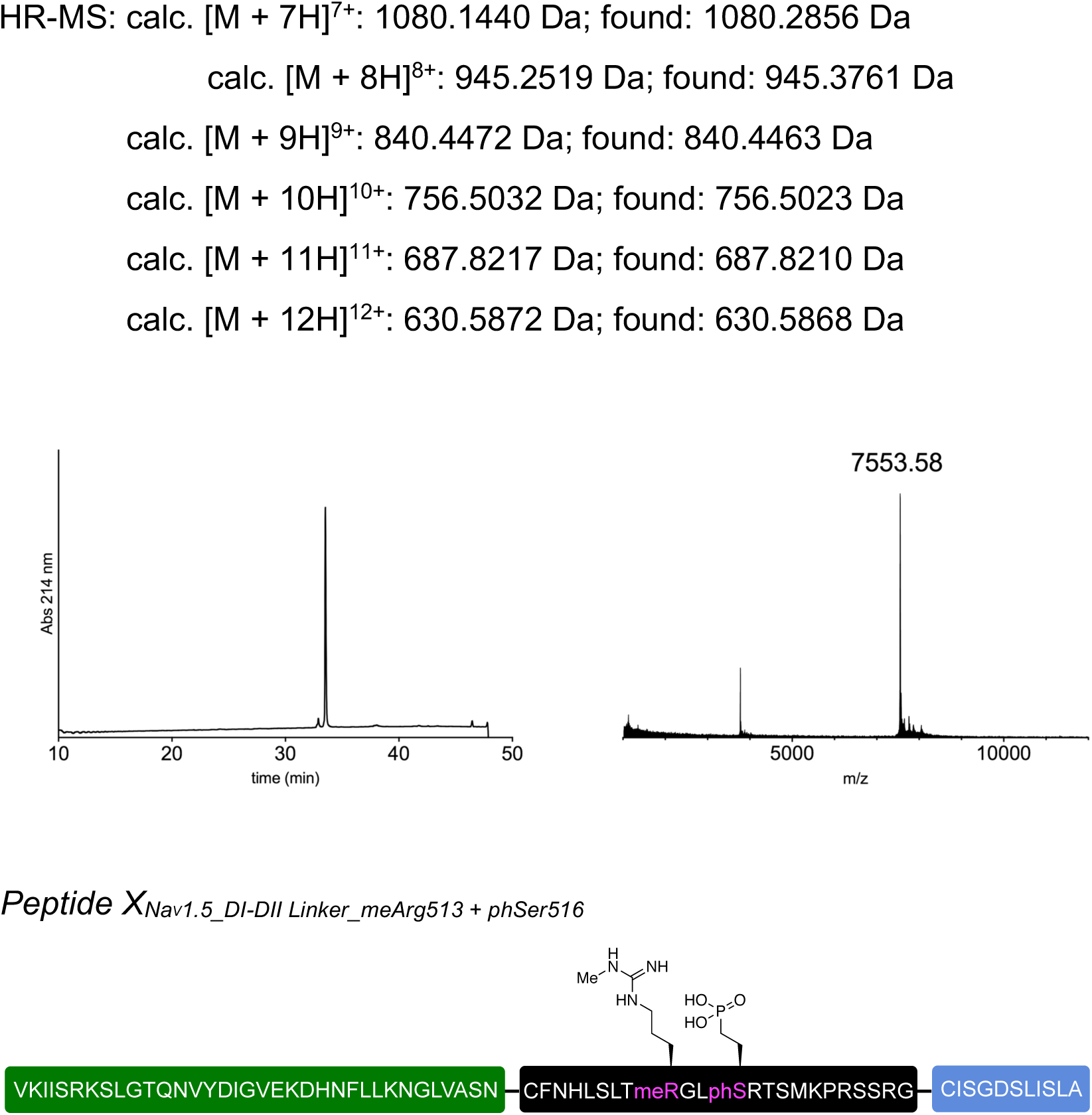

The full peptide was assembled according to the general procedure “C to N directed ligations” described above, using TFET as the activating thiol. For the second ligation, the reaction mixture was incubated at room temperature. Preparative HPLC purification followed by lyophilization yielded the peptide as a fluffy solid (1.7 mg as a TFA salt; yield 33%).

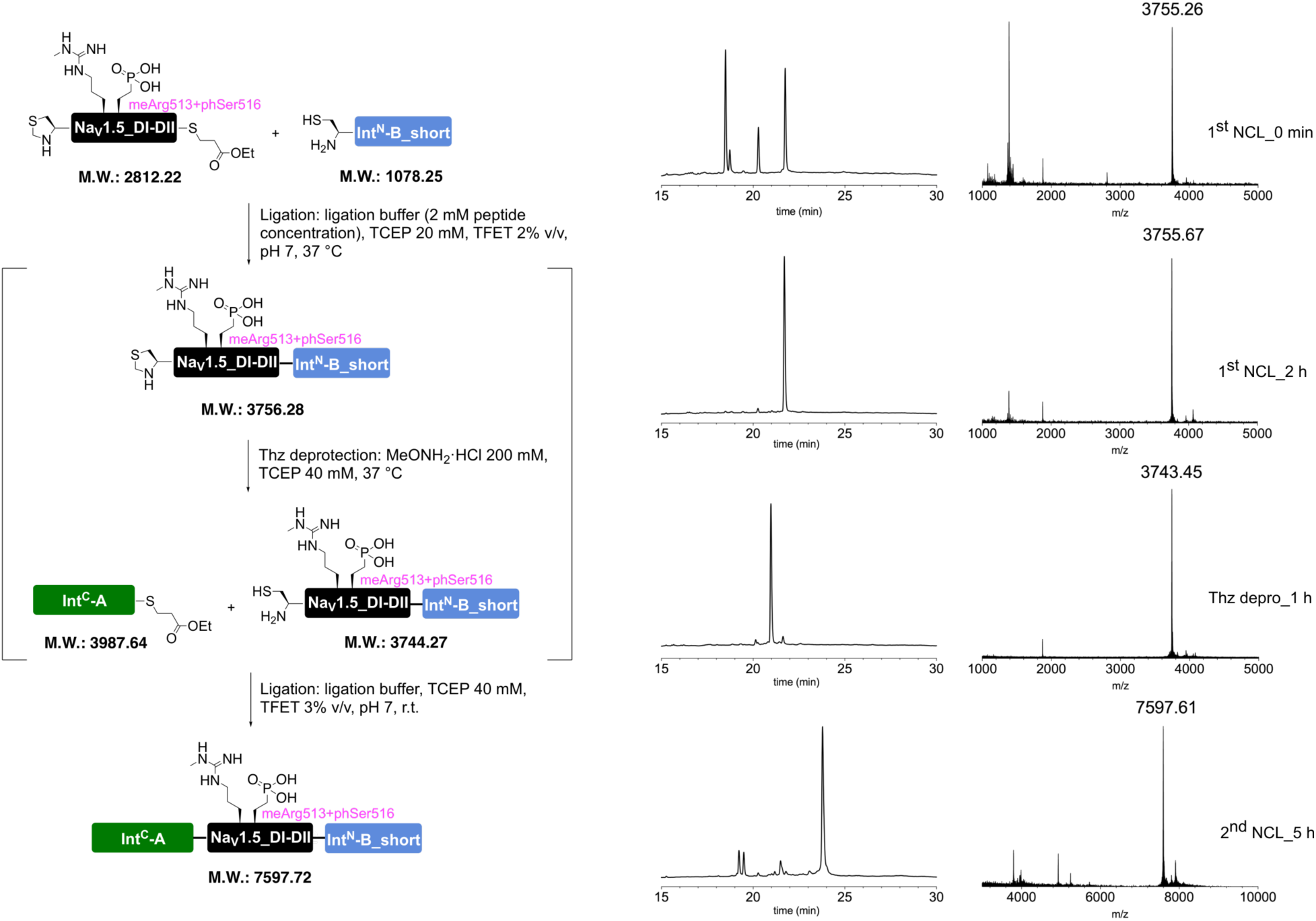

Prep-HPLC purification conditions (C8 column): 0–10% eluent II in eluent I (5 min gradient) followed by 10–45% eluent II in eluent I (45 min gradient).

Low resolution MS (MALDI-TOF): calc. [C_326_H_549_N_99_O_101_PS_3_]^+^ [M + H]^+^: 7596.99 Da; found: 7595.97 Da.

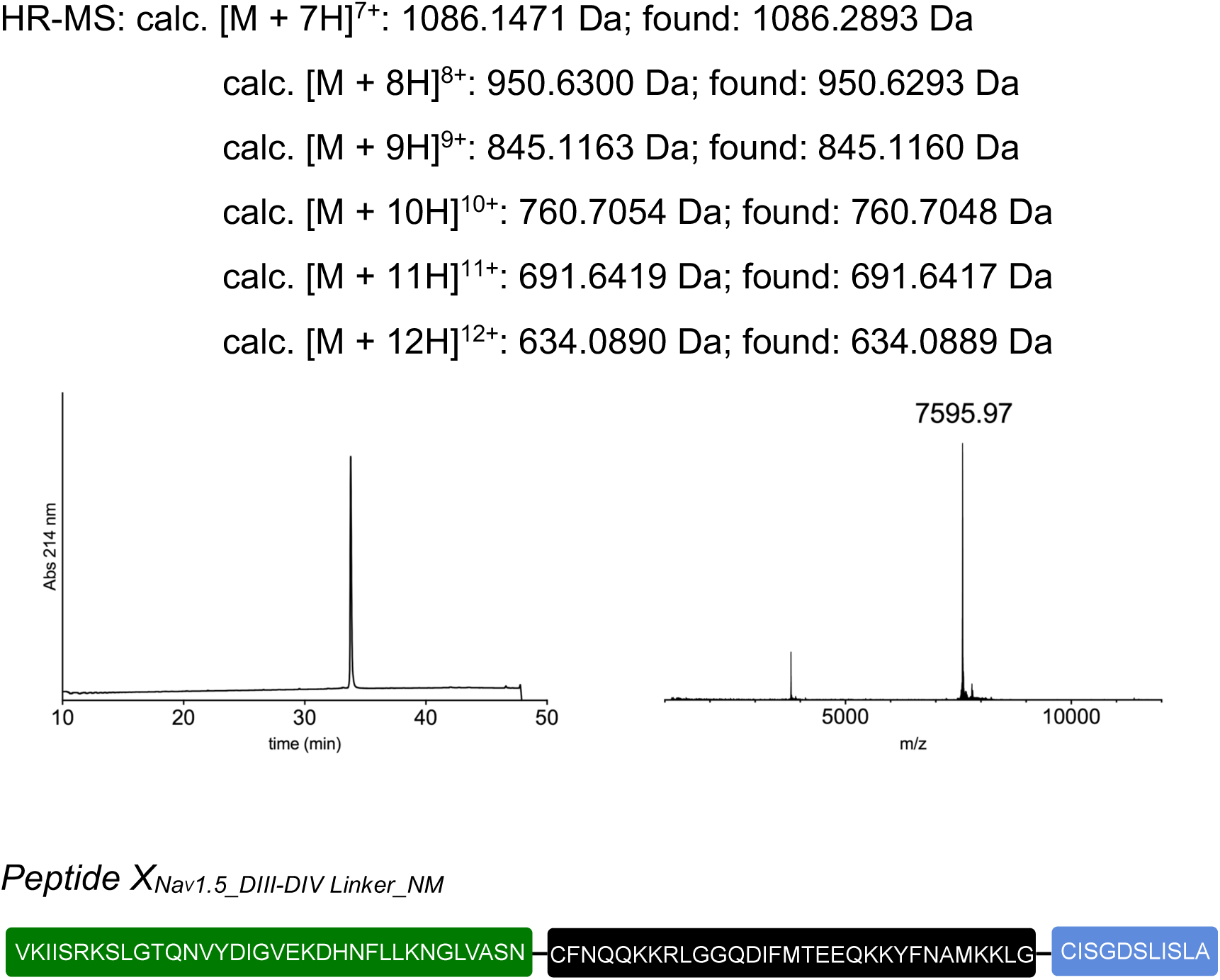

The full peptide was assembled according to the general procedure “C to N directed ligations” described above, using TFET as the activating thiol. For the second ligation, the reaction mixture was incubated at room temperature. Preparative HPLC purification followed by lyophilization yielded the peptide as a fluffy solid (0.9 mg as a TFA salt; yield 15%).

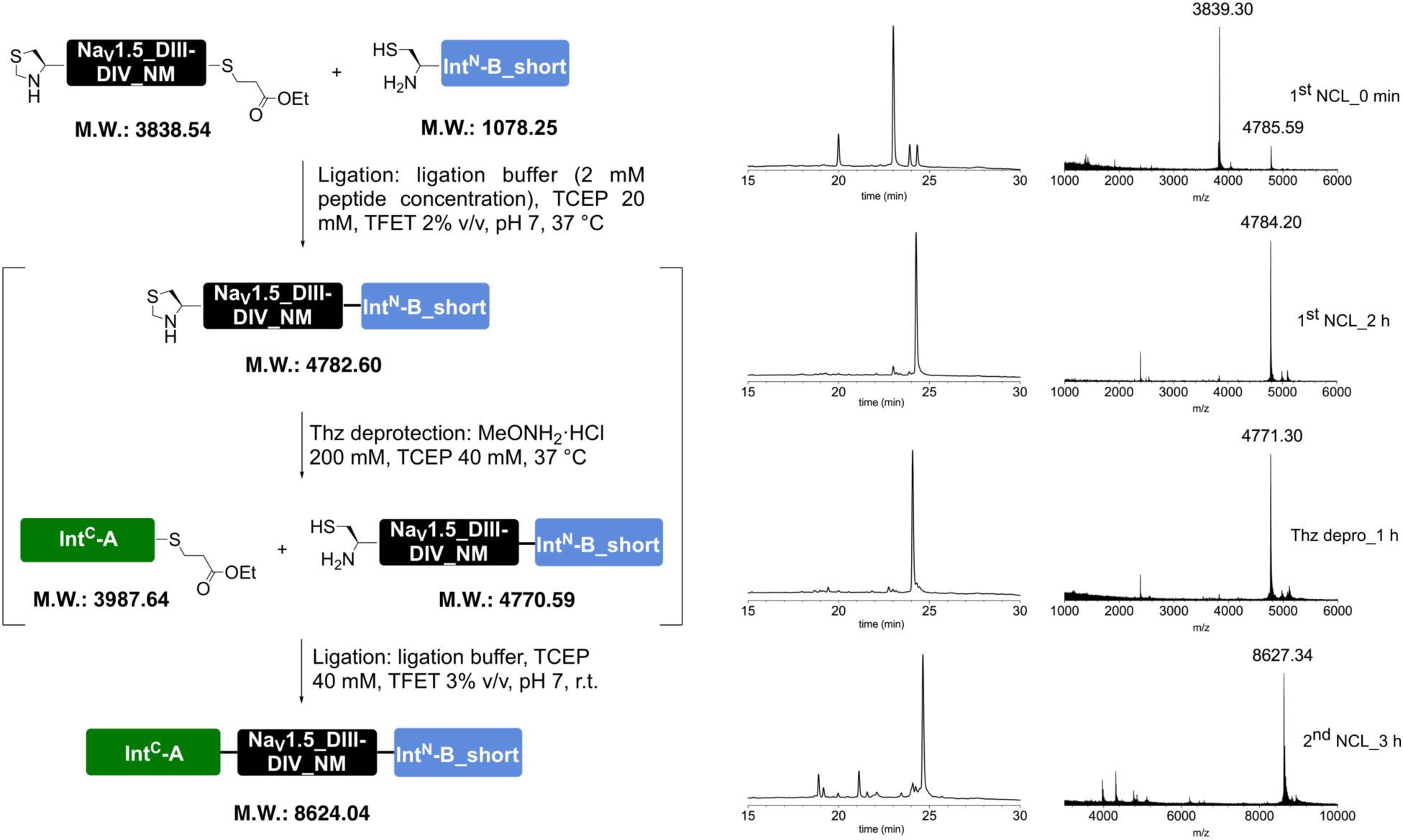

Prep-HPLC purification conditions (C8 column): 0–10% eluent II in eluent I (5 min gradient) followed by 10–45% eluent II in eluent I (45 min gradient).

Low resolution MS (MALDI-TOF): calc. [C_381_H_623_N_106_O_113_S_4_]^+^ [M + H]^+^: 8623.53 Da; found: 8622.99 Da.

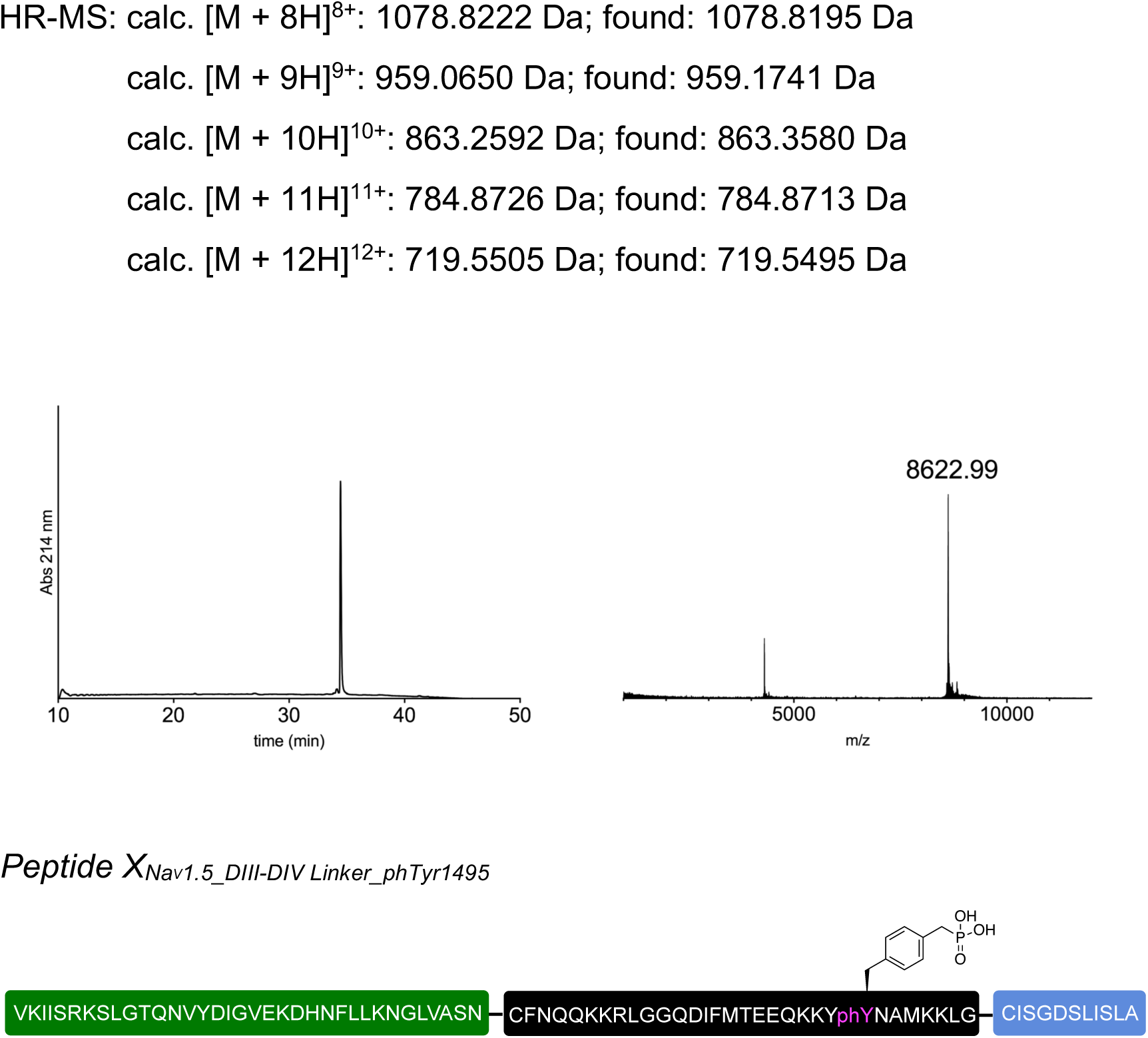

The full peptide was assembled according to the general procedure “C to N directed ligations” described above, using TFET as the activating thiol. For the second ligation, the reaction mixture was incubated at room temperature. Preparative HPLC purification followed by lyophilization yielded the peptide as a fluffy solid (2.0 mg as a TFA salt; yield 34%).

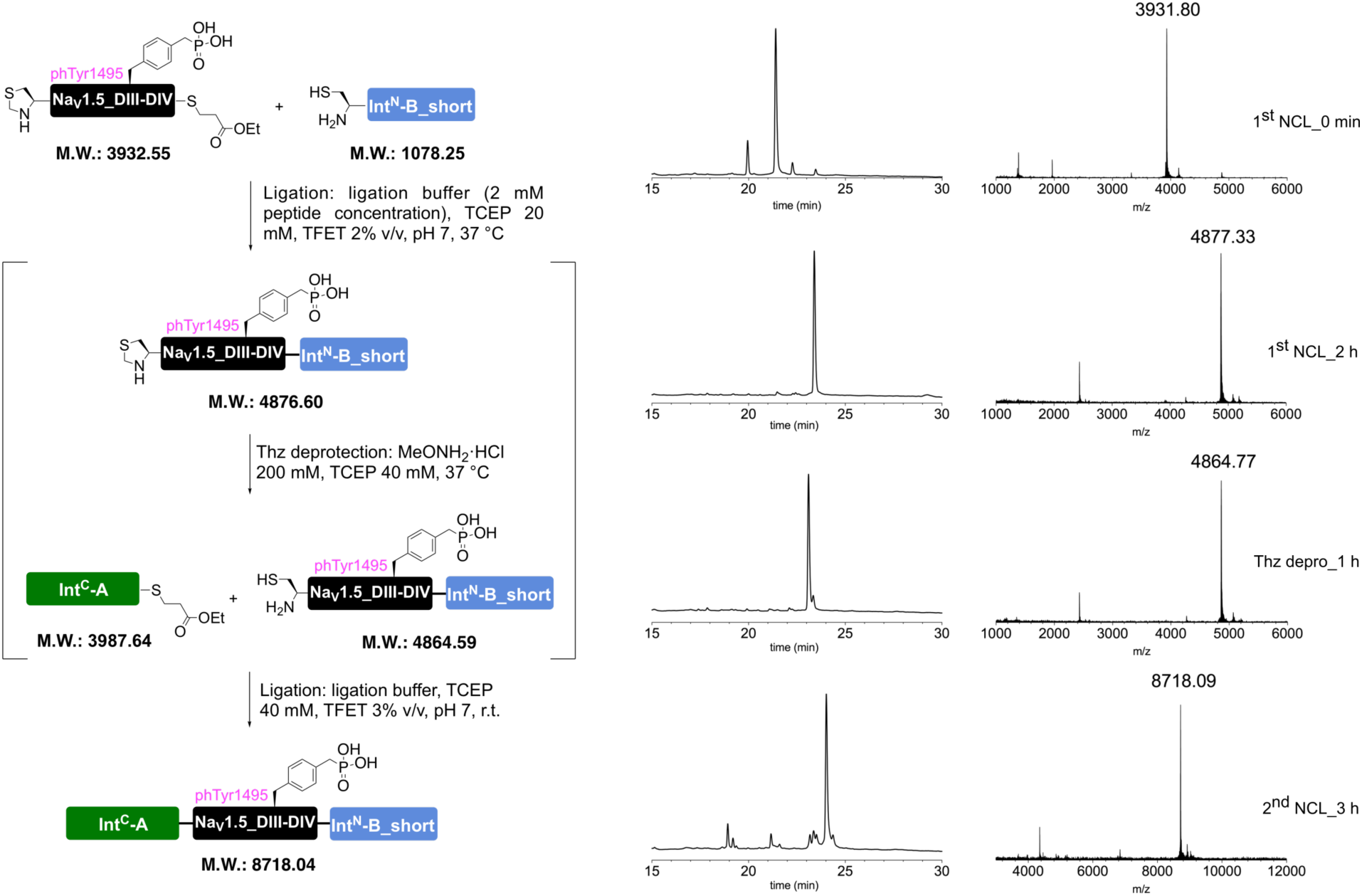

Prep-HPLC purification conditions (C8 column): 0–10% eluent II in eluent I (5 min gradient) followed by 10–45% eluent II in eluent I (45 min gradient).

Low resolution MS (MALDI-TOF): calc. [C_382_H_626_N_106_O_116_PS_4_]^+^ [M + H]^+^: 8717.51 Da; found: 8717.22 Da.

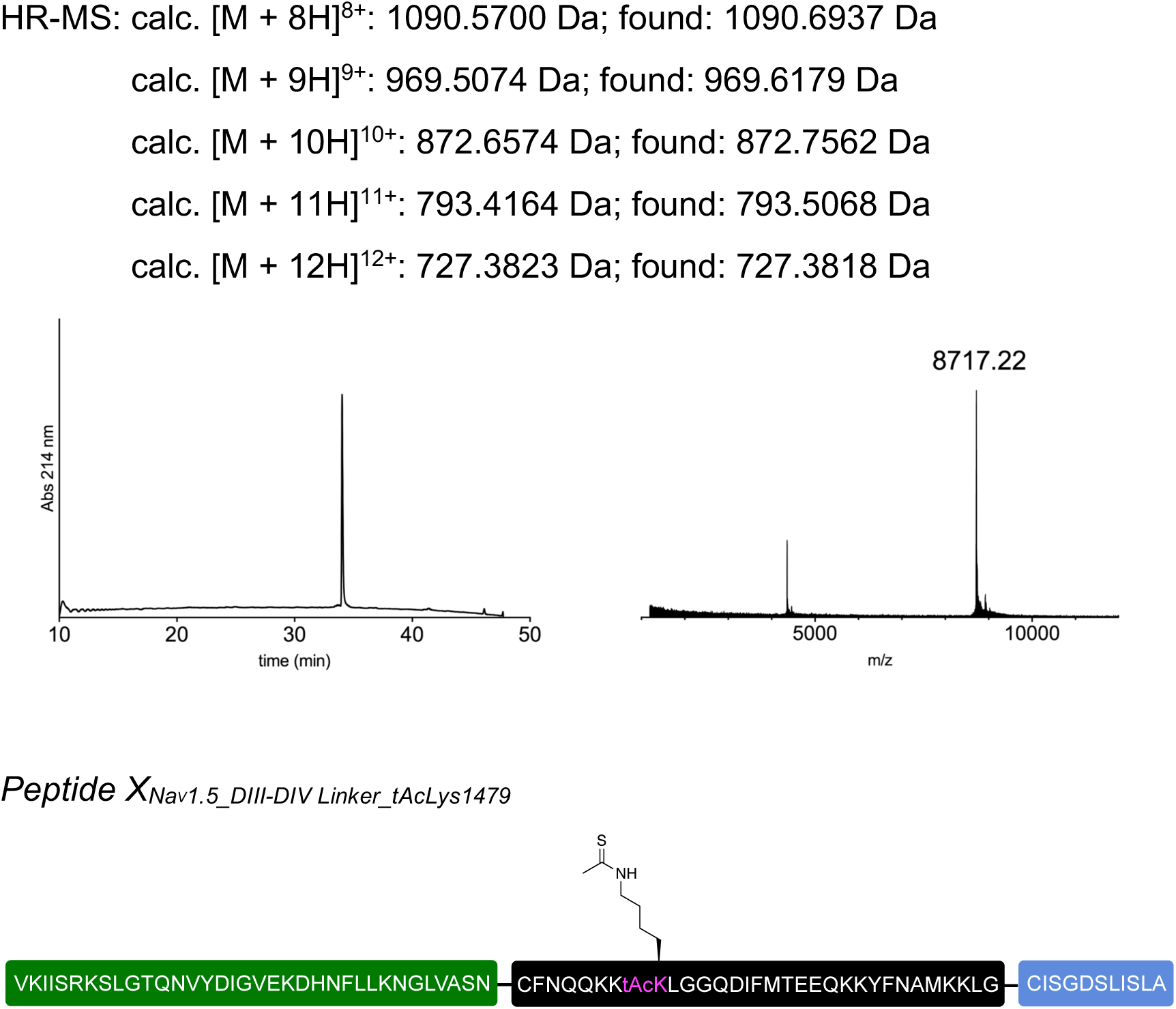

The full peptide was assembled according to the general procedure “N to C directed ligations” described above. For the second ligation, the reaction mixture was incubated at room temperature. Preparative HPLC purification followed by lyophilization yielded the peptide as a fluffy solid (1.2 mg as a TFA salt; yield 20%).

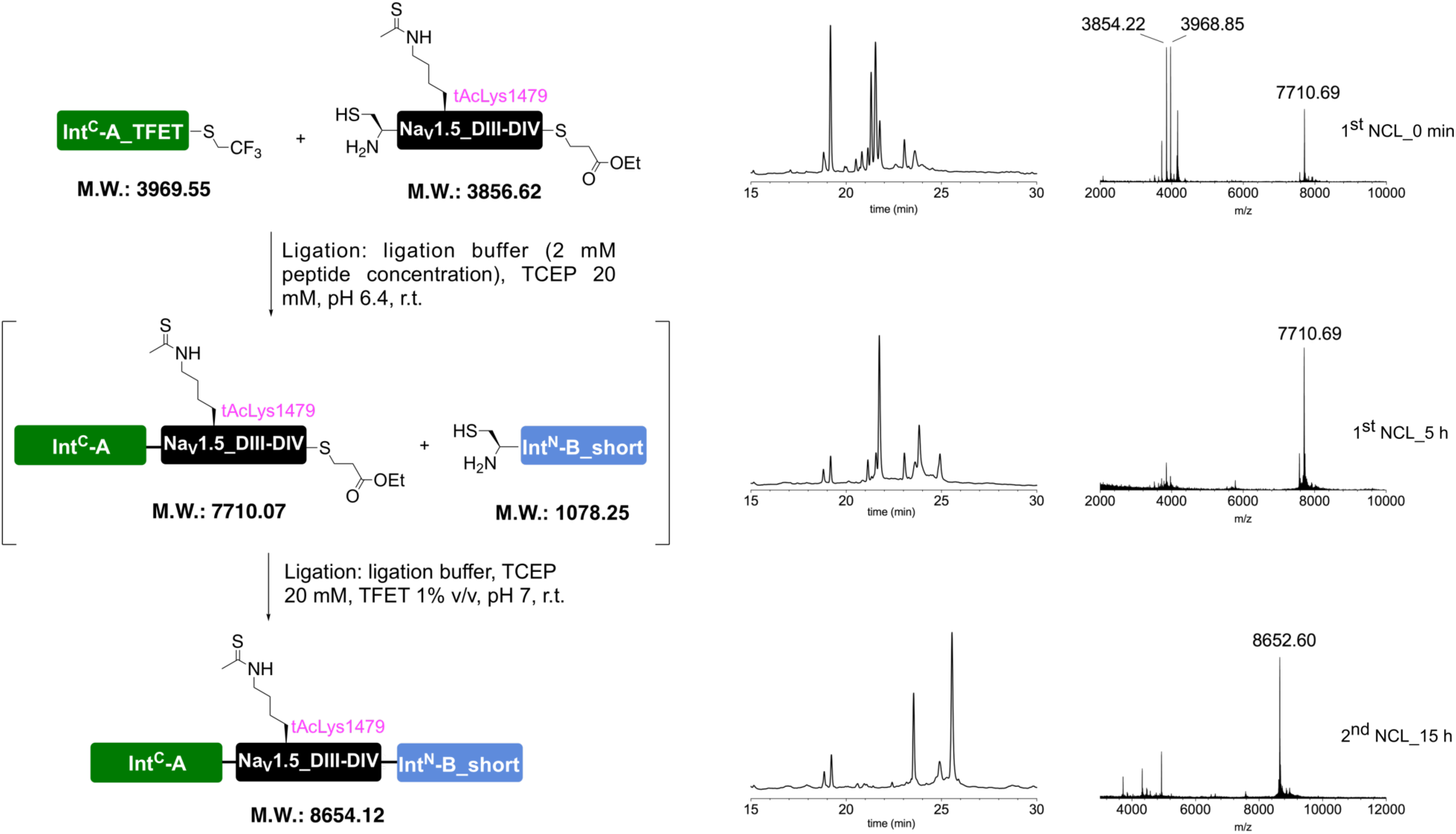

Prep-HPLC purification conditions (C8 column): 0–10% eluent II in eluent I (5 min gradient) followed by 10–45% eluent II in eluent I (45 min gradient).

Low resolution MS (MALDI-TOF): calc. [C_383_H_625_N_104_O_113_S_5_]^+^ [M + H]^+^: 8653.51 Da; found: 8653.47 Da.

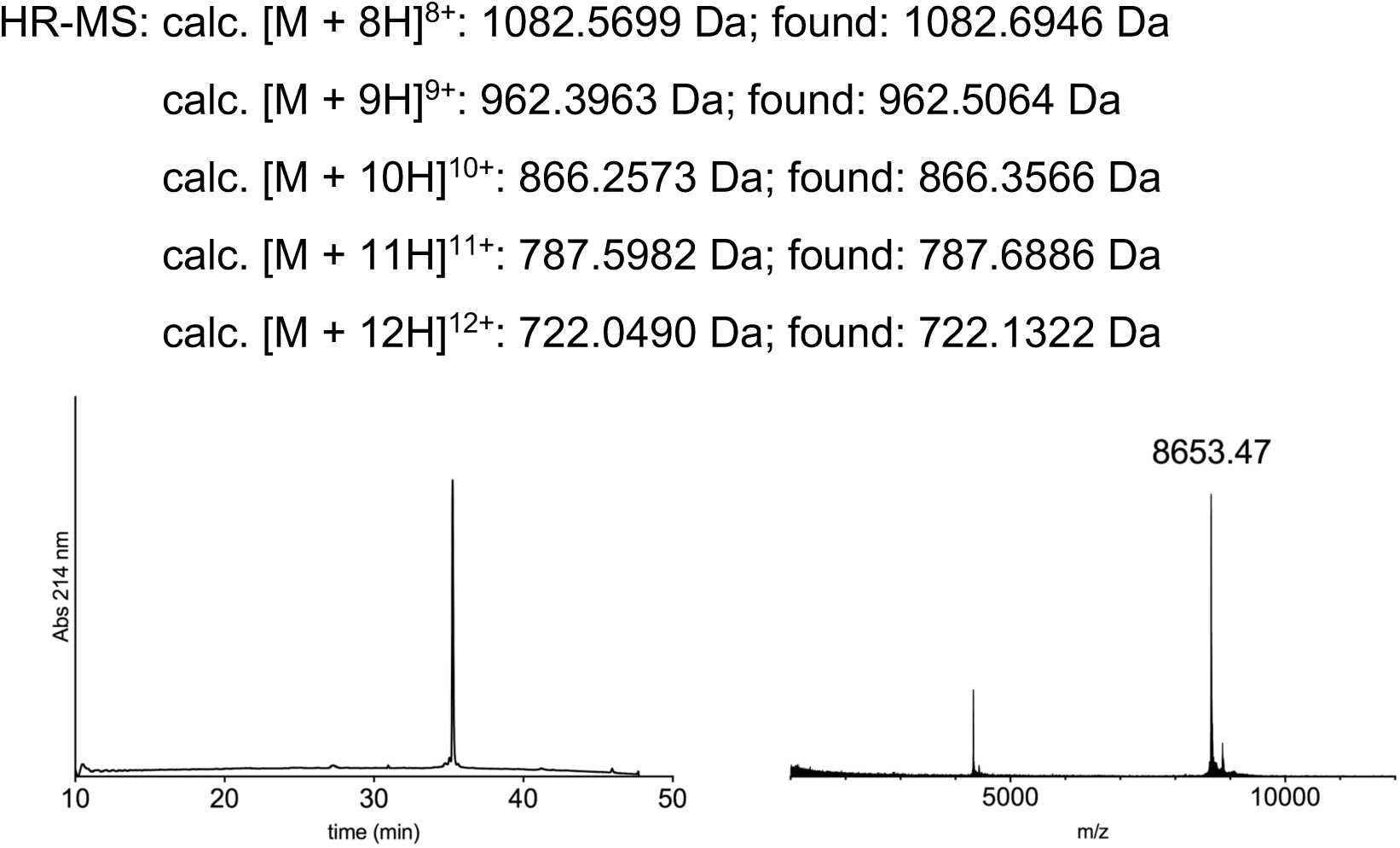

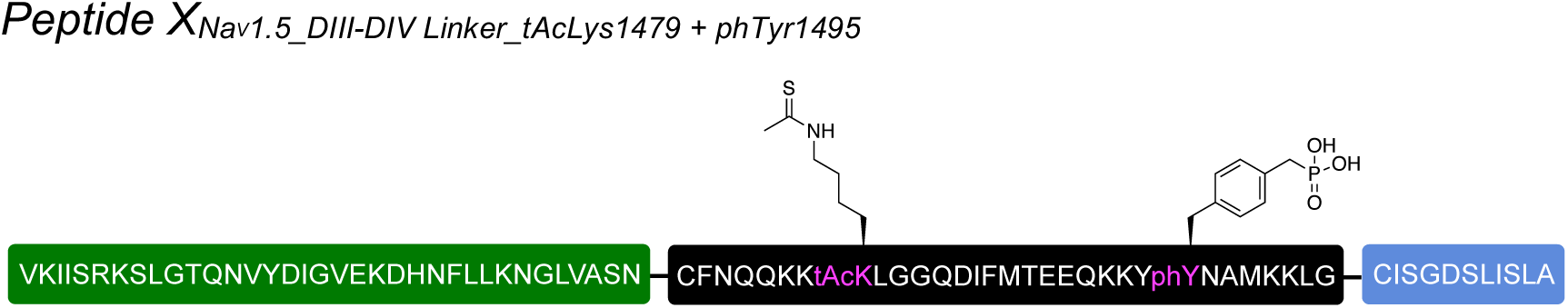

The full peptide was assembled according to the general procedure “N to C directed ligations” described above. Preparative HPLC purification followed by lyophilization yielded the peptide as a fluffy solid (1.2 mg as a TFA salt; yield 20%).

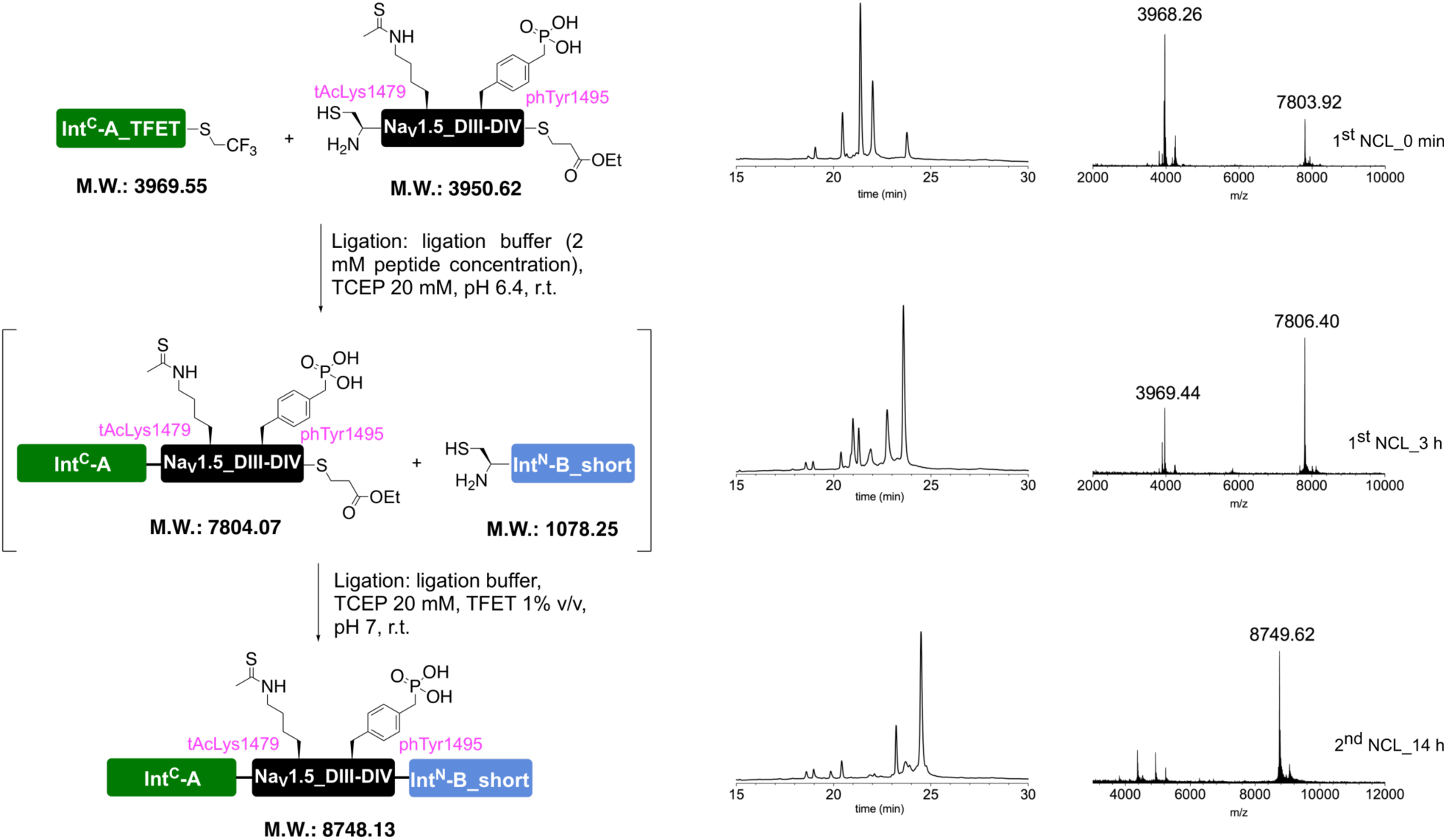

Prep-HPLC purification conditions (C8 column): 0–10% eluent II in eluent I (5 min gradient) followed by 10–45% eluent II in eluent I (45 min gradient).

Low resolution MS (MALDI-TOF): calc. [C_384_H_628_N_104_O_116_PS_5_]^+^ [M + H]^+^: 8747.49 Da; found: 8745.23 Da.

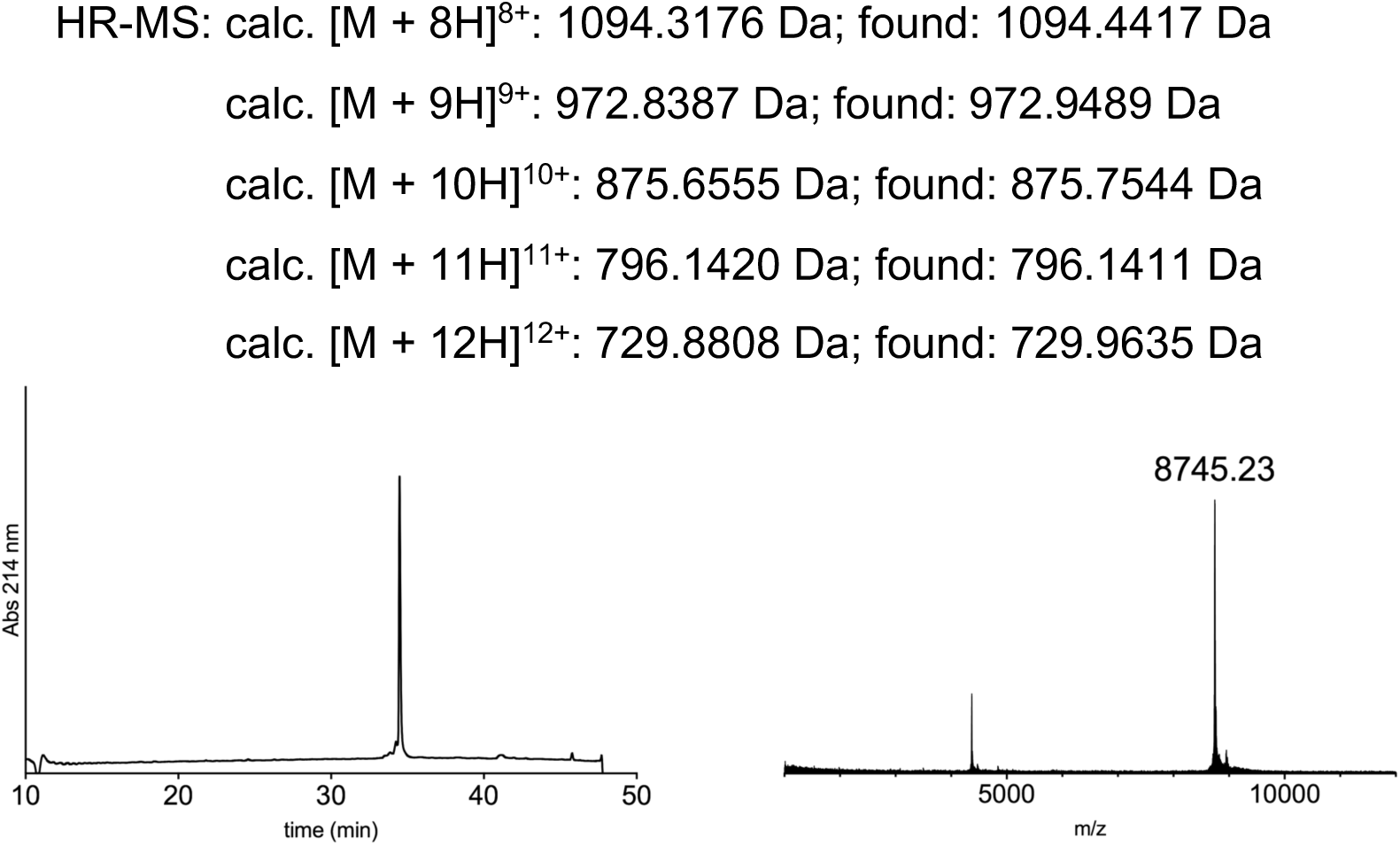

## Notes

#### Summary of Updates

Revised discussion and additional data on reconstitution of GFP using tPTS.

